# Towards understanding of allostery in MALT1: a possible role of interdomain motions as revealed by NMR and AlphaFold

**DOI:** 10.1101/2024.02.01.578365

**Authors:** Johan Wallerstein, Xiao Han, Maria Levkovets, Dmitry Lesovoy, Daniel Malmodin, Claudio Mirabello, Björn Wallner, Renhua Sun, Tatyana Sandalova, Peter Agback, Göran Karlsson, Adnane Achour, Tatiana Agback, Vladislav Orekhov

## Abstract

Mucosa-associated lymphoid tissue lymphoma-translocation protein 1 (MALT1) has emerged as an attractive target for the development of modulatory compounds, particularly in the treatment of lymphoma and other cancers. While the three-dimensional structure of MALT1(PCASP-Ig3)_339–719_ has been previously determined through X-ray analysis, its dynamic behaviour in solution has remained largely unexplored. We present here inaugural dynamic analyses of the apo MALT1(PCASP-Ig3)_339–719_ form along with its mutated variant, E549A. This investigation harnessed an array of NMR relaxation techniques, including longitudinal and transverse ^15^N auto-relaxation, heteronuclear NOE, transverse cross-correlated relaxation and NOE measurements between side-chain methyl groups. Our findings unequivocally confirm that MALT1(PCASP-Ig3)_339–719_ exists solely as a monomer in solution, and demonstrate that the two domains display semi-independent movements in relation to each other. Our extensive dynamic study, covering a range of time scales, along with the assessment of diverse conformational populations for MALT1(PCASP-Ig3)_339–719_, by Molecular Dynamic simulations, Alpha Fold modelling and PCA analysis, shed light at potential mechanisms underlying the allosteric regulation of this enzyme, and the specific importance of interdomain motions.

## 1.0 INTRODUCTION

The human mucosa-associated lymphoid tissue lymphoma-translocation protein 1 (MALT1) is a unique human paracaspase recognized for its distinct cleaving capacity and substrate specificity (Hamp *et al*., 2021). Crucial for the survival, proliferation and functions of B and T cells in adaptive immune responses, MALT1 activates the NF-κB signalling pathway following antigen stimulation (Ruland *et al*., 2003, Ruefli-Brasse *et al*., 2004, Thome, 2008). Dysregulated MALT1 activity is implicated in various lymphoid malignancies, leukaemia (Hailfinger *et al*., 2009, Davis *et al*., 2010) and several other cancers including glioblastoma, melanoma, and breast cancer (Wang *et al*., 2017, Ekambaram *et al*., 2018, Jacobs *et al*., 2020), as well as a large array of autoimmune diseases (Jaworski *et al*., 2014, Gewies *et al*., 2014, Howes *et al*., 2016, Afonina *et al*., 2016). As a key regulator of adaptive immune responses, MALT1 represents a clear target for developing modulatory/inhibitory compounds to treat lymphoma, cancers, and autoimmune diseases.

By forming filament CBM complexes alongside BCL10 and CARD11 (Ruland & Hartjes, 2019, Qiao *et al*., 2013, Li & Wang, 2018), MALT1 serves both as a scaffolding template and a protease, recruiting signalling factors and cleaving substrate proteins to regulate *e.g.* T cell activation (Jaworski & Thome, 2016, Rebeaud *et al*., 2008, Coornaert *et al*., 2008). MALT1 exhibits a complex multi-domain structure comprising an N-terminal death domain (DD), followed by the two immunoglobulin domains Ig1 and Ig2, the paracaspase domain (PCASP), and a C-terminal Ig3 domain (**Figure 1a**). Recent cryo-EM analyses (Schlauderer *et al*., 2018) unveiled that BCL10 forms filaments adorned by MALT1, with specific emphasis on the incorporation of the DD domain. However, the structural flexibility of the remaining MALT1 protein precluded its visualisation, leaving the exact conformation ensembles of these parts of the formed complexes unknown (**Figure 1b**) (Schlauderer *et al*., 2018).

**Figure 1.**
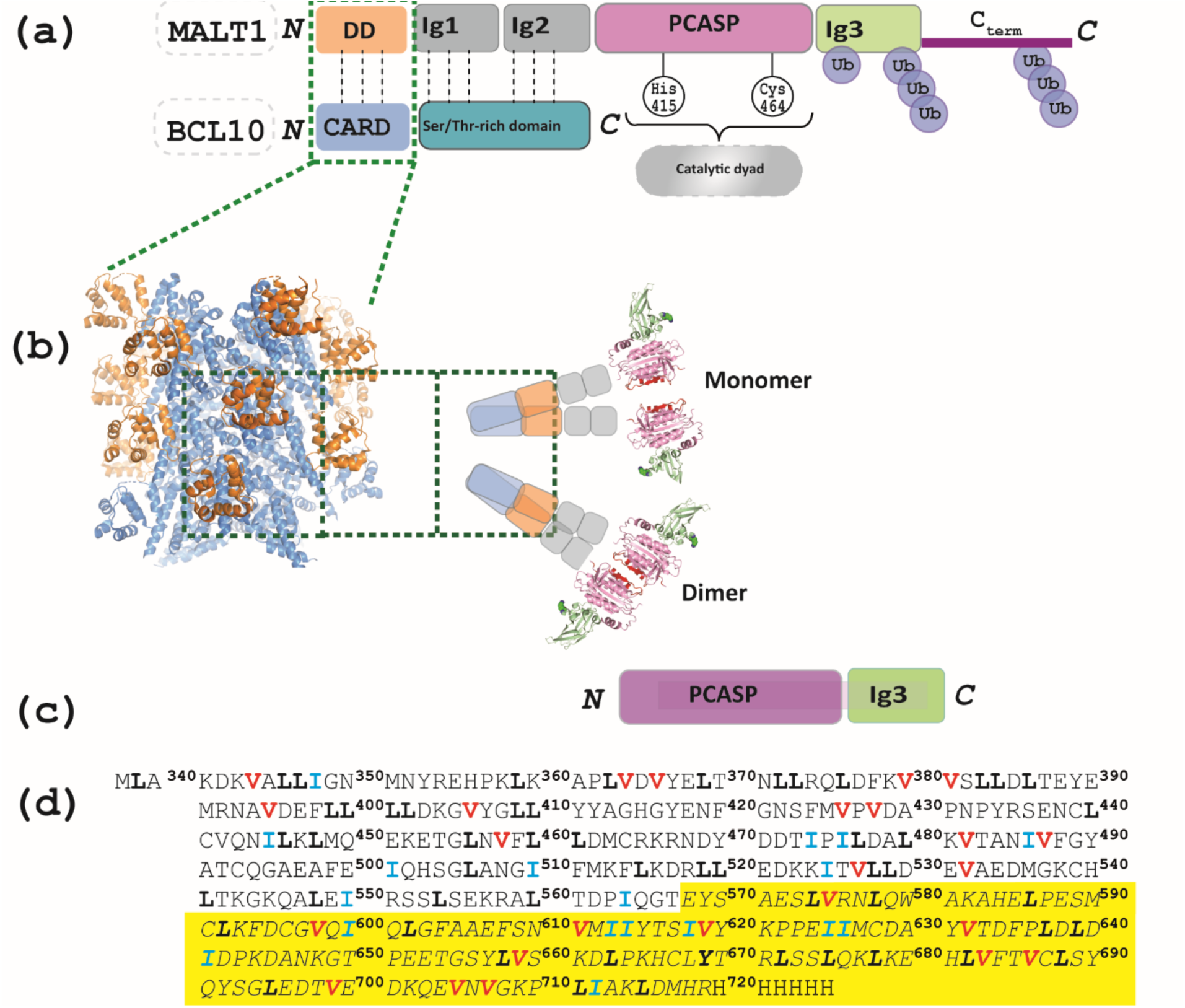
The MALT1-BCL-10 complex (MB10). **(a)** Protein domain structure: MALT1 forms a complex with the protein BCL10 through interactions between the DD and CARD domains. Sites for mono- and poly-ubiquitination are indicated. **(b)** The 4.5Å cryo-EM model of the MALT1-BCL10 complex (MB10) (Schlauderer *et al*., 2018) reveals a filament core formed by BCL10 (blue) into which the MALT1 DD domain (orange) is inserted. Notably, none of the other domains of MALT1 are resolved in the cryo-EM study due to their high flexibility. It has been proposed that the putative arrangement of the MALT1 complex is in equilibrium between monomer and dimer forms. **(c)** MALT1 construct used within this study. **(d)** Sequence of the MALT1(PCASP-Ig3)_339–719_ construct, with the methyl-containing residues I, L and V labelled in blue, black and red, respectively, and the sequence covering the Ig3 domain in yellow.

MALT1, initially identified as a distant relative of caspases during efforts to trace their evolutionary origin (Uren *et al*., 2000), has been postulated to exist as an inactive monomer that becomes activated through oligomerization/dimerization in the CBM complex(**Figure 1b**). It has been demonstrated that MALT1 constructs, initially obtained as monomers through size exclusion chromatography under physiological salt concentrations, show a tendency to spontaneously form dimers in solution (Wiesmann *et al*., 2012). This finding aligns well with the concept that dimerization is crucial for the formation of an active MALT1 complex (Yu *et al*., 2011, Wiesmann *et al*., 2012). *In vitro* studies provided further support for the significance of dimerization in MALT1 activation. Notably, elevated MALT1 activity was observed in the presence of the kosmotropic ammonium citrate buffer (Hachmann *et al*., 2012, Pelzer *et al*., 2013, Coornaert *et al*., 2008), known for activating initiator caspases by promoting their dimerization (Roschitzki-Voser *et al*., 2012, Boatright *et al*., 2003), which pinpoints the crucial role of dimerization in MALT1 activation. However, this enhanced activity was completely abolished in the E549A mutant, affecting both MALT1 dimerization and activity (Cabalzar *et al*., 2013).

To elucidate how the formation of the CBM complex activates MALT1, pioneering biochemical and crystallographic studies have been previously conducted *in vitro* (Wiesmann *et al*., 2012, Yu *et al*., 2011). Truncated ligand-free constructs of MALT1, encompassing the PCASP and the adjacent Ig3 domain (MALT1(PCASP-Ig3)_339–719_ (**Figure 1c**), were purified and separated into monomeric and dimeric forms using size-exclusion chromatography for subsequent structural analysis. The three-dimensional structure of the dimeric form was determined to 1.8 Å resolution (PDB code 3V55), revealing an inactive self-inhibited configuration (Wiesmann *et al*., 2012). Notably, no MALT1 monomers were observed to crystallise under these conditions (Wiesmann *et al*., 2012, Yu *et al*., 2011).

The PCASP domain of MALT1 harbours the histidine/cysteine catalytic dyad H415/C464 within its active centre (**Figure 1a**), that triggers cleavage activity in T lymphocytes upon antigen receptor engagement (Coornaert *et al*., 2008, Rebeaud *et al*., 2008). The catalytic activity of PCASP has been previously established, with a particular specificity towards peptide substrates featuring an arginine residue at position 1 (P1) (PDB code 3V4O) (Hachmann *et al*., 2012, Snipas *et al*., 2004). The crystal structure of an active conformation has also been previously determined, as an irreversibly modified dimer of MALT1(PCASP-Ig3)_339–719_ in complex with a substrate-mimicking inhibitor (Wiesmann *et al*., 2012, Yu *et al*., 2011, Rebeaud *et al*., 2008). It was concluded from these pioneering studies that the transition from the inactive to active conformation of MALT1, which was induced by the inhibitor, involves profound structural changes propagating throughout the PCASP domain, particularly affecting the active site and dimerization interface, along with a long-range of structural alterations. Notably, significant allosteric conformational changes were introduced in the Ig3 domain during PCASP activation and dimerization, a phenomenon that seemed to be accentuated by binding of the inhibitor to the active site.

Mono-ubiquitination of MALT1 highlights an alternative allosteric activation pathway connecting the PCASP and Ig3 domains (Schairer *et al*., 2020, Pelzer *et al*., 2013, Cabalzar *et al*., 2013). Ubiquitin (Ubq) binds to the Ig3 via a negatively charged surface patch, including residues E696 and D697, opposite the Ig3-protease domain interaction surface (Schairer *et al*., 2020). Mutation of Y657 in Ig3 induces coordinated conformational changes in the loop connecting K644 to Y657, and the active site. This mutational result led to the proposal that Ubq’s covalent attachment to K644 may structurally alter the loop extending from K644 to Y657, potentially affecting hydrophobic inter-domains interactions, such as those between Y657 on Ig3, and both L506 and Y367 in PCASP. These interactions were notably altered upon binding of the substrate analogue z-VRPR-fmk to the active site of MALT1 (Yu *et al*., 2011). Emphasising the role of Y657 as a key mediator in signalling between Ig3 and protease domains, this previous study suggested that Ubq attachment to K644 induces allosteric conformational changes in the Ig3-protease interface, ultimately enhancing MALT1 activation.

These previous studies explored also the complex network of allosteric communication within and between different protein domains in MALT1(PCASP-Ig3)_339–719_. Allostery which is a phenomenon that couples a ligand binding at one site of a protein with a conformational or dynamic change at a distant site, is crucial for biological signalling. A thorough understanding and ultimately potential targeting of such regulatory allosteric pathways could represent a key step for the adequate design of new high specificity drugs and disease treatments. Significant progress has been made in identifying a specific allosteric target pocket localized between the PCASP and Ig3 domains. This advancement has facilitated the development of compounds that inhibit MALT1 function in a non-competitive, allosteric manner (Hamp *et al*., 2021, Schlauderer *et al*., 2013, Quancard *et al*., 2019). However, relying solely on a static structural view (Changeux & Edelstein, 2005) may be insufficient, and several challenges persist in identifying and characterising other functional allosteric sites in MALT1. Previous studies on the nature of allostery (Motlagh *et al*., 2014, Jiang & Kalodimos, 2017) suggest that long-range communication is often not limited to changes in the mean protein conformation, but involve also a redistribution of structural ensembles, and changes in amplitudes and time scales of dynamic fluctuations about the mean conformation (Wand, 2001, Tsai *et al*., 2008).

NMR spectroscopy provides a unique possibility to investigate how protein motions contribute to the transmission of allosteric signals (Wand, 2001, Zhuravleva *et al*., 2007, Henzler-Wildman & Kern, 2007, Wieteska *et al*., 2017, Strotz *et al*., 2020, Köhler *et al*., 2020). It offers important insights into structural details, allows the characterisation of molecular motions across various time scales (Palmer, 2004, Mittermaier & Kay, 2009), provides access to sparsely populated conformational states (Baldwin & Kay, 2009), and to residue-specific information about conformational entropy (Motlagh *et al*., 2014, Wand, 2013, Tzeng & Kalodimos, 2011). From this perspective, allosteric transitions are embedded within the composition of structural ensembles, whose relative populations can be modulated under different conditions (Motlagh *et al*., 2014, Shukla *et al*., 2023). Thus, the ensemble allosteric model postulates that the potential for various allosteric mechanisms is inherently pre-encoded in protein structure ensembles (Motlagh et al., 2014, Motlagh et al., 2012).

Long molecular dynamics (MD) simulations in explicit solvent and modelling utilising such advanced methods such as AlphaFold (AF) (Jumper *et al*., 2021) are methods that can be used to create conformational ensembles at atomic resolution. Additionally, even though AF was initially designed to predict three-dimensional protein structures with a high quality comparable to those obtained using experimental methods (Jumper *et al*., 2021), recent advancements demonstrated also its capability to generate multiple conformations and predict dynamic regions at the individual residue level (Wallner, 2023, Ma *et al*., 2023, del Alamo *et al*., 2022, Wayment-Steele *et al*., 2023). Despite the vast number of publications on MALT1 during the last decade, there are to the best of our knowledge no reports of NMR dynamic studies on the paracaspase MALT1(PCASP-Ig3)_339–719_ nor on any similar construct. Here, we studied the dynamics of MALT1(PCASP-Ig3)_339–719_, and of the E549A mutated variant in solution. We have previously presented the almost complete assignment of the ^15^N/^13^C/^1^H backbone (Unnerstale *et al*., 2016) and of the IVL-Methyl side chains (Han *et al*., 2022) for the apo form of human MALT1(PCASP-Ig3)_339–719_ using high-resolution NMR. By using ^15^N-backbone *R*_1_- and *R*_2_-relaxation, CCR rates, we here delved into crucial information concerning overall rotational diffusion, internal dynamics, and inter-domain motion. Additionally, we employed the recently developed AF and an array of other structure modelling methods to identify and characterise conformational ensembles for MALT1(PCASP-Ig3)_339–719_, which we thereafter used to assess the allosteric pathways using the principal component analysis (PCA).

## 2.0 MATERIALS AND METHODS

### 2.01 Theoretical background

The theory of nuclear magnetic relaxation is well described (Korzhnev *et al*., 2001, Cavanagh *et al*., 2007, Jarymowycz & Stone, 2006, Luginbühl & Wüthrich, 2002). In the two-spin ^1^H-^15^N system, the measured relaxation parameters *R*_1_, *R*_2_, the ^15^N–(^1^H) nuclear Overhauser effect (NOE), and the cross-correlated relaxation *η_xy_* are connected to the molecular dynamics via the spectral density function *J(ω)*:

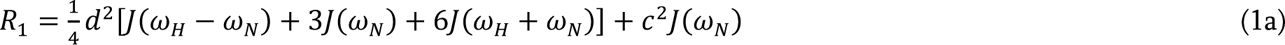

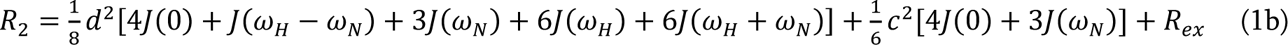

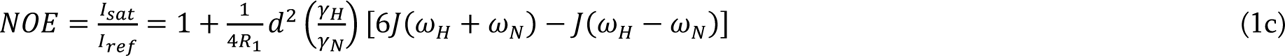

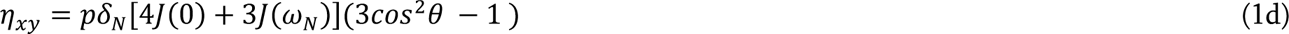

where 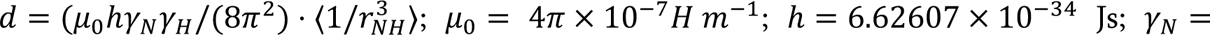 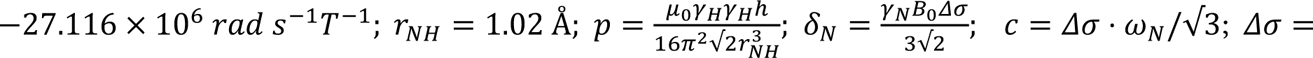 172 *ppm* is the ^15^N chemical shielding anisotropy; *θ* = 17° is the angle between the ^15^N−H vector and the unique principal axis of the ^15^N chemical shift tensor; and *R_ex_* is a contribution to *R*_2_ due to conformational exchange in the slow μs-ms time scale. The most simplified model of the spectral density function corresponds to an isotropic molecular rotational diffusion in solution and does not account for the intramolecular motions:

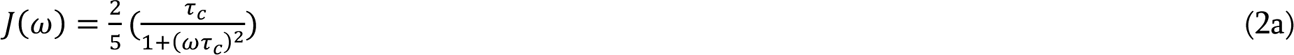

where *τ*_C_ is the global molecular rotational correlation time.

In the case of anisotropic rotational diffusion, there is dependence of the spin-relaxation of a given ^15^N nucleus on the orientation of the NH-bond in the molecular coordinate frame, hence the rotational diffusion has directional dependence. The anisotropic model is represented as a sum of five terms:

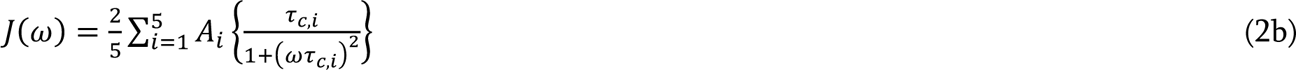

where correlation times *τ_c,i_* d the corresponding weights *A*_i_ depend on parameters of the rotational diffusion tensor ***D*** and the orientation of the H-N vector relative to the tensor principal axes. Tensor ***D*** is defined by six independent values including its principal components *D_i_* (*i = x, y, z*) and the Euler angles *α, β, γ* that characterise the directions of its principal axes in the reference molecular frame.

The effective overall rotational correlation time is computed according to:

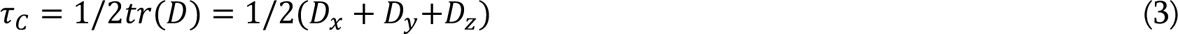

For the axially symmetric rotational diffusion, Eq. (2b) simplifies by reducing from five to three terms (Korzhnev *et al*., 2001, Barbato *et al*., 1992).

When the orientations of the HN vectors are known from the protein 3D structure, the tensor parameters can be calculated using experimental *R*_2_/*R*_1_-ratios for multiple HN vectors (Korzhnev *et al*., 2001, Fushman *et al*., 2004). The *R*_2_/*R*_1_ method was successfully validated for protein systems up to 25 kDa (Cavanagh *et al*., 2007). For larger systems, the TRACT approach (Lee *et al*., 2006) was proposed, which uses measurements of rates of the cross correlated relaxation *η_xy_*. The TRACT approach makes use of the relaxation interference between the two dominating relaxation mechanisms: the dipole-dipole interaction between amide ^15^N and its directly attached proton, and the amide ^15^N chemical shielding anisotropy (CSA). For ^15^N nuclei attached to a proton in the *β* spin state, the sum of the dipolar and CSA relaxations is smaller than for ^15^N nuclei attached to a proton in the *α* spin state, and thus the two types of ^15^N nuclei relax at very different rates. While experimental *η_xy_*are more susceptible to the impact of internal motions compared to *R*_2_/*R*_1_ ratios, it is worth noting that the cross-correlated relaxation is immune to the conformational exchange in the microsecond-millisecond time scale.

The relationship between *η*_xy_ and the apparent residue-specific rotational correlation time *τ*_C_ is provided by the equation:

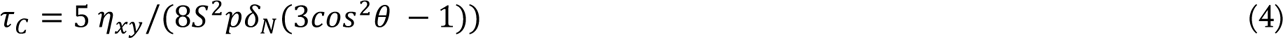

where *S*^2^ is the model free order parameter allowing correction for the fast picosecond motions of the HN vector in the molecular coordinate frame. The above equation is a simplified equation [40], where in the expression for *η_xy_* we kept only the terms proportional to *J*(0) and *J*(*ω_N_*). We estimated that contribution from the other terms related to *J*(*ω*) at the higher frequencies is less than 1%.

### 2.02 Expression of isotope-labelled MALT1(PCASP-Ig3)_339–719_ and preparation of NMR samples

The DNA sequence encoding for the PCASP and Ig3 domains of human MALT1, corresponding to residues 338–719 (**Figure 1c**), and a C-terminal His6-tag was cloned into the expression vector pET21b (Novagen). The MALT1(PCASP-Ig3)_339–719_-his construct was transformed into *Escherichia coli* strain T7 express competent cells, and thereafter expressed in different isotopic labelling combinations in ^1/2^H, ^15^N, ^12/13^C-labelled M9 medium. Chemicals for isotope labelling (ammonium chloride, ^15^N (99%), D-glucose, ^13^C (99%) and deuterium oxide were purchased from Cambridge Isotope Laboratories, Inc. The E549A mutation was introduced using the QuikChange Lightning kit (Agilent, Santa Clara, CA, USA). All cloned constructs and introduced modifications were verified by DNA sequencing. Both wild-type MALT1(PCASP-Ig3)_339–719_ and MALT1(PCASP-Ig3)_339– 719_(E549A) were produced and purified as described here below. MALT1(PCASP-Ig3)_339–719_ was expressed in 1 L of D_2_O M9 medium using 3g/L of U-[^13^C,^2^H]-glucose (CIL, Andover, MA) as the main carbon source, and 1g/L of ^15^NH_4_Cl (CIL, Andover, MA) as the nitrogen source. One hour prior to induction, precursors were added to the growth medium as previously described (Tugarinov *et al*., 2006). For precursors, 70 mg/L alpha-ketobutyric acid, sodium salt (^13^C4, 98%, 3,3-^2^H, 98%) and 120 mg/L alpha-ketoisovaleric acid, sodium salt (1,2,3,4-^13^C4,99%, 3, 4, 4, 4, -^2^H 97%) (CIL, Andover, MA) were used. Bacterial growth was continued for 16 h at 16°C and cells were thereafter harvested by centrifugation. Cells were resuspended in lysis buffer 20 mM TrisHCl (pH7.6), 150 mM NaCl, 2mM DTT and lysed using an ultrasonicator, followed by centrifugation at 40,000 g for 30 min to remove cell debris. The supernatant was collected and incubated with Ni^2+^ Sepharose 6 Fast Flow (GE Healthcare) for 1h at 4°C. The target protein was eluted with a lysis buffer containing 200-500 mM imidazole. A Q-Sepharose HP column (GE Healthcare) was used to separate monomeric MALT1(PCASP-Ig3)_339–719_ from the dimer form. A final size exclusion chromatography (SEC) step was performed using a HiLoad 16/600 Superdex 200 prep grade column (GE Healthcare), with running buffer 20mM HEPES 7.4, 50mM NaCl, 1mM DTT. The final monomer MALT1(PCASP-Ig3)_339–719_ protein sample was subsequently exchanged to a buffer (10 mM Tris 7.6, 50mM NaCl, 2mM TCEP, 0.002% NaN3, 10% D_2_O) suitable for NMR experiments using gravity flow PD10 desalting columns (GE Healthcare). Final yields from a four litres M9 culture were approximately 8 mg, and the purified MALT1(PCASP-Ig3)_339–719_-his tagged proteins were concentrated to at least 0.3-0.5 mM for NMR data acquisition.

A notable problem during data collection was the sample stability, manifested in a loss of signal intensity by approximately 20-30% over a period of one week. Nevertheless, the signal drop did not lead to systematic errors in the acquired data and analysis, but rather resulted in slightly more scattered and noisy data, which is expected when the signal-to-noise ratio diminishes. To overcome this problem, we performed the following steps: i) all measurements were performed with a completely fresh MALT1(PCASP-Ig3)_339–719_ sample at each of the two fields, ii) decrease of the signal intensity with time was not accompanied by any shift or broadening of the peaks, indicating that there was no reason to believe that the populations of protein conformations changed during the measurements, iii) the measurements were performed using sorted NUS table, which converted the intensity drop into a small additional broadening of the ^15^N without any effect on the relaxation decay, iv) the relaxation parameter vs residue over the two fields were consistent.

### 2.03 NMR relaxation experiments and data processing

All NMR data were acquired at 298K on 800 and 900 MHz Bruker spectrometers equipped with TCI cryo probes which are optimised for triple resonance experiments on biological macromolecules. All spectra were processed using either the mddnmr (Orekhov & Jaravine, 2011) or the NMRPipe (Delaglio *et al*., 1995) softwares at the NMRbox server (https://nmrbox.nmrhub.org/) (Maciejewski *et al*., 2017) or with the TopSpin 4.2.0 Bruker. To determine diffusion tensor MATLAB (MathWorks) and version 7 of the MATLAB-based ROTDIF (Walker *et al*., 2004, Fushman, 2002) were used. In house MATLAB scripts used for treatment of the ROTDIF-calculations and all subsequent data analyses are available upon request.

### 2.04 Determination of ^15^N relaxation rates *R*_1_, *R*_2_ and ^15^N-(^1^H) nuclear Overhauser effect

A complete set of ^15^N relaxation rates *R*_1_, *R*_2_ and ^15^N-(^1^H) nuclear Overhauser effect (NOE) experiments for the backbone amides were acquired with the standard Bruker pulse sequences using TROSY (Transverse Relaxation-Optimised Spectroscopy Nuclear Overhauser Effect Spectroscopy) and sensitivity enhancement (Lakomek *et al*., 2012, Zhu *et al*., 2000) at 800 MHz and 900 MHz. The *R*_1_ and *R*_2_ spectra were recorded in the pseudo 3D mode with randomised order of the relaxation delays and the NUS table sorted in the ^15^N spectral dimension to mitigate signal decay during the experiment due to the slow sample degradation. The following 11 relaxation delays were used to measure *R*_1_ values at 800 MHz: 0.08, 0.16, 0.32, 0.48, 0.72, 0.88, 1.12, 1.36, 1.60, 2.00 and 2.40 s; and at 900 MHz: 0.08, 0.16, 0.32, 0.48, 0.72, 0.88, 1.12, 1.36, 1.76, 2.24 and 2.4 s. For the *R*_2_, the following nine relaxation delays were used at 800 MHz: <u>0.0058</u>, 0.0115, 0.0173, <u>0.0230</u>, 0.0288, 0.0346, 0.0403s; and the following 11 relaxation delays were used at 900 MHz: <u>0.0058</u>, <u>0.0115</u>, 0.0173, <u>0.0230</u>, 0.0288, <u>0.0346</u> and 0.0403s, where each of the underlined delays were used twice. NOE experiments were recorded at 800 MHz and used the ^1^H saturation time of 5s. Experimental parameters for *R*_1_, *R*_2_ and NOE measurements are summarised in Table 1.

**Table 1.** List of acquisition parameters used for NMR experiments.

| Experiments | Maximum evolution time, (ms)/ carrier frequency (ppm)/sweep width (ppm) |  |  | D1 (s) | Scans | NUS % | Time (h) |
| --- | --- | --- | --- | --- | --- | --- | --- |
|  | F3 | F2 | F1 |  |  |  |  |
| 3D $^1\text{H}$-$^{15}\text{N}$ SF-TROSY-NOESY<sup>(a)</sup> | 79.9($^1\text{H}$)/4.67/16.0 | 27.4($^{15}\text{N}$)/118.0/36.0 | 28.4($^1\text{H}$)/4.67/11.0 | 0.5 | 16 | 23 | 68 |
| 4D $^{13}\text{C}$-SF-HMQC NOESY<sup>(b)</sup> | F4<br>81.0($^1\text{H}$)/4.7/14.0 | F3/F2<br>9.8($^{13}\text{C}$)/17.0/18.0 | F1<br>19.7($^1\text{H}$)/4.7/1.8 | 0.7 | 8 | 10.5 | 84 |
| 3D pseudo TROSY-T1 R1<sup>(a)</sup> 3D trt1etf3gpsitc3d.3 R1<sup>(b)</sup> | 119.8($^1\text{H}$)/4.7/16.0 | 37.7($^{15}\text{N}$)/118.0/36.0 | - | 1.2 | 40 | 44 | 28 |
| | 70.9($^1\text{H}$)/4.7/16.0 | 37.6($^{15}\text{N}$)/118.0/32.0 | - | 1.2 | 40 | 50 | 33 |
| 3D pseudo TROSY-T2 R2<sup>(a)</sup> 3D trt2etf3gpsitc3d.3 R2<sup>(b)</sup> 3D | 79.8($^1\text{H}$)/4.7/16.0 | 37.7($^{15}\text{N}$)/118.0/36.0 | - | 1.5 | 56 | 45 | 23 |
| | 70.9($^1\text{H}$)/4.7/16.0 | 37.6($^{15}\text{N}$)/118.0/32.0 | - | 1.5 | 56 | 54 | 33 |
| NOE<sup>(a)</sup> 3D trnoetf3gpsi3d.3 3D pseudo TROSY-NOE | 79.8($^1\text{H}$)/4.7/16.0 | 34.2($^{15}\text{N}$)/118.0/36.0 | - | 6 | 40 | - | 27 |
| CCR-$^{15}\text{N}$<sup>(b)</sup> | 79.8($^1\text{H}$)/4.7/16.0 | 37.6($^{15}\text{N}$)/118.0/32.0 | - | 1.5 | 64 | - | 26 |

<sup>(a)</sup> Experiments performed on an 800MHz spectrometer.
<sup>(b)</sup> Experiments on 900MHz spectrometer

The relaxation rates were obtained by extracting peak volume from CcpNmr Analysis Assign while keeping the original peak position for all spectra in a relaxation decay data set. To extract the relaxation rates, monoexponential functions were fitted to the relaxation decays using MATLAB’s built-in nonlinear regression programs, with errors estimated from the covariance matrix. NOEs were calculated as the ratio of the peak intensities in the saturated and reference experiments, and the standard deviations for the NOEs were determined by propagating the errors of intensities estimated from the base plane noise determined from regions without any peaks. All error estimates are reported as one standard deviation (SD).

### 2.05 Determination of ^15^N cross-correlated transverse relaxation of backbone

The ^15^N cross-correlated transverse relaxation experiment (CCR) was run for both MALT1(PCASP-Ig3)_339–719_ and mutated MALT1(PCASP-Ig3)_339–719_(E549A). The CCR experiment (Table 1) was performed at 900 MHz using 2D ^1^H-^15^N TROSY and anti-TROSY experiments (Vallurupalli *et al*., 2007) with relaxation delays Δ of 0 and 10ms. The cross-correlation rates *η_xy_* were calculated from two peak intensities *I^α^* in the TROSY spectrum, and *I^β^* in the anti-TROSY for

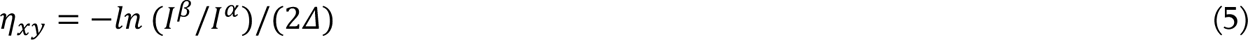

The initial estimate of the error in *η_xy_* was propagated from the difference between *I^α/β^* intensities measured for the relaxation delays Δ=0. However, since the estimate from just two repeated measures was too crude, we averaged the errors obtained for MALT1(PCASP-Ig3)_339–719_ and MALT1(PCASP-Ig3)_339–719_(E549A). We also regularised the values by using a minimal relative error of 5%.

### 2.06 Estimation of exchange contribution to the transverse relaxation rate *R*_2_

An assessment of the *R*_ex_-contribution to *R*_2_ can be made by comparing *R*_2_ with the transverse cross-correlated relaxation rate according to Eq. (1) (Palmer *et al*., 2001):

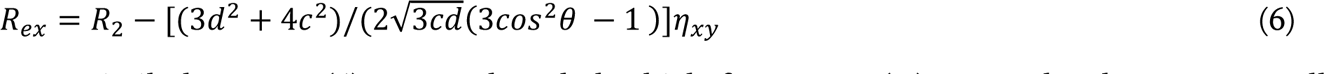

were, similarly to Eq. (4), we neglected the high frequency *J*(ω) terms that have very small contributions compared to the dominant *J*(0) and *J*(ω*_N_*) terms. The residue specific error in conformational exchange σ_Rex_ was determined by propagating the errors from *η*_xy_ and *R*_2_. In our qualitative analysis, we defined residues with statistically significant conformational exchange as those that show *R*_ex_/σ_Rex_ > 5.

### 2.07 Calculation of Rotational diffusion tensor

ROTDIF takes the ^15^N relaxation parameters and the ^1^H-^15^N bond vector coordinates from the MALT1(PCASP-Ig3)_339–719_ crystal structure (PDB code 3V55) as input. Hydrogens were added to the crystal structure using the *addh* function in the Chimera 1.16 software (Pettersen *et al*., 2004). For the illustration of the structures presented in this article, an AlphaFold-generated model was employed to replenish two regions (residues 495–497 and 469–480) within the PCASP domain that are missing in the crystal structure.

We used ROTDIF to fit the 900 MHz *R*_2_/*R*_1_, 800MHz *R*_2_/*R*_1_ and 900 MHz CCR data to the axially symmetric model of MALT1(PCASP-Ig3)_339–719_, and obtained the principal values of two tensor components D_XX_D_YY_ and D_ZZ_, and of the two Euler angles α and β. The standard errors of the fitted parameters were estimated using a resampling technique reminiscent of the *delete-d* jack-knife estimation (Mosteller & Tukey, 1977). Since the ROTDIF software requires *R*_2_/*R*_1_-ratios as input, we converted the *η_xy_*-derived residue specific *τ*_C_, (Eq. 4) to residue-specific *R*_2_/*R*_1_-ratios for 900 MHz spectrometers by inverting equation 6 in (Fushman *et al*., 1994):

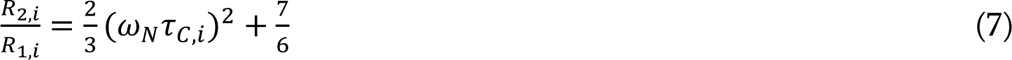

When calculating the residue specific *τ*_C_ using Eq. (4), we adjusted *S^2^* to 0.88. This value was adjusted by equating the average *τ*_C_ obtained from the CCR-measurements (*η_xy_*) with those from the 800 and 900 MHz *R*_2_/*R*_1_-ratio measurements. The used *S^2^* value is in line with the range 0.8-0.9 usually observed for structured parts of globular proteins and indicates a general good agreement between our *η_xy_*and *R*_2_/*R*_1_ data.

### 2.08 Selection of residues for diffusion tensor calculations

The selection of residues for the ROTDIF-optimization in MATLAB constitutes a crucial step in our analysis. In this study, we applied the following four selection criteria for choosing the amide group of a specific amino acid residue for the calculation of the rotational diffusion of MALT1(PCASP-Ig3)339–719 : i) we kept only residues with a relative *R*_2_/*R_1_* error of below 10%, 15% and 25% in the 900 MHz, 800 MHz and CCR data sets, respectively; ii) we excluded residues with B-factors for backbone nitrogen atoms in the crystal structure above 35 Å^2^. The average nitrogen backbone B-factor for the 368 residues (PDB code 3v55) is 33.3 ± 12 Å^2^, with a median of 29.6 Å^2^; iii) we excluded residues whose peaks strongly overlapped in the ^1^H-^15^N TROSY spectrum; iv) for reducing impact of milliseconds (conformational exchange) and nanosecond motions, we kept only residues within two standard deviations (SD) from the mean *R*_2_/*R*_1_-ratio for *R_2_, R_1_* 800/900 MHz data and 1.7 SD from the mean *R*_2_/*R*_1_-ratio for the CCR-data. These four selection criteria reduced the number of residues from 270 assigned and characterised protein peaks to 116 (900 MHz *R*_2_/*R*_1_), 115 (800 MHz *R*_2_/*R*_1_) and 105 (900 MHz CCR). These three sets of selected peaks were used for the calculations of the rotational diffusion tensor.

### 2.09 Estimation of diffusion tensors using statistical resampling

The experimental relaxation data for MALT1(PCASP-Ig3)_339–719_ have relatively poor signal-to-noise due to the large system size, and to degradation of the sample during experiment. The data quality worsens in the order *R*_2_/*R_1_* 900 MHz → *R*_2_/*R_1_* 800 MHz → CCR, and therefore we attributed the most explanatory power to the *R*_2_/*R_1_* 900 MHz data. The two other sets of data, *R*_2_/*R_1_* 800 MHz and CCR were used, extra carefully and mainly for the validation of the *R*_2_/*R_1_* 900 MHz data. Furthermore, as far as we know, the method of calculation of the diffusion tensor from CCR data was not described before and thus requires validation on a well characterized protein system with high-quality experimental data. Thus, the rotation diffusion tensor results obtained from CCR are used in this study only qualitatively. To perform the analysis with the noisy data we assessed the criteria for selecting amide HN pairs as outlined above, and performed the error analysis using the bootstrap resampling technique that resembles the *delete-d* jack-knife approach (Bonamente, 2017, Zaman & Alakus, 2015). When the number of HN pairs used for the analysis exceeded 100 (n > 100), the rigorous delete-*d* jack-knife approach which exhaustively and systematically excludes all possible combinations of *d* samples from the set, requires too many parameter optimizations. A very similar result was obtained much faster by the statistical bootstrap sampling of a relatively small (*i.e.* 10^2^ in our case) number of parameter optimizations. The parameter uncertainties are approximated by 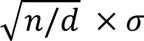, where in our case *σ* is the standard deviation of a tensor component D_XX_D_YY_ or D_ZZ_ obtained in 100 ROTDIF-simulations. The resampling method belongs to non-parametric statistics which do not require the assumption of normally distributed measurements. Statistical theory does not give a unified answer to what is a reasonable number for *d* and the degree of deletion seems to be model-dependent (Shi, 1988). We found that omission *d* = 50% represented the lower limit, as no good fitting of the rotational diffusion tensor could be obtained below this level and we therefore used d = 20% for our calculations.

### 2.10 Measurement of intramolecular interactions

Intramolecular amide-methyl interactions were confirmed by observing cross peaks in 3D SOFAST (SF), ^1^H-^15^N TROSY NOESY experiments. Additional intramolecular methyl-methyl interactions were obtained from 4D methyl-methyl SF HMQC-NOESY-HMQC experiments (Zwahlen *et al*., 1998). The acquisition parameters are presented in Table 1.

### 2.11 Chemical shift perturbation studies comparing wild-type MALT1(PCASP-Ig3)_339– 719_ and mutated MALT1(PCASP-Ig3)_339–719_(E549A)

Assignment of backbone NH chemical shifts in the mutated MALT1(PCASP-Ig3)_339–719_(E549A) was based on assignment of the wild-type MALT1(PCASP-Ig3)_339–719_ previously reported by us (Unnerstale *et al*., 2016), and additional sets of 3D TA-acquisition experiments (Isaksson *et al*., 2013, Jaravine *et al*., 2008), with parameters described in (Unnerstale *et al*., 2016) and deposited in the Biological Magnetic Resonance Data Bank (http://www.bmrb.wisc.edu/) with the BMRB accession code 52265. Excellent reproducibility of the samples was confirmed by the perfect overlay of the spectra (**Figure S3**). Effective chemical shift perturbations were determined for the comparison of the chemical shifts of mutated MALT1(PCASP-Ig3)_339–719_(E549A) and wild type MALT1(PCASP-Ig3)_339–719_, and was calculated as Δδ_eff_ = ([Δδ(^1^H)]^2^ + [0.16Δδ(^15^N)]^2^)^1/2^.

### 2.12 AlphaFold, PCA, ESMFold2, RoseTTAFold2 and NMA analyses

Ensembles of MALT1(PCASP-Ig3)_339–719_ structures were generated using AlphaFold version 2.3.1, and a massive sampling technique proposed by Wallner (Wallner, 2023). A total of 2000 structures were predicted with AF model *monomer_ptm* without templates, using dropouts at the inference stage. The dropout scheme was kept the same as during the AF training. The PCA (Kitao & Go, 1999) was performed using the *Scikit-learn* Python library (Pedregosa *et al*., 2011). To analyse the set of structures, we performed the PCA in Cartesian coordinates for two sets of atoms: (a) PCAa, which includes the N, C_α_, C, O atoms of protein residues 344 to 717, excluding flexible N- and C-termini, and all side-chain heavy atoms of amino acids exhibiting multiple rotamers in ξ_2_ torsion angles, including residue W580; (b) PCAb, akin to PCAa, but focusing only on the stretches of residues 579-584 and 652-659 including site chains of W580 and Y657. All structures were aligned using the PDB Superimpose module from Biopython (http://citebay.com/how-to-cite/biopython/). The alignment was done based on the Cα atoms of the stretches of residues 344-467, 484-500 and 510-562. These residues were selected for the performed alignment due to their location in the PCASP domain and high pLDDT scores.

For PCAa, the eigenvalues for the first three components were 0.48, 0.17, and 0.08. Thus, the analysis was limited to the 1^st^ and 2^nd^ principal components. For PCAb, the eigenvalues for the first three components were 0.74, 0.13 and 0.04. We kept the first two principal components, which notably caught rotamer changes for residues W580 and Y657. An inverse transformation was applied to the first and second principal components, converting them back into their original 3D coordinates, and reconstructing the specific molecular motions they represent. Normal mode analysis (NMA) was performed using the online server elNémo (Suhre & Sanejouand, 2004, Bauer *et al*., 2019). The crystal structure of MALT1(PCASP-Ig3)_339–71_9 (PDB code 3V55) was used as input, and all program parameters were left as default. The first three lowest frequency normal modes were used for further analysis. Both ESMFold2 (Lin *et al*., 2023) and RoseTTAFold2 (Baek *et al*., 2023, Liu *et al*., 2022) were used with default parameters to produce MALT1(PCASP-Ig3)_339–719_ structural models.

### 2.13 Molecular dynamic *simulations*

MD simulations were performed in Gromacs version 2023.1 (Abraham *et al*., 2015). All MD simulations were performed using the all-atom force field charmm36-mar2019_cufix.ff [https://github.com/intbio/gromacs_ff/tree/master/charmm36-mar2019_cufix.ff] with a refinement of the Lennard-Jones parameters (CUFIX) (Yoo & Aksimentiev, 2018) both for protein and, as recommended, TIP3P water model. For the initial structure, we used the MALT1(PCASP-Ig3)339– 719 structure predicted by model 1.1 of AlphaFold (Jumper *et al*., 2021) using ColabFold version 1.5.5 (Mirdita *et al*., 2022) when run with default parameters without templates. The protein was centred in a periodic 104Å cubic box, and 55 Na^+^ and 41 Cl^-^ ions were added to the system to emulate ionic strength and achieve electro-neutrality as the protein had a total charge of -14. Since the charge of the protein residues was calculated according to pH=7.6, all histidine residues displayed neutral charges. Long-range electrostatic and van der Waals 10Å cut-off were used. The structure was relaxed through a process called energy minimization to ensure a reasonable starting structure in terms of geometry and solvent orientation. Convergence was achieved at a maximum force of less than 1000 kJ/mol/nm in any atom. Equilibration was conducted in two phases, namely NVT ensemble (with protein molecular models heating from 0 to 300 K for 100 ps), and NPT ensemble for 1000ns until the system became well equilibrated and reached a plateau in the root mean square deviation (RMSD) values (**Figure. S6**). A Berendsen-type thermostat and barostat were used. Hydrogen-containing covalent bonds were constrained using the SHAKE algorithm and time steps of 2 fs were used. Following the equilibration, MD simulations were continued as a production run for 2000 ns under the same conditions. System stability was calculated using standard tools available in Gromacs (Abraham *et al*., 2015) including control of the temperature, pressure, energy, secondary structure, the box border, and RMSD. The PCA analysis was performed as described above for the AF structures.

## 3.0 RESULTS

### 3.1 MALT1(PCASP-Ig3)_339–719_ forms a monomer in solution

Several three-dimensional structures have been reported for different separate regions of the multidomain MALT1 (Wiesmann *et al*., 2012, Yu *et al*., 2011, Qiu & Dhe-Paganon, 2011). The low-resolution cryo-EM model of the BCL10-MALT1 complex (Schlauderer *et al*., 2018) (**Figure 1b**) unveiled the organisation of MALT1 N-terminal DD domains within the CBM filament. However, this cryo-EM structure did not provide any information about the configuration of the other regulatory and PCASP domains, likely due to their inherent conformational dynamics. All the previously determined crystal structures of MALT1(PCASP-Ig3)_339–719_ (**Figure 1c**) in the apo-form or in complex with ligands have the same dimerization interface of about 1000Å^2^ (**Figure 1b**). Up to now, there is to our knowledge no available three-dimensional structure of the monomeric form of MALT1(PCASP-Ig3)_339–719_ in solution.

NMR provides an alternative way, compared to gel filtration analyses, to discriminate monomeric and multimeric protein forms in solution, through the direct estimation of the rotational correlation time *τ*_C_. The experimental *τ*_C_ can be compared with the values predicted using an empiric equation that takes into account the temperature and molecular weight (Cavanagh *et al*., 2007). Such estimates gave *τ*_C_ values of 26.6ns and 52.9ns for the monomeric and dimeric forms of MALT1(PCASP-Ig3)_339–719_, respectively. The former value agrees well with our experimental results (Table 2), thus confirming the predominantly monomeric form of MALT1(PCASP-Ig3)_339–719_ in solution. Another way to address this topic is to compare the NMR-derived *τ*_C_ values for the wild-type and the E549A-mutated forms of MALT1(PCASP-Ig3)_339–719_. The mutation which is located within the dimerization interface of MALT1(PCASP-Ig3)_339–719_ prevents dimerization and stabilises the molecule in its monomeric state in solution (Cabalzar *et al*., 2013).

**Table 2.** Hydrodynamic characteristics of the PCASP and Ig3 domains in MALT1(PCASP-Ig3)_339–719_.

| Data set | Domain | # Res | $\tau_C$ <sup>(a)</sup> | $D_{XX}D_{YY}$ <sup>(b)</sup> | $D_{ZZ}$ <sup>(b)</sup> | Anisotropy <sup>(c)</sup> | $\alpha$ <sup>(d)</sup> | $\beta$ <sup>(d)</sup> |
| --- | --- | --- | --- | --- | --- | --- | --- | --- |
| | PCASP | 55 | $27.4 \pm 1.5$ | $0.52 \pm 0.02$ | $0.78 \pm 0.04$ | $1.48 \pm 0.05$ | 18 | 43 |
| 900<br>MHz<br>$R_2/R_1$ | Ig3 | 55 | $25.5 \pm 1.5$ | $0.62 \pm 0.02$ | $0.72 \pm 0.04$ | $1.17 \pm 0.05$ | 47 | 131 |
| | full | 116 | $26.5 \pm 1.2$ | $0.56 \pm 0.02$ | $0.77 \pm 0.03$ | $1.37 \pm 0.04$ | 11 | 59 |
| 800<br>MHz<br>$R_2/R_1$ | PCASP | 57 | $28.3 \pm 1.9$ | $0.52 \pm 0.02$ | $0.74 \pm 0.04$ | $1.43 \pm 0.05$ | 15 | 41 |
| | Ig3 | 52 | $26.8 \pm 2.5$ | $0.57 \pm 0.04$ | $0.73 \pm 0.05$ | $1.28 \pm 0.06$ | 14 | 62 |
| | full | 115 | $27.1 \pm 1.7$ | $0.56 \pm 0.03$ | $0.73 \pm 0.04$ | $1.32 \pm 0.05$ | 7 | 64 |
| 900<br>MHz<br>CCR | PCASP | 44 | $28.0 \pm 2.8$ | $0.54 \pm 0.03$ | $0.70 \pm 0.06$ | $1.29 \pm 0.07$ | N/A | N/A |
| | Ig3 | 52 | $27.8 \pm 2.1$ | $0.56 \pm 0.03$ | $0.68 \pm 0.05$ | $1.22 \pm 0.06$ | N/A | N/A |
| | full | 105 | $27.8 \pm 1.5$ | $0.55 \pm 0.02$ | $0.69 \pm 0.04$ | $1.25 \pm 0.05$ | N/A | N/A |

- a) Axially symmetric rotational correlation time (in ns) of the molecule. Error propagated from $D_{zz}$ and $D_{xx}D_{yy}$ using equation 3. - b) Principal values (in $10^7 \text{ s}^{-1}$ ) of the rotational diffusion tensor. - c) Degree of anisotropy of the diffusion tensor $D_{zz}/D_{xx}D_{yy}$ . - d) The angles cannot be reliably calculated in the resampling procedure for the 900 MHz CCR data due to poor data quality.

First, we investigated whether the E549A mutation triggers significant conformational changes in the PCASP domain, which should be revealed by differences in peak positions in the ^1^H-^15^N TROSY spectra for the apo form of MALT1(PCASP-Ig3)_339–719_ and the E549A variant (**Figure S1**). Comparison of the spectra revealed that the most significant changes in the chemical shifts were observed only in the vicinity of the mutation site, indicating an expected alteration in local environments rather than extensive conformational changes in the mutated protein. Secondly, we compared the cross correlated transverse relaxation rates *η*_xy_ of the protein amide groups in MALT1(PCASP-Ig3)_339–719_ (E549A) and wild-type MALT1(PCASP-Ig3)_339–719_ (**Figure S2d**). The *η*_xy_ values observed for the two molecules are statistically very similar. Furthermore, since *η*_xy_ is related to the apparent residue-specific rotational correlation time *τ*_C_, as described in equation (4), it follows that the values of *τ*_C_ are also similar between MALT1(PCASP-Ig3)_339–719_(E549A) and wild-type MALT1(PCASP-Ig3)_339–719_ (**Table 2**). Noteworthy is that the overall correlation time of MALT1(PCASP-Ig3)_339–719_ estimated from *η*_xy_ values is in agreement with the ‘empirically’ predicted values for the monomer. This finding supports our conclusion that the overall correlation time of MALT1(PCASP-Ig3)_339–719_ corresponds to a monomer as the main form in solution.

### 3.2 Contribution of the slow chemical exchange in the transverse relaxation R2 of backbone

The conformational exchange *R_ex_*, occurring in the protein at rates ranging from tens to thousands of inverse seconds can contribute significantly to the measured transverse relaxation rates *R_2_* (See equation 1b in material and methods section). Consequently, *R_ex_* can exert a notable influence on the *R_2_/R_1_* analysis of molecular rotational diffusion. In our efforts to exclude residues that exhibited a statistically significant *R_ex_* contribution to *R_2_* from the below discussed *R_2_/R_1_* analysis, we calculated *R_ex_* values using Eq. 6.

The most substantial exchange was observed in three specific regions corresponding to the stretches of residues 470-484, 492-510 and 560-590 (**Figure 2a**). These stretches are localized within the extensive loop region in the MALT1 dimerization interface, comprising the interdomain linker, including the α1-helix, and the region near the active site in the PCASP domain. Importantly, this observed significant exchange corresponds very well to regions in which residues are either absent or exhibit high B-factors in the crystal structures of MALT1(PCASP-Ig3)_339–719_ (**Figures 2a and 2b**). Another important observation is that the baseline values of *R*_ex_/σ*_R_*_ex_ for all residues, except those involved in *R_ex_*, exhibit remarkable similarity between the PCASP and Ig3 domains. This observation is important for the analysis of individual diffusion tensors presented here below.

**Figure 2.**
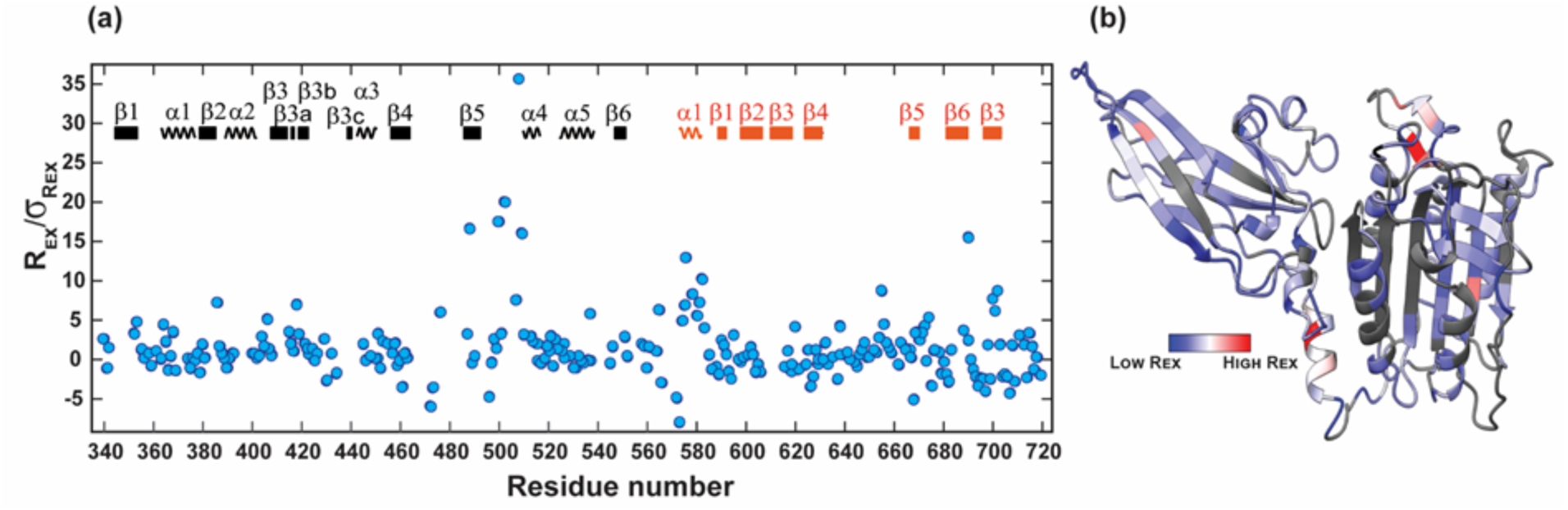
Conformational exchange in mille-micro-second time scale. **(a)** *R*_ex_/σ*_R_*_ex_ ratios. The secondary structure elements are indicated and numbered for each domain with domains, corresponding to PCASP and Ig3 in black and red, respectively. **(b)** *R*_ex_ values from the 900 MHz data were mapped on the crystal structure of MALT1(PCASP-Ig3)_339–719_.

### 3.3 Determination of rotational diffusion parameters from relaxation data

Global tumbling provides important insights into the size and shape of proteins in solution. This aspect is especially pertinent in the case of protein homo and hetero complexes and, as demonstrated in the present study, can even shed light on interdomain motions of multidomain proteins (Barbato *et al*., 1992). The process that we used to obtain reliable diffusion tensors is explained in the material and methods section. **Figure 3** shows the ratios of the transverse and longitudinal relaxation rates *R*_2_/*R*_1_, which are used for calculations of the diffusion tensors as well as the local per-residue rotational correlation times *τ*_C,i_ (**Figures 3b and 3c**) and heteronuclear ^1^H-^15^N NOEs (**Supplementary Figures S2a, S2b and Tables S1-3**). The *τ*_C,i_ and/or the ^1^H-^15^N NOE values that are significantly lower than average are depicted in **Figures 3c and 3d**, indicating the regions (*i.e.* the α1 helix and the three loops close to the catalytic site) that display significant mobility in the pico-nano second time range. It should be noted that the corresponding residues were not used in the diffusion tensors calculations.

**Figure 3.**
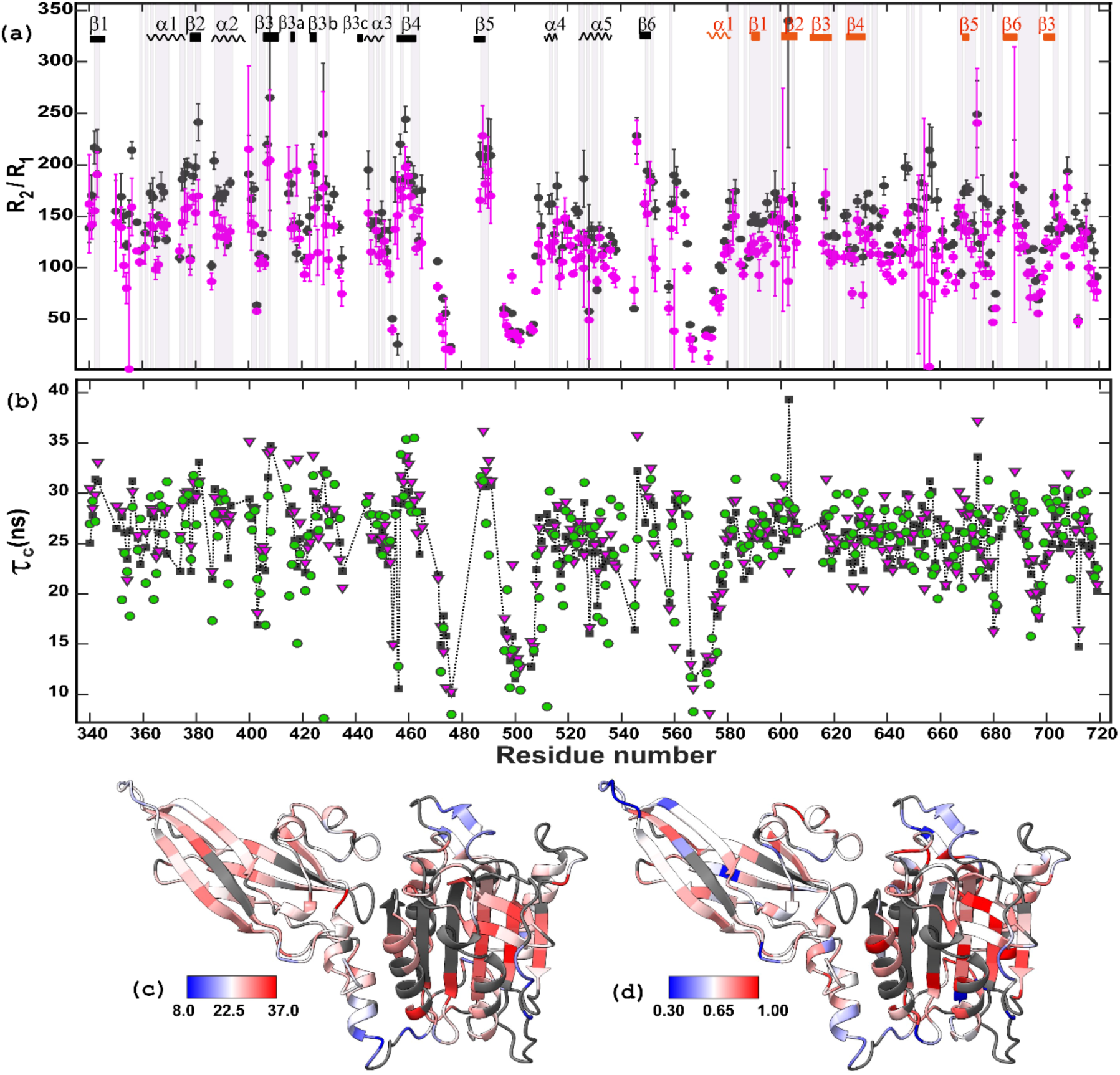
Residue specific dynamics in the nano-picosecond time scale. **(a)** ^15^N *R*_2_/*R*_1_ ratios for MALT1(PCASP-Ig3)_339–719_. Black and magenta dots represent 900 and 800 MHz data, respectively. Secondary structure elements are indicated and numbered for each domain at the top of the panel, with PCASP in black and Ig3 in red. α1 at residue 574 marks the division between the PCASP and Ig3 domains. Residues highlighted by grey vertical bars in panel (a) are the 116 selected residues that were used for the hydrodynamic calculations for 900 MHz data. Error bars show one SD. **(b)** Local correlation time per residue *τ*_C,i_ for MALT1(PCASP-Ig3)_339–719_ calculated from the 900 MHz ^15^N *R*_2_/*R*_1_ ratio (black square), 800 MHz ^15^N *R*_2_/*R*_1_ ratio (magenta triangle) and cross-correlated transverse relaxation (green circles). *τ*_C,i_ were calculated from Eqs. 1 and 7. **(c)** Local correlation time ranging from 8 to 37ns, from 900 MHz ^15^N *R*_2_/*R*_1_ mapped on the crystal structure of MALT1(PCASP-Ig3)_339– 719_ (PDB code 3V55). **(d)** Heteronuclear NOE ranging from 0.3 to 1.0, from the 800MHz data mapped on the crystal structure of MALT1(PCASP-Ig3)_339–719_. The ^15^N-(^1^H)NOE data are presented in **Supplementary Figure S1**. Residues in black are unassigned.

The diffusion tensor components D_ZZ_ and D_XX_D_YY_ from resampled ROTDIF-optimisations are shown in **Figure 4**, separately for PCASP and Ig3 domains calculated using the 900 MHz ^15^N *R*_2_/*R*_1_ data, the 800 MHz ^15^N *R*_2_/*R*_1_ data and the 900 MHz CCR data, respectively. The diffusion tensor data for the full protein is provided in **Supplementary Figure S3**. As evident from **Figure 4** and **Table 2**, the diffusion tensors of the PCASP and Ig3 individual domains are systematically different from each other in all measured data, which clearly indicates a semi-independent movement of these domains relatively to each other in solution. The most obvious differences observed in the highest quality *R*_2_/*R_1_* 900 MHz data were still seen in the *R*_2_/*R_1_* 800 MHz and CCR data, demonstrating that all the gathered data are consistent within the experimental error margins. More specifically, the results from the 900 MHz ^15^N *R*_2_/*R*_1_ data revealed an anisotropy ratio (D_ZZ_/D_XX_D_YY_) of 1.48 and 1.17 for PCASP and the Ig3 domains, respectively. These ratios were 1.43 and 1.28 for the PCASP and Ig3, respectively, when assessing the 800 MHz ^15^N *R*_2_/*R*_1_ data. Finally, the corresponding values were 1.29 and 1.22 for PCASP and Ig3, respectively, when assessing the 900 MHz CCR data. In accordance to the *R*_2_/*R*_1_ values, the effective overall correlation times calculated for the PCASP domain were systematically higher compared to the Ig3 domain (**Table 2**). Thus, for the 900 MHz data, the *τ*_C_ values were 27.4±1.5ns and 25.5±1.5ns for PCASP and Ig3, respectively, while the 800 MHz *τ*_C_ values were 28.3±1.9ns and 26.8±2.5ns for PCASP and Ig3, respectively. The CCR-data revealed only a minor difference in correlation times, with values of 28.0±2.8ns and 27.8±2.1ns for PCASP and Ig3, respectively.

**Figure 4.**
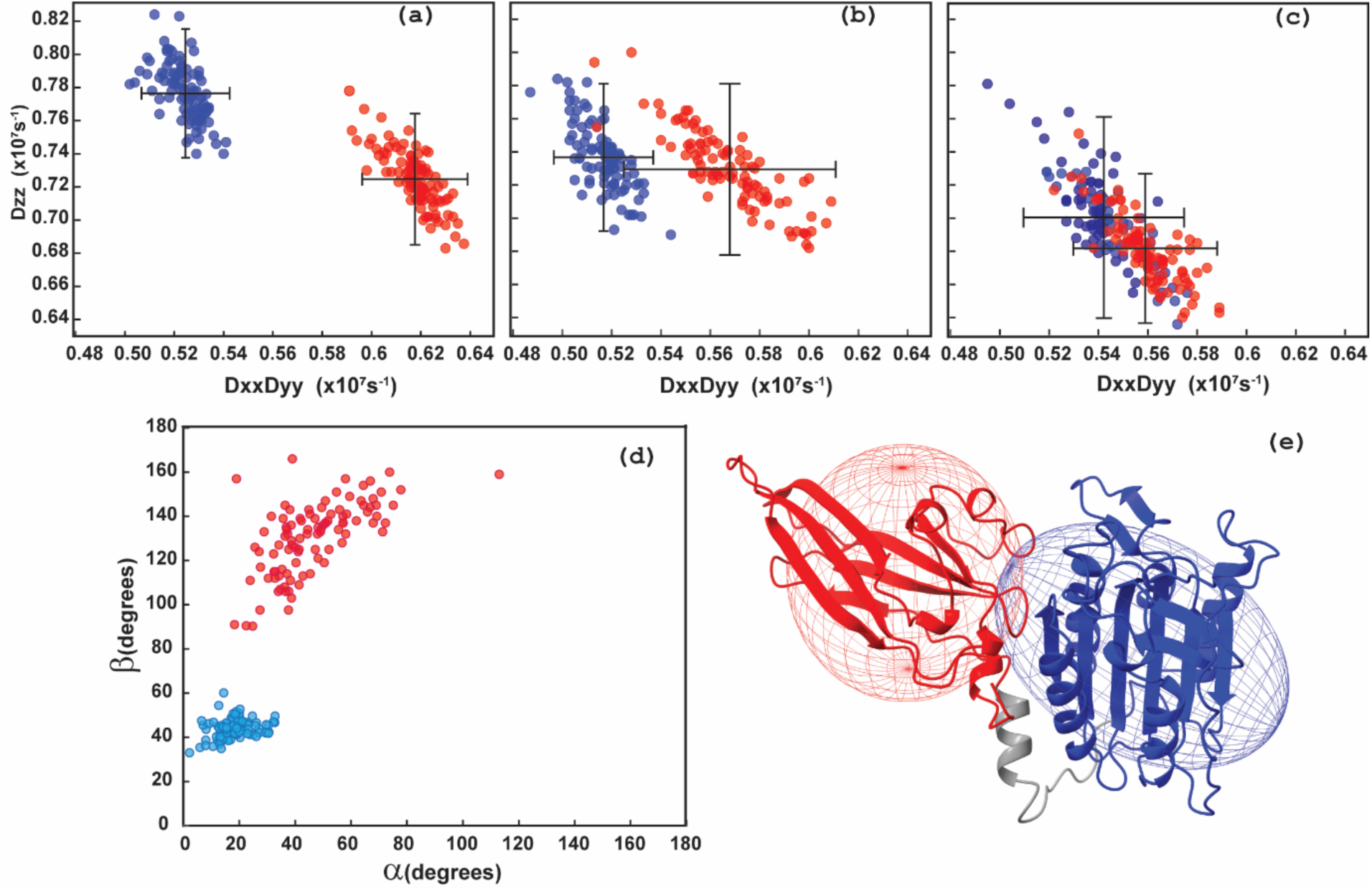
Diffusion tensor components D_ZZ_ and D_XX_D_YY_ from resampled ROTDIF optimisations. Each red and blue dot corresponds to a tensor calculated for one resampling instance for the Ig3 and PCASP domains, respectively, for **(a)** 900 MHz ^15^N *R*_2_/*R*_1_ data, **(b)** 800 MHz ^15^N *R*_2_/*R*_1_ data, **(c)** 900 MHz and cross-correlated transverse relaxation data. Error bars in (a)-(c) show one SD and are calculated taking the resampling fraction of *d=*20% into account. **(d)** α and β angles for the diffusion tensors for the 900 MHz ^15^N *R*_2_/*R*_1_ data set. The corresponding data for the full protein is shown in **Supplementary Figure S2**. **(e)** Ellipsoid representation of the diffusion tensors (900 MHz ^15^N *R*_2_/*R*_1_ data) mapped on the crystal structure of MALT1(PCASP-Ig3)_339–719_ with Ig3 and PCASP depicted in red and blue, respectively. The grey part of the structure comprising residues 563-582, including the loop and the α1 helix was excluded from optimization in ROTDIF. The procedures underlying the selection of residues in each domain is presented in the material and method section. The principal axis of the axially symmetric diffusion tensors crosses the poles of the ellipsoids. The diffusion tensor calculated for the full protein is shown in **Supplementary Figure S3**.

Although all three datasets agree within the experimental errors and consistently point to differences in the diffusion tensors of the two domains, a trend indicating a progressive reduction of these differences in the 900 MHz, 800 MHz and CCR data warrants a specific discussion. The first factor to consider is the potential impact of the contribution of the conformational exchange *R_ex_* to the *R*_2_ values in the 900 MHz and 800 MHz ^15^N *R*_2_/*R*_1_ data sets, as opposed to the 900 MHz CCR data. The major *R_ex_* contribution can be ruled out, as it would necessitate a longer apparent overall correlation time at 900 MHz than at 800 MHz. Furthermore, the *R_ex_* explanation of the opposite trends in the diffusion tensor values for PCASP and Ig3 requires significantly different *R_ex_* in the two domains, which stands in contrast to the same baseline level of the *R*_ex_/σ*_R_*_ex_ for the two domains (**Figure 2a**). Another potential factor may lie within the ROTDIF procedure, where the diffusion tensor is determined with the assumption of no significant motions in the nano-second time scale, which may affect the *R*_2_/*R*_1_ data primarily via the *R*_1_ values. Indeed, the tendency for higher *τ*_C_ values in *R*_2_/*R*_1_ analyses in lower magnetic fields has been previously reported for several proteins and attributed to the unaccounted effects of nanosecond motions (Orekhov *et al*., 1999). However, similarly to the above arguments for the *R_ex_* case, the nano-second motions explanation requires a rather unlikely assumption that these motions are significantly different in the stable/rigid part of the domains used for the tensor calculations. It should be noted that the trend for reduced differences between diffusion tensors in the two domains correlates with the decrease in data quality. Consequently, the primary emphasis in explaining the relative dynamics of PCASP and Ig3 should be based in our opinion only on the highest quality 900 MHz *R*_2_/*R*_1_-data set (**Figure 3a**). Finally, it suffices to conclude that the rotation diffusion tensors of PCASP and Ig3 are statistically different (**Figure 4**), which can be attributed to the semi-independent rotational diffusion of these domains in MALT1(PCASP-Ig3)_339–719_ in solution.

### 3.4 Methyl NOE-contacts reveal a pivotal point for the relative motions between PCASP and Ig3

To assess the semi-independent motion of MALT1(PCASP-Ig3)_339–719_ in its apo form, we examined and analysed the NOE contacts between the CH_3_-CH_3_ protons in the 4D spectrum. Numerous intra and inter domain NOE contacts between the methyl groups could be predicted from the crystal structure of the apo form of MALT1(PCASP-Ig3)_339–719_ (**Figure 5a**). The predicted interdomain NOEs (illustrated by black dashed line in **Figure 5b**) are not detected in the upper region part of the domain interface (Blue box in **Figure 5a**). Conversely, a chain of NOE intra domain contacts for the same methyl groups is clearly observed (red lines in **Figure 5b**). This specific NOE pattern, where contacts exist within the domains but are absent between them, aligns well with our results on the rotational diffusion tensors of the PCASP and Ig3 domains, and the conclusion of their semi-independent motions in solution. On the other hand, a different NOE pattern is also evident in the lower-middle region of the domain interface (Green box in **Figure 5a**). Notably, strong NOEs are observed between the methyl groups in both PCASP and Ig3, both within and between these domains (**Figure 5c**). This finding suggests that dynamics between these two domains is restricted in this particular patch. Further down at the bottom, the α1 helix followed by the loop connecting the two domains exhibit high structural flexibility as evident from our NMR results (**Figures 2b, 3c, 3d, and 6b**), as well as the missing electron densities and the high B-factors observed in the previously determined crystal structure of the apo form of MALT1(PCASP-Ig3)_339–719_ (PDB code 3V55). In summary, our results lead us to conclude that the lower-middle region of MALT1(PCASP-Ig3)_339–719_ may serve as a pivotal point for semi-independent movements of the two domains.

**Figure 5.**
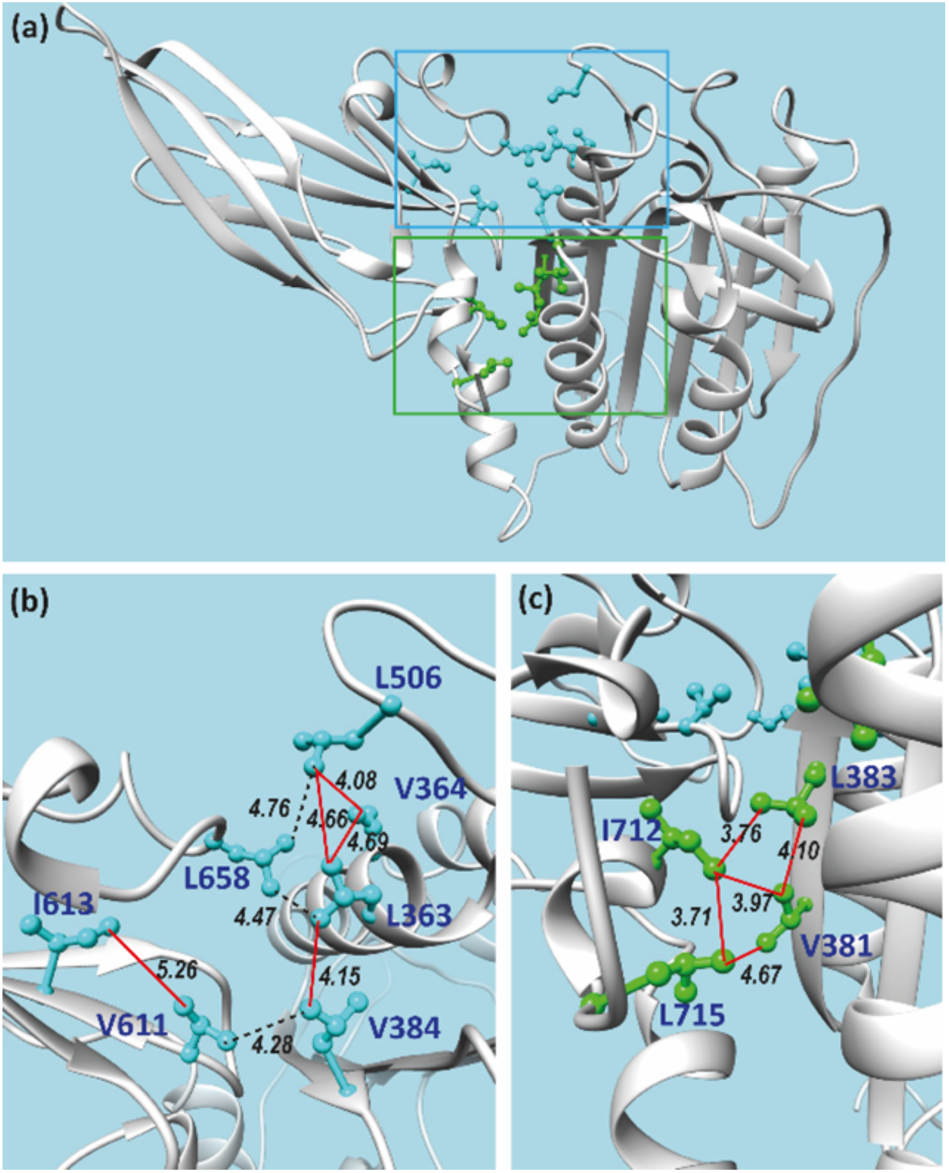
NOE contacts between the side chain methyl groups of MALT1(PCASP-Ig3)_339–719_ at the PCASP and Ig3 domains interface. **(a)** Overview of the side chains with methyl groups. **(b)** and **(c)** Zoomed areas in panel (a) are indicated as blue and green boxes, respectively. Based on the crystal structure of MALT1(PCASP-Ig3)_339–719_ in its apo form, a set of NOEs between the methyl groups were expected in our 4D NOESY experiments. For these, red and dashed black lines mark observed and unobserved NOEs, respectively. Distances (in Å) between the carbons of the methyl groups are indicated. The labels and side chains of the methyl-containing amino acids are also shown.

### 3.5 Dynamics analyses using AlphaFold, NMA, and MD

Although several important static structural snapshots of MALT1 have been obtained through X-ray crystallography studies, an unambiguous understanding of allosteric interactions and their relation(s) to the activity regulation of MALT1 remains challenging. The present study aims to shed light on these complex interdomain relations. Our experimental data revealed unambiguous fast dynamics between PCAP and Ig3 domains, suggesting the existence of multiple conformational substates.

#### 3.5.1 Quality assessment and dynamics of molecular models of MALT1(PCASP-Ig3)_339–719_ predicted by AlphaFold

We generated 2000 molecular structural models of MALT1(PCASP-Ig3)_339–719_ using AlphaFold. AF assesses its prediction confidence through two main scores; i) the predicted template modelling score (pTM) (Zhang Y 2004) (Jumper *et al*., 2021) and ii) the per-residue model confidence score (pLDDT) (Jumper *et al*., 2021, Mariani *et al*., 2013). The pLDDT score provides a per-residue estimate of model accuracy, ranging from 0 (low confidence) to 100 (high confidence). Notably, all 2000 structures predicted by AF consistently displayed high pTM scores, ranging from 0.89 to 0.92. As illustrated in **Figure 6a**, the pLDDT scores were also high throughout the structures of each created model of MALT1(PCASP-Ig3)_339–719_. However, specific regions of the protein exhibited relatively lower scores as highlighted in yellow in **Figure 6**. Recently a direct correlation was established between protein pLDDT and order parameters S^2^ characterising intramolecular dynamics of protein backbone (Ma *et al*., 2023). Accordingly, the regions with low pLDDT scores, such as the α1 helix of the Ig3 domain formed between residues 569 and 583, the loop in the active centre ranging between residues 496 and 509, and the region comprising residues 640-674 in MALT1(PCASP-Ig3)_339–719_, should also have low order parameters S^2^, indicating their high intramolecular dynamics. Importantly, as demonstrated in this study (**Figures 3c and 3d**), the same regions in MALT1(PCASP-Ig3)_339–719_ displayed low values for the local correlation times *τ*_C,i_ and the ^15^N-(^1^H)NOEs values, thus experimentally indicating that the dynamics are within the ps-ns time scale. Furthermore, the B-factors derived from the crystal structure of MALT1(PCASP-Ig3)_339–719_ (**Figure 6b**) demonstrated notable correlation with the pLDDT scores. The lowest B-factors aligned well with conformational heterogeneity, specifically in the third loop within the active centre (residues 496-509) and the region comprising residues 640-674. The α1 helix in Ig3 appeared somewhat better defined and rigid in the crystal structure, which does not correlate with our relaxation and AF data. This may be due to the stabilisation of this region by crystal packing. Indeed, the same region is poorly defined in other crystal structures of MALT1(PCASP-Ig3)_339–719_ with different space groups (PDB codes 3UOA and 3UO8) .

**Figure 6.**
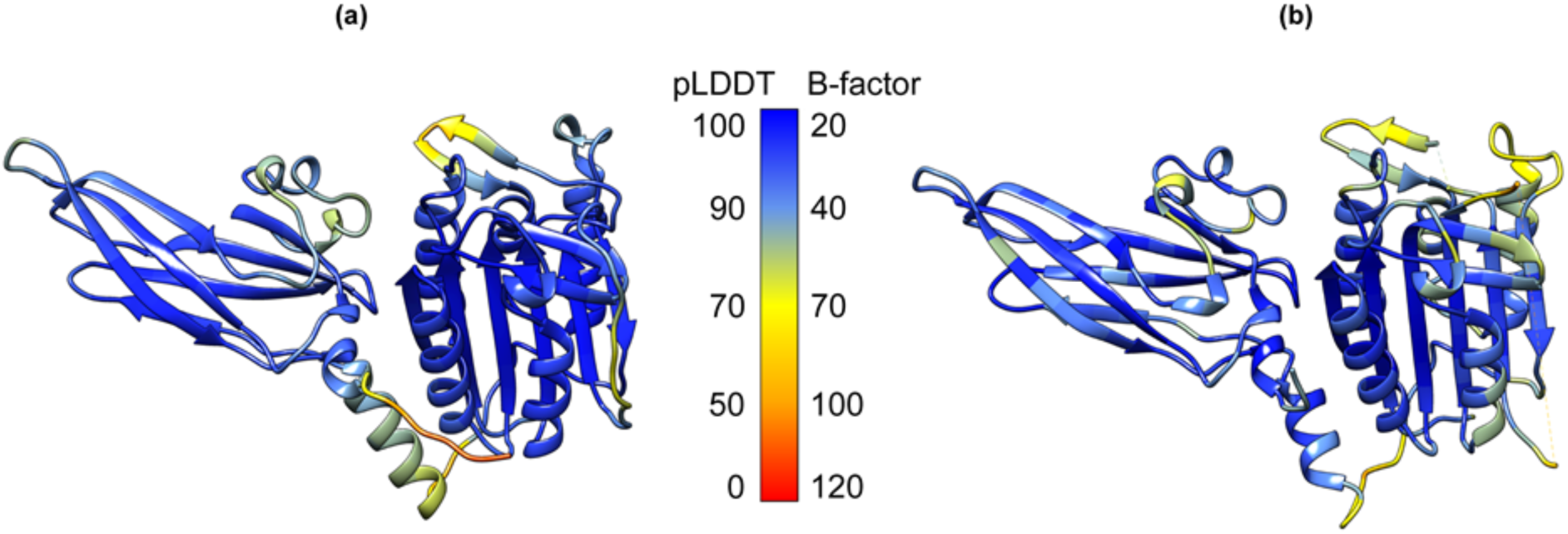
Local confidence of the MALT1(PCASP-Ig3)_339–719_ structures. **(a)** Per-residue AF confidence scores (pLDDT) calculated as a mean over all the 2000 AF structural models are mapped on the mean structure AF ensemble of MALT1(PCASP-Ig3)_339–719_. The low pLDDT scores 0-50 indicate conformation heterogeneity. **(b)** B-factor values ranging from 20 to 120 are mapped on the crystal structure of the apo form of MALT1(PCASP-Ig3)_339–719_ (PDB code 3V55).

#### 3.5.2 PCA analysis of the AlphaFold ensemble of structures

To place the predictions by AF of conformational variance of MALT1(PCASP-Ig3)_339–719_ in context with the experimentally obtained dynamic data within this study, we utilised principal component analysis. PCA effectively reduces the multidimensional space to a smaller, representative space that emphasises the primary conformational heterogeneity and motions. We initially conducted PCA (denoted as PCAa) on all 2000 structures predicted by AF encompassing the backbone N, Cα, C, and O atoms of protein residues 344 to 717. The first two principal components (PC1, and PC2) captured 48% and 17% of the structural variations among the AF models, respectively. The PC1/PC2 scores for individual structures from the ensemble shown in **Figure 7a**, revealed that the predicted conformations do not follow a simple interpolation between two end states, but instead, form a conformationally diverse ensemble of models. To reveal heterogeneity and cooperativity within the conformation ensembles of MALT1(PCASP-Ig3)_339–719_ captured by the PC1 and PC2 components in PCAa, we created a visual representation of the PCA loadings by superposing two structures representing the beginning and the end of the structural variations identified in PC1 and PC2, respectively (**Figures 7b and 7c**). Following the direction of coordinated variation between Ig3 and the α1 helix of MALT1(PCASP-Ig3)_339–719_ across the PC1 and PC2 components, it became evident that conformational changes in Ig3 are highly intricate (**Figure 7b**). Notably, a significant correlations unfolded within the PC1 component including bending of the Ig3 domain and displacement of the α1 helix between the PCASP and Ig3 domains (**Figure 7b**). These observations align well with our NMR data, highlighting the semi-independent movements of Ig3 and PCASP, both pivoting around a crucial point in the lower-middle region of their interface. It should also be noted that the PC2 component indicates that the Ig3 domain partially rotates around an internal axis (**Figure 7c**).

**Figure 7.**
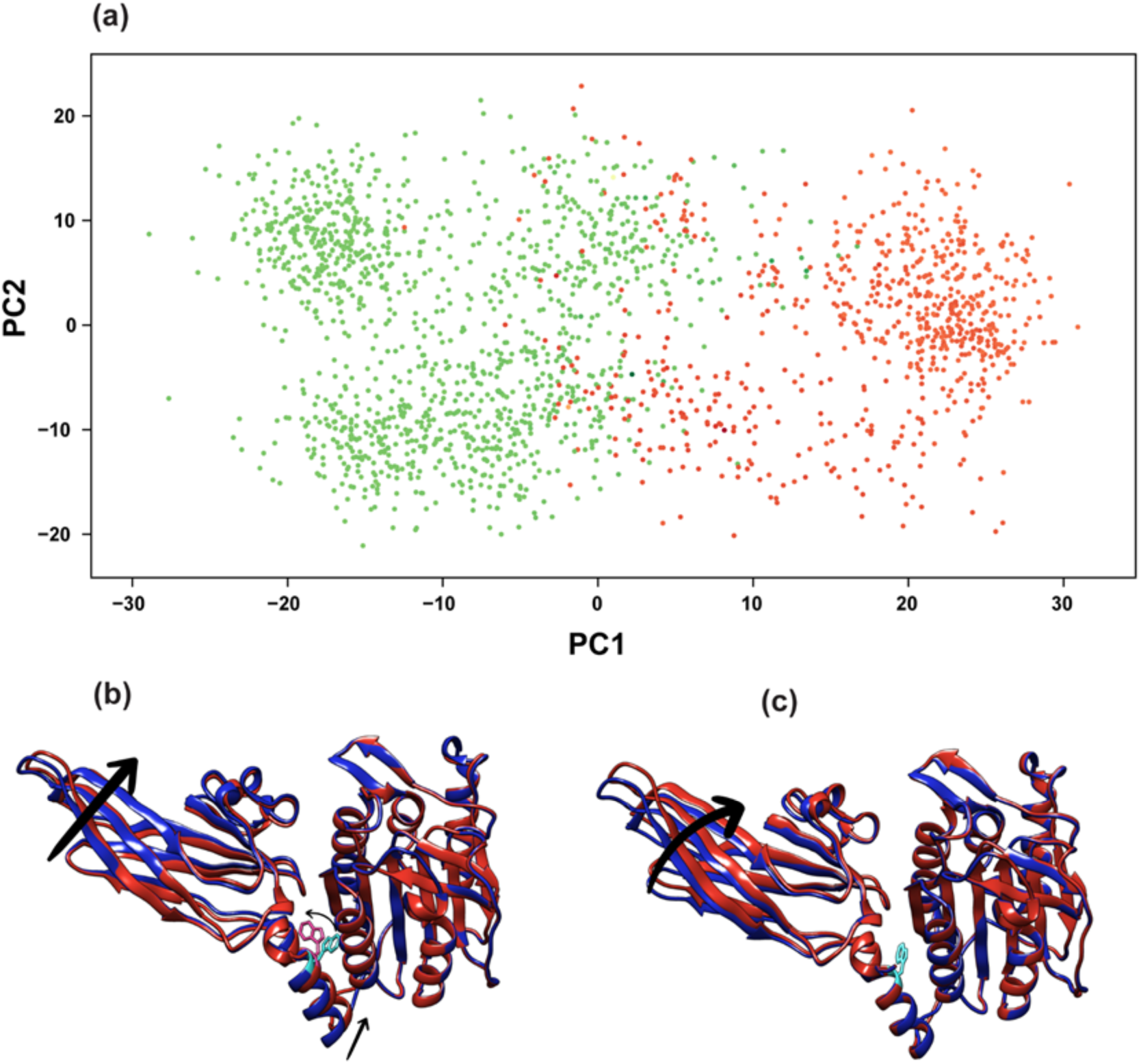
Principal component analysis performed on backbone nuclei (PCAa) of MALT1(PCASP-Ig3)_339– 719_ models generated using AlphaFold indicates semi-independent movements of both domains pivoting around the tryptophan residue W580. **(a)** Scatter plots depict 2000 distinct conformations in the PC1 and PC2 frames, revealing the conformational heterogeneity within AF models. Conformations with outward-facing (OF) and inward-facing (IF) W580 are represented in green and orange, respectively. **(b)** and **(c)** The superposition of the two structures shown in red and blue captures the two extremes of the structural variations in the conformation ensemble along the PC1 **(b)** and PC2 **(c)** components, respectively. Arrows indicate the direction of coordinated variation between Ig3, the α1 helix, and residue W580 in MALT1(PCASP-Ig3)_339–719_.

An intriguing yet unexpected discovery in our PCAa analysis of the AF dataset’s PC1 component is the conformational flip of the aromatic ring of residue W580 over the χ2 angle. Conformations with outward-facing (OF) and inward-facing (IF) orientations of residue W580 were identified, coloured in green and orange respectively, and clustered into two distinct families (**Figures 7a and 7c**). Notably, the conformational flip observed for W580 is strongly correlated with the PC1 score and hence with the conformation of the α1 helix.

Next, to further explore the potential structural rearrangements within the conformational ensembles of the Ig3 domain in the AF molecular models, induced by variations in the conformation of the aromatic ring of residue W580, we performed an additional PCA analysis, denoted as PCAb (PC1 74% and PC2 13%). This analysis presented in **Figure 8** focused on the two short stretches of residues 579-584 and 652-659 in MALT1(PCASP-Ig3)_339–719_, encompassing both aromatic rings of W580 and Y657. The primary and pivotal observation derived from these analyses was that when the aromatic ring of W580 flips out of the allosteric pocket, taking an outward-facing (OF) orientation with a χ2 angle of -154°, the aromatic ring of residue Y657 can adopt either of two χ1 angles, ranging from -92.8° to 57.3°. These values correspond to conformational states I and II (**Figure 8b**). In contrast, when the aromatic ring of W580 flips towards the allosteric pocket with a χ2 angle of 150.4°, the aromatic ring of Y657 consistently maintains a dominant position with a χ1 value of -92.8°, turned away from the third loop of the active site. This corresponds to conformational states III and IV, as presented in **Figure 8b**. The identified cluster populations are distributed as 55.5%, 8.3%, 35.0%, and 1.1% for states I, II, III, and IV, respectively (**Figure 8b**). According to our findings based on the AF prediction models, an equilibrium could exist between the two main conformations taken by the aromatic residue W580. Nevertheless, models with the highest confidence, according to the pLDDT score of W580, were predominantly found in states I and II (**Figure 8a**), which led us to the conclusion that, based on AF predictions, a conformational ensemble with the side chain of W580 in an outward-facing (OF) orientation is more likely. Additionally, we repeated the sampling of predictions for MALT1(PCASP-Ig3)_339–719_ using alternative statistical methods RosettFold and ESMFold, which both provided results that closely resemble those obtained with the AF models (**Supplementary Figure S5**). This is illustrated by the distribution of the pLDDT scores depicted on the structures, and by the predicted conformations adopted by the side chains of the aromatic residues W580 and Y657. Notably, similar to the most populated AF conformation, both RosettFold and ESMFold structures have the side chain of W580 taking an outward-facing orientation.

**Figure 8.**
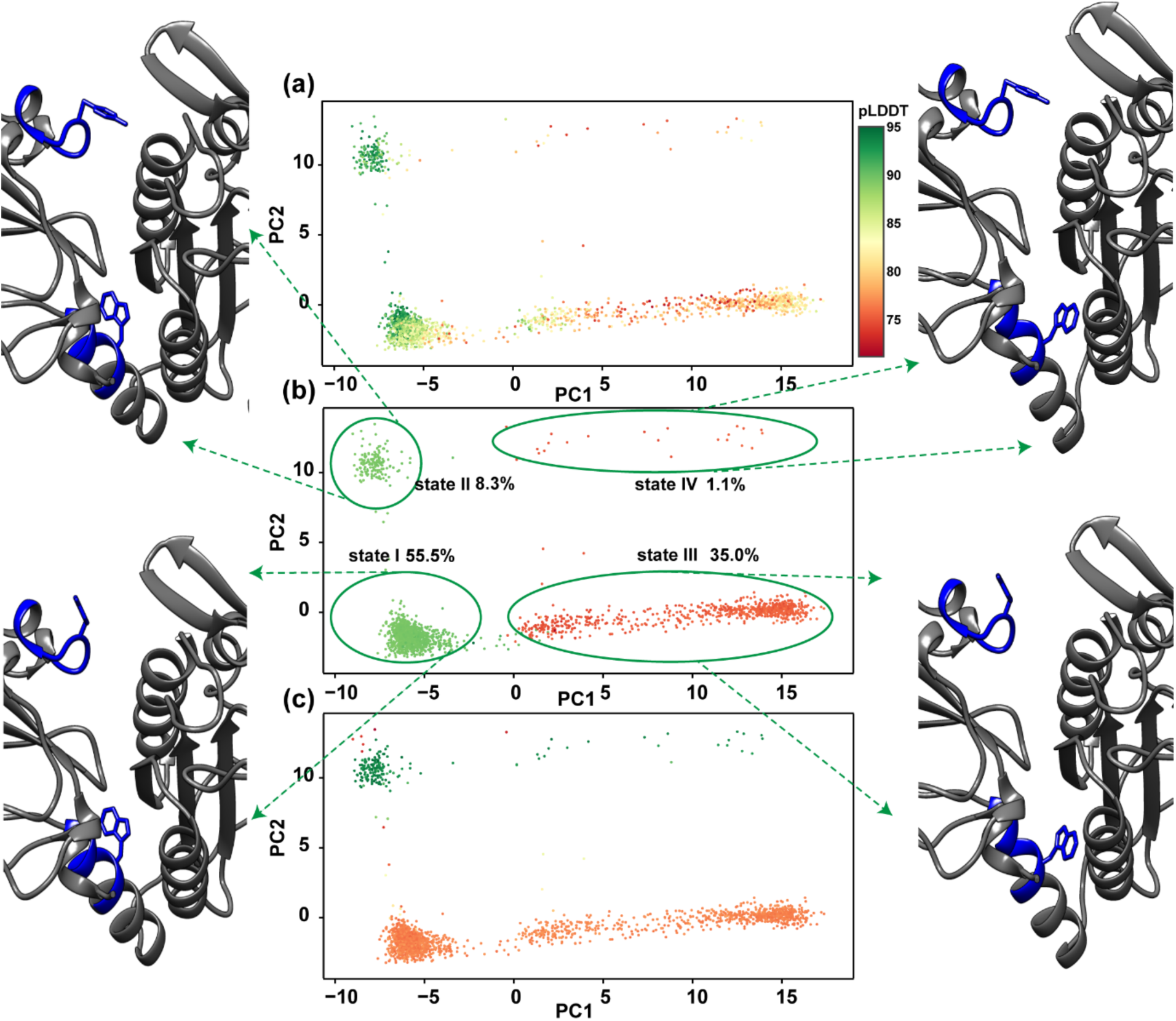
PCAb analysis of the MALT1(PCASP-Ig3)_339–719_ conformational ensemble predicted by AF identifies four distinct clusters. The alternative conformations of MALT1(PCASP-Ig3)_339–719_ modelled by AF are depicted in panels **(a)**, **(b)**, and **(c)**, featuring dimensionality reduction through the principal component analysis (PCAb), and illustrating the clustering of predicted conformations. In panel **(a)**, the model confidence is colour-coded using the pLDDT score for residue W580 in the range 71.29-95.13, with higher values indicated in green, signifying greater model confidence. As illustrated in panel **(b)**, dimensionality reduction revealed four distinct states (I-IV) based on the position of the aromatic rings of residues W580 and Y657, surrounded by green curves with estimated populations. Structural changes between the AF clusters are highlighted, and representative structures for each state are presented as green-coloured ribbons. The different orientations of the aromatic rings of W580 and Y657 are depicted in orange. Additionally, PCA models are coloured in panel **(b)** according to the χ2 angle score of W580, ranging from -154° (green) to 150.4° (orange). This clustering corresponds to either outward-facing (OF) or inward-facing (IF) W580 with respect to PCASP, respectively. In panel **(c)**, PCA models are coloured based on the χ1 angle score of residue Y657, ranging from -92.8° (green) to 57.3° (orange). This colouring separates the inward-facing (IF) and outward-facing (OF) orientations of the side chain of residue Y657, respectively.

#### 3.5.3 PCA analysis of MD and NMA structures

To investigate potential transitions between the alternative conformations of MALT1(PCASP-Ig3)_339– 719_, revealed in the AF ensemble of structures, we employed two computational approaches for modelling molecular dynamics (**Supplementary Figure S4**); an all-atom Molecular Dynamics with explicit solvent (MD) and Normal Mode Analysis (NMA). The methods explore local and global minima of conformation ensembles, depending on the force field used in the calculations. **Supplementary Figures S4a, S4b and S4c** display the superpositions of two structures captured at the opposite ends of a structural span in all the conformation ensembles obtained from the first three NMA modes. Additionally, **Supplementary Figures S4d and S4e** depict the loadings of the PC1 and PC2 components of a PCA performed on the 2000 structures extracted from a 2μs MD trajectory. Visual inspection of **Supplementary Figure S4** highlights similarities between the conformational diversity exhibited by the Ig3 domain in the MD and NMA calculations, which is also well in line with the analyses of the AF conformational ensembles described here above. It is noteworthy that both NMA and MD do not display any significant diversity in the orientations of the α1 helix in MALT1(PCASP-Ig3)_339–719_. Accordingly, no coupling was observed between the positioning of the Ig3 domain and the orientation of the α1 helix. Furthermore, during the full MD trajectory, we were unable to detect any transition from the inward-facing (IF) orientation of the aromatic ring of W580, as identified in the initial structure, towards the outward-facing (OF) orientation preferred by AF, ESMFold2 and RoseTTAFold2. This may be explained by the intrinsically local conformational sampling in NMA and possibly insufficiently long runs of the MD traces to identify rare conformational transitions.

## 4.0 DISCUSSION

Over the last decade, MALT1 has gained significant attention in cancer research for its crucial role in regulating specific signalling pathways that influence the development and proliferation of T and B cells. Unlike many other signalling mediators, MALT1 does not only act as a major scaffold for the formation of the oligomeric BCL10-MALT1 filament complex (**Figure 1a**), but serves also as an essential cysteine protease through its PCASP domain. This dual functionality renders MALT1 an exceptionally attractive target for drug development. Recent advances in cryo-electron microscopy (cryo-EM) have enabled determination of the low-resolution structure of the BCL10-MALT1 filament complex (Schlauderer *et al*., 2018). However, discerning the PCASP domains within this filament structure has proven to be a challenge due to its intrinsic conformational dynamics. Furthermore, successful determination of the crystal structure of the active portion of MALT1, comprising only the PCASP and Ig3 domains, has allowed a first atomic visualization of the functional core of MALT1 (**Figure 1b**), which still retains its protease activity (Wiesmann *et al*., 2012). Notably, the crystal structure indicated the formation of a dimer of MALT1(PCASP-Ig3)_339–719_ in both its apo state and in complex with different ligands. However, to date, no three-dimensional structure of the MALT1(PCASP-Ig3)_339–719_ monomer has been reported, despite that it is the most prevalent form in solution in the absence of ligands or kosmotropic salts (Cabalzar *et al*., 2013).

The exploration of protein dynamics is essential for gaining insights into biological processes at an atomic and molecular level. Despite the extensive biochemical and structural research conducted on MALT1 over the last decade, an investigation of its dynamics has not been performed. In this study, we focused on the dynamics of MALT1(PCASP-Ig3)_339–719_ in order to assess and better understand the complex interplay between the monomeric and dimeric states, domain interactions, conformational space sampling. Our investigation began by addressing whether the apo form of MALT1(PCASP-Ig3)_339–719_ in solution exists predominantly as a monomer or as an equilibrium between monomeric and dimeric states. Using classical gel-filtration, we isolated the monomer form of MALT1(PCASP-Ig3)_339–719_, which was subsequently employed in our solution-state NMR experiments. Our NMR results and analyses confirmed that MALT1(PCASP-Ig3)_339–719_ remains primarily monomeric in our experimental settings.

According to the crystal structure of the apo form of MALT1(PCASP-Ig3)_339–719_, the Ig3 domain interacts tightly with the PCASP domain through the formation of several hydrophobic contacts, which are affected upon substrate binding. Residue W580 plays a crucial role in maintaining this interaction, contributing to a stable protein fold by binding to a hydrophobic pocket in PCASP and bridging it to Ig3. In contrast, our present NMR-based dynamic study of the monomeric form of MALT1(PCASP-Ig3)_339–719_ reveals a captivating complexity in domain interactions. Instead of forming a tight complex or being fully independent, the PASC and Ig3 domains exhibit a dynamic, semi-independent movement around a pivotal point.

Utilising AlphaFold and PCA, our study provides insights into the conformational space sampling of the PCASP and Ig3 domains detected by our NMR relaxation experiments. The bend and twist movement of the Ig3 domain (**Figure 7**) seems to be coupled to the orientation taken by the α1 helix connecting the PCASP and the Ig3 domains. The AF structural ensemble that we obtained in this study is generally well in line with the structural variations observed in the multiple crystal structures of MALT1(PCASP-Ig3)_339–719_ that have been previously determined both in its apo- and its ligand-bound forms, including the span of relative orientations of PCASP and Ig3, as well as the multiple orientations taken by the α1 helix of the Ig3 domain.

As a control for the results predicted by AF, we produced structural models of MALT1(PCASP-Ig3)_339– 719_ using the two alternative AI-driven approaches ESMFold2 (Lin *et al*., 2023) and RoseTTAFold2 (Baek *et al*., 2023) that, similar to AF, exploit multiple sequence alignments (MSA). Despite their functional similarity, it is crucial to note significant conceptual and architectural differences in these three programs, as previously highlighted (Baek *et al*., 2023, Liu *et al*., 2022). Our results revealed that both RoseTTAFold2 and ESMFold2 successfully reproduced the structural models obtained by AF. The quality of the predicted models was assessed using the per-residue RoseTTAFold2 and ESMFold2 confidence scores (pLDDT) (**Supplementary Figure S5**), which show clear similarities with the AF pLDDT scores (**Figure 6**) and more specifically in the connecting α1 helix of Ig3. Notably, both RoseTTAFold2 and ESMFold2 models show the outward orientation of the W580 side chain, which is also the most dominant conformation in the AF conformational ensemble analysis (Figure 8). The consistency in the MALT1(PCASP-Ig3)_339–719_ structures obtained by AF, RoseTTAFold2, and ESMFold2 modelling methods despite differences in their underling algorithms increase confidence in our results.

A new intriguing aspect emerged in the detailed analysis of the AF ensemble obtained for the apo form of MALT1(PCASP-Ig3)_339–719_. Here, the aromatic ring of residue W580 adopted both orientations found in previously determined crystal structures, with an inward conformation, where the side chain ring resides within the hydrophobic groove localized between the Ig3 and PCASP domains, often also called the allosteric pocket, and the outward-projecting conformation. However, somewhat surprisingly, the most represented of the two conformation taken by W580 was the one with its aromatic ring projecting out (**Figure 8b**), which was previously found in crystal structures of MALT1(PCASP-Ig3)_339–719_ in complex with allosteric ligands occupying the allosteric site. Conversely, in the apo form of MALT1(PCASP-Ig3)_339–719_, W580 adopts the inward-looking orientation and occupies the allosteric pocket. Based on the static crystallographic data, it is commonly accepted that the aromatic ring of W580 is pushed out from its position in the hydrophobic groove into a solvent-exposed environment upon binding of an allosteric inhibitor by an induced fit mechanism, followed by a substantial displacement of the α1 helix of Ig3 (Schlauderer *et al*., 2013).

Our current NMR analysis of the apo form of MALT1(PCASP-Ig3)_339–719_ allowed us to unambiguously identify in solution only one of the AF-predicted forms. To determine the dominant form of W580 in the apo form of MALT1(PCASP-Ig3)_339–719_ in solution, we utilised the fact that in the conformation where the aromatic ring W580 resides in the allosteric pocket (**Figure 8b, states III and IV**), one of the methyl groups in each of V381 and V401 residues are positioned almost exactly above and below, respectively, relative to the aromatic ring of W580. In this configuration, the protons of these methyl groups experience a significant upfield chemical shift due to the aromatic ring effect of the side chain of W580 (Gunther, 1987). Indeed, the chemical shifts of the protons of V381Cγ2 and L401Cδ1 were observed in the ^1^H-^13^C HSQC spectrum of MALT1(PCASP-Ig3)_339–719_ at very high upfield positions, - 0.577 and -0.505 ppm, respectively. This contrasts with V381Cγ1 and L401Cδ2, where the shielding effect of the aromatic ring is minimal, at 0.467 and 0.414 ppm, respectively. Surprisedly, the dominant form of MALT1(PCASP-Ig3)_339–719_ existing in our experiments in solution with inward orientation of W580 corresponds to the least populated cluster in the AF ensemble. Does this mean that the AF, ESMFold2, and RoseTTAFold2 algorithms, which are based on statistics and/or artificial intelligence (AI) combined with genome sequence analysis, provide erroneous molecular models for MALT1(PCASP-Ig3)_339–719_ by preferring the conformation of the side chain of W580 localized outside the allosteric hydrophobic grove?

There is credible criticism in recent literature (Nussinov *et al*., 2023, Carugo, 2023, Terwilliger *et al*., 2023) regarding the ability of AF to predict correct partitioning in structural ensembles. It was argued that one can only give full credence to conformations that have been experimentally validated and dismiss those that have not undergone validation. Consequently, although the outward-facing conformation of the side chain of W580 configuration have been observed in the experimental crystal structures of ligated MALT1(PCASP-Ig3)_339–719_, its dominance of the AF ensemble appearance in the apo form of the protein may be seen as a bias in the AF model. On the other hand, despite acknowledging limitations in the AF conformation predictions, a number of interesting results were recently presented (Wayment-Steele *et al*., 2023) that encourage further studies on AF structural ensembles. It is essential to acknowledge that the AF algorithm lacks specific information about the environment conditions for the conducted experiments. One can argue that modelling using statistics from three-dimensional structures found in the Protein Data Bank (PDB) and amino acid sequences (i.e. MSA) could instead reflect the likeliest PDB structural motives and cellular context and functional relevance in evolution. Specifically, while the distance constrains derived from the MSA reflect evolutionary and thus functionally important structures, statistics derived from PDB structures reflects likely structural features and contexts, e.g. interactions with other molecules, found in the PDB. To this end, it is not surprising that all our AF molecular models display PCASP in its functional enzymatically active configuration, although this state was crystallised only with ligands bound at the active site (Wiesmann *et al*., 2012), but not for the apo-form of the protein. Furthermore, even when in an inactive resting state, MALT1 can be activated by a substrate or changing environmental conditions (Wiesmann *et al*., 2012), such as in the presence of kosmotropic salts (Coornaert *et al*., 2008). This underscores the importance of recognizing the potential for shifts in populations within the conformation ensemble in solution. Both AF-predicted conformations of W580 may have a sizable representation in solution, even though the facing-outward conformation of W580 was not observed and thus low populated under the specific conditions of our experiments. Altogether, our results allows us to extend the structural perspectives that emerged from previous X-ray studies of MALT1(PCASP-Ig3)_339–719_, towards acknowledging the existence of diverse conformational ensembles in solution and underscoring the necessity of considering solution dynamics when designing function modulating ligands for MALT1. More specifically, based on our present results, we argue that binding of allosteric drug involves a conformational selection that repartitions the conformational ensemble with stabilization of the flipped-out configuration rather than causing W580 to switch (Quancard *et al*., 2019) its conformational positions to vacate the groove space for the inhibitor. Finally, and perhaps the most disputable and intriguing possibility would be to accept that the AF partitioning of the conformational states reflects a real albeit hitherto unknown environment situation *in vivo*. Applied to the MALT1(PCASP-Ig3)_339–719_ system, AF strongly prefers molecular models where W580 turns away from the hydrophobic groove (**Figure 8a**). This conformation is usually observed *in vitro* at experimental conditions when the proteolytic activity is inhibited by an allosteric ligand. Noting that the AF algorithm runs without the structural templates and lacks knowledge about the MALT1(PCASP-Ig3)_339–719_ crystal structures with specific unnatural inhibitors, we could hypothesise that the predicted AF dominant conformation prevails *in vivo*, may be stabilized by an as-yet-unknown natural inhibitor/factor. This may be used by the cell to limit the function of MALT1 solely to its scaffolding role. Indeed, recent *in vitro* study involving the mutation of W580 to a serine (Quancard *et al*., 2019) demonstrated that upon binding allosteric inhibitors to MALT1-W580S, the protein scaffold functions can be restored through the rescue of canonical NF-κB and c-Jun N-terminal kinase (JNK) signalling.

The PCA analysis (PCAb) of AF molecular models conducted here on the two short stretches of residues 579-584 and 652-659 in MALT1(PCASP-Ig3)_339–719_ allowed us to examine further potential structural rearrangements within the conformational ensemble of the Ig3 domain. These regions encompass both the aromatic ring of residues W580 and Y657. Here, the key observation is that when the aromatic ring of W580 faces outward from the allosteric pocket, the aromatic ring of Y657 can adopt either of two χ1 angles, ranging between -92.8° and 57.3°, representing conformational states I and II (**Figure 8b**). Conversely, when the aromatic ring of W580 is oriented towards the allosteric pocket, the aromatic ring of Y657 consistently maintains a dominant position with a χ1 value of -92.8°, turned away from the third loop of the active site. This corresponds to conformational states III and IV (**Figure 8b**).

It is also important to highlight a recently reported alternative allosteric pathway for restoring and regulation of the protease function of MALT1 through monoubiquitination (Schairer *et al*., 2020). The study suggests that Ubq attachment to residue K644 in the Ig3 domain induces conformational changes in the Ig3-protease interface, ultimately enhancing MALT1 activation while pointing to the importance of Y657 as a key mediator in signalling between Ig3 and protease domains. It is noteworthy that in the AF-predicted state II (**Figure 8b**), the aromatic Y657 potentially impacts on hydrophobic inter-domain interactions, resembling the proposed structural alteration seemingly provoked by covalent attachment of Ubq to K644 (Schairer *et al*., 2020). Additionally, cluster II (**Figure 8b**) features orientation of the Y657 aromatic ring as in the crystal structure where the protein is bound the substrate analogue z-VRPR-fmk at the active site. Based on this similarity, we argue that the conformational equilibrium between states I and II for MALT1 may shift through monoubiquitination, leading to the activation of MALT1 protease function. It is noteworthy that only states III and IV, characterized by the inward configuration of W580, are observed in the NMR experiments of MALT1(PCASP-Ig3)_339–719_ in solution. Furthermore, the active form IV, in which residue Y657 interacts with the active site, has a cluster population of only 1.1% as predicted by AF. Altogether, these observations may explain the significantly lower activity of the monomer in solution.

Next, we made an additional effort to understand the movements responsible for potential transitions between different conformations of the aromatic amino acids W580 and Y657 in MALT1(PCASP-Ig3)_339–719_, as identified in the AF models. Using a structure from state III family as a starting point (**Figure 8**), we conducted long-sampling conformation trajectories through a 3μs Molecular Dynamics (MD) simulation. This MD simulation successfully reproduced the substantial flexibility of the Ig3 domain and predicted the conformational flips of the aromatic ring of Y657, as observed in the AF modelling. However, the most crucial finding was that the MD simulation did not indicate any conformational changes for the side chain of W580 in the allosteric pocket, as seen in the State I or II families from the AF modelling (**Figure 8b**). Similar results were obtained in Normal Mode Analysis (NMA). Additionally, a PCA of the 2000 structural ensemble extracted from the MD trajectory shows neither diversity in the conformation ensemble between the α1 -helix and the PCASP domain of MALT1(PCASP-Ig3)_339–719_ nor the conformational flips of the aromatic ring W580 out of the allosteric pocket. This is not surprising since the PCA analysis the AF ensemble indicated the coupling of these two movements. Two main factors may explain the divergence between the MD and AF results. First, the MD force field and AF modelling are not precise. Secondly, the transition from the global energy minima of the W580 conformation likely requires even more extended MD trajectory time. This is supported by the measured higher value of the exchange rate Rex in the interdomain linker, including in the α1 -helix, indicating the presence of slow exchange. To further investigate the possibility of slow exchange or transitions between W580 outward and inward orientations future measurements of relaxation dispersion on the aromatic ring W580 and methyl protons of the V381 and L401 amino acids are needed. These measurements could provide insights into the dynamics and transitions of W580 between different conformations.

## 5.0 CONCLUSION

We have performed the first comprehensive NMR study of MALT1 dynamic behaviour in solution. Measurements and analyses of the ^15^N-backbone *R*_1_- and *R*_2_-relaxation, as well as the CCR rates of MALT1(PCASP-Ig3)_339–719_ and the mutated E549A variant, revealed a complex picture of motions including overall rotational diffusion, inter-domain motion and local internal dynamics. Our NMR relaxation results confirm that MALT1(PCASP-Ig3)_339–719_ is a monomer in our experimental conditions. Furthermore, the PASC and Ig3 domains do not form a tight complex but instead exhibit semi-independent movements around a pivotal point. To model the movement between PASC and Ig3 domains and characterise the conformational ensemble for the MALT1(PCASP-Ig3)_339–719_, we employed AlphaFold, followed by a mechanistic rationalisation of the conformational space using principal component analysis. The AF modelling unveils significant correlations between the interdomain movements, bending of the Ig3 domain, and the displacement of the α1-helix located between the PCASP and Ig3 domains. Furthermore, displacements of the α1-helix in the AlphaFold ensemble of structures are highly correlated with the orientation of the aromatic ring of W580 located in the interdomain pivot region. Then, the orientation of W580 is a marker of the accessible states of the Y657 located further away in the interdomain interface close to the PCASP catalytic site, which is most probably consequential for the allosteric regulation of the protein catalytic activity.

According to the AF structural ensemble, an equilibrium could exist, probably even in the apo form of MALT1(PCASP-Ig3)_339–719_, between conformations with two orientations of the aromatic ring in W580, notably including the outwardly facing conformation previously experimentally observed in MALT1(PCASP-Ig3)_339–719_ complexes with the allosteric inhibitors occupying the “native” position of the W580 side chain. The outward conformations were also found in models provided by alternative AI-based methods RosettFold and ESMFold. To further explore the putative potential transitions between the alternative conformations of MALT1(PCASP-Ig3)_339–719_ predicted by AF models, we performed a 2 μs MD simulation. The observed conformational diversity in the Ig3 domain aligns with data obtained in AF models, yet the transition of the aromatic ring of W580, as predicted in AF, is not detected within the MD trajectory. Finally, we discuss validity of the AF-predicted conformational ensembles and specifically the relevance and possible implications in the context of complex cellular environment of the preference of the outward facing W580 conformation preferred by AF, RoseTTAFold, and ESMFold without presence of the allosteric inhibitor.

In summary, our study significantly extends the understanding of MALT1 dynamics and may shed light on structural details and its functional and regulation. The revelations regarding domain interactions, conformational space sampling, and apo form dynamics pave the way for further exploration of MALT1’s role in cellular processes and the development of targeted therapeutic interventions. As we delve deeper into the intricate realm of protein dynamics, MALT1 emerges as a captivating case study, illustrating the dynamic nature of molecular systems in biological processes.

## Author contributions

JW performed the analysis of relaxation data and diffusion tensors. XH and RS have contributed with production of all necessary protein variants, including their purifications and the developing a construct of labelled MALT1 proteins. JW and TA wrote the original manuscript draft. TA, AA, TS and VO contributed with writing and finalizing of the manuscript. VO, DL contributed with NMR measurements and methodology. DL performed MD calculation and analysis. TA and PA performed the NMR studies and assignments of MALT1 using the ccpn program. TA performed the NOE analysis between methyl groups. TS analysed earlier crystallographic data. PA, TA, TS, AA, BW, CM, GK and VO conceptualised the project, supervised different parts of the project. ML, DM, and CM performed the AlphaFold and PCA analyses.

## Acknowledgements

The authors thank the Swedish NMR centre for access to the instruments and support.

This work was supported by the Swedish Foundation for Strategic Research grant ITM17-0218 to T.A and P.A., grant RSF 23-44-10021 to D.M.L, Swedish Cancer Society (21 1605 Pj 01 H), Cancer och Allergi Fonden (10399), and Swedish Research Council (2021-05061 and 2018-02874) to A.A. and 2019-03561, 2023-03485 to V.O. This study used NMRbox: National Center for Biomolecular NMR Data Processing and Analysis, a Biomedical Technology Research Resource (BTRR), which is supported by NIH grant P41GM111135 (NIGMS).

The AlphaFold analyses were done under the *BeyondFold* Technology Development Project at the Science for Life Laboratory (SciLifeLab). CM is financially supported by the Knut and Alice Wallenberg (KAW) Foundation as part of the National Bioinformatics Infrastructure Sweden at SciLifeLab. Some computations were performed at NSC BerzeLiUs provided by the National Academic Infrastructure for Supercomputing in Sweden (NAISS) and the Swedish National Infrastructure for Computing (SNIC), partially funded by the Swedish Research Council through grant agreements no. 2022-06725 and no. 2018-05973.

## Data availability

The ensemble of AF and MD structures and other information and data are available upon request from the authors.

## Supporting Information

**Figure S1.**
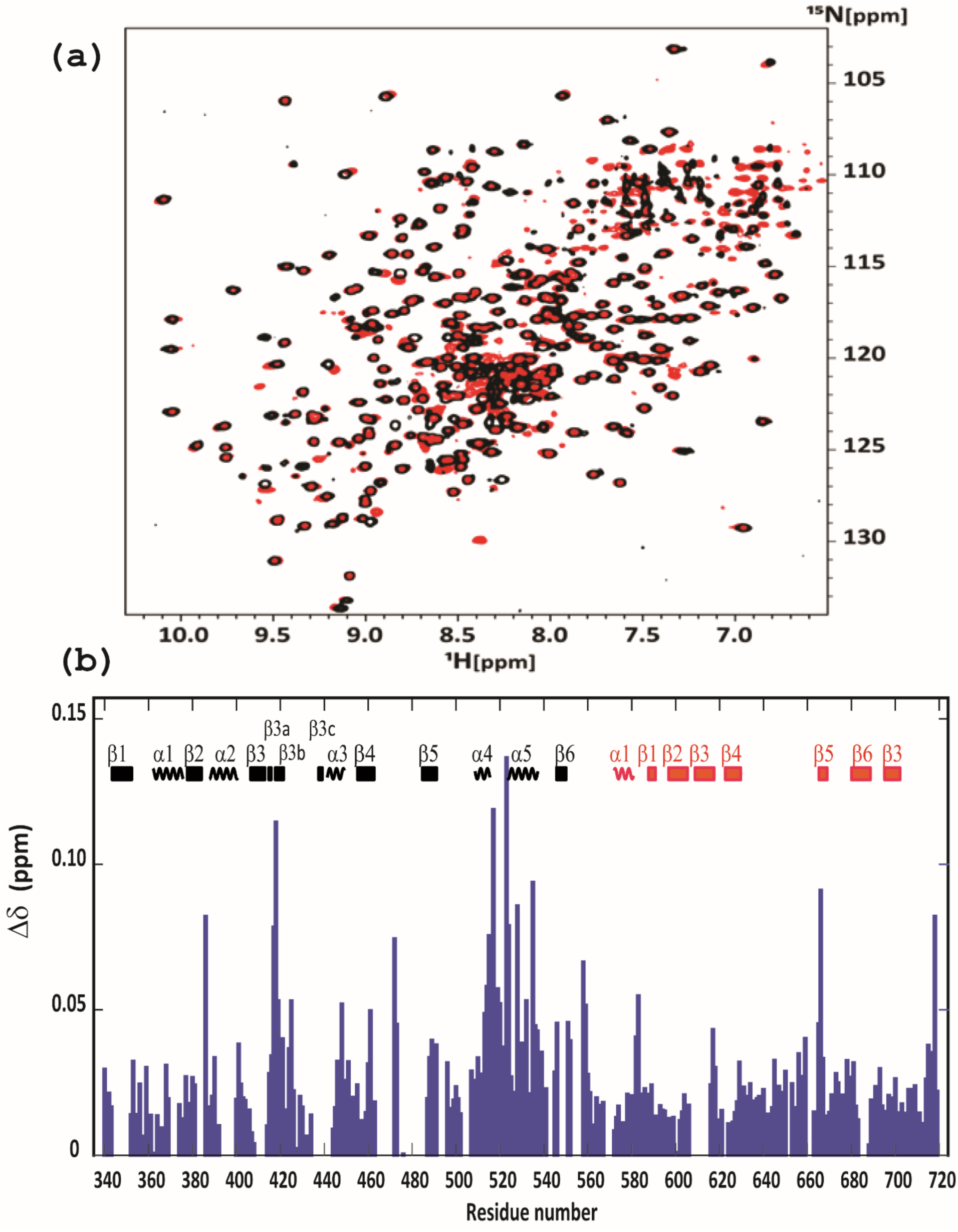
Chemical shift difference for the wild time and E549A mutant of MALT1(PCASP-Ig3)_339–719_. **(a)** The superposition of the TROSY ^1^H-^15^N spectra of the apo form of MALT1(PCASP-Ig3)_339–719_ (black) and its mutated variant E549A (red) recorded at 25°C on a 900MHz spectrometer. **(b)** Chemical shift perturbations for wild-type MALT1(PCASP-Ig3)_339–719_ (blue circle) and MALT1(PCASP-Ig3)_339–719_(E549A). Secondary structure elements are indicated and numbered for each domain at the top with PCASP in black and Ig3 in red.

**Figure S2.**
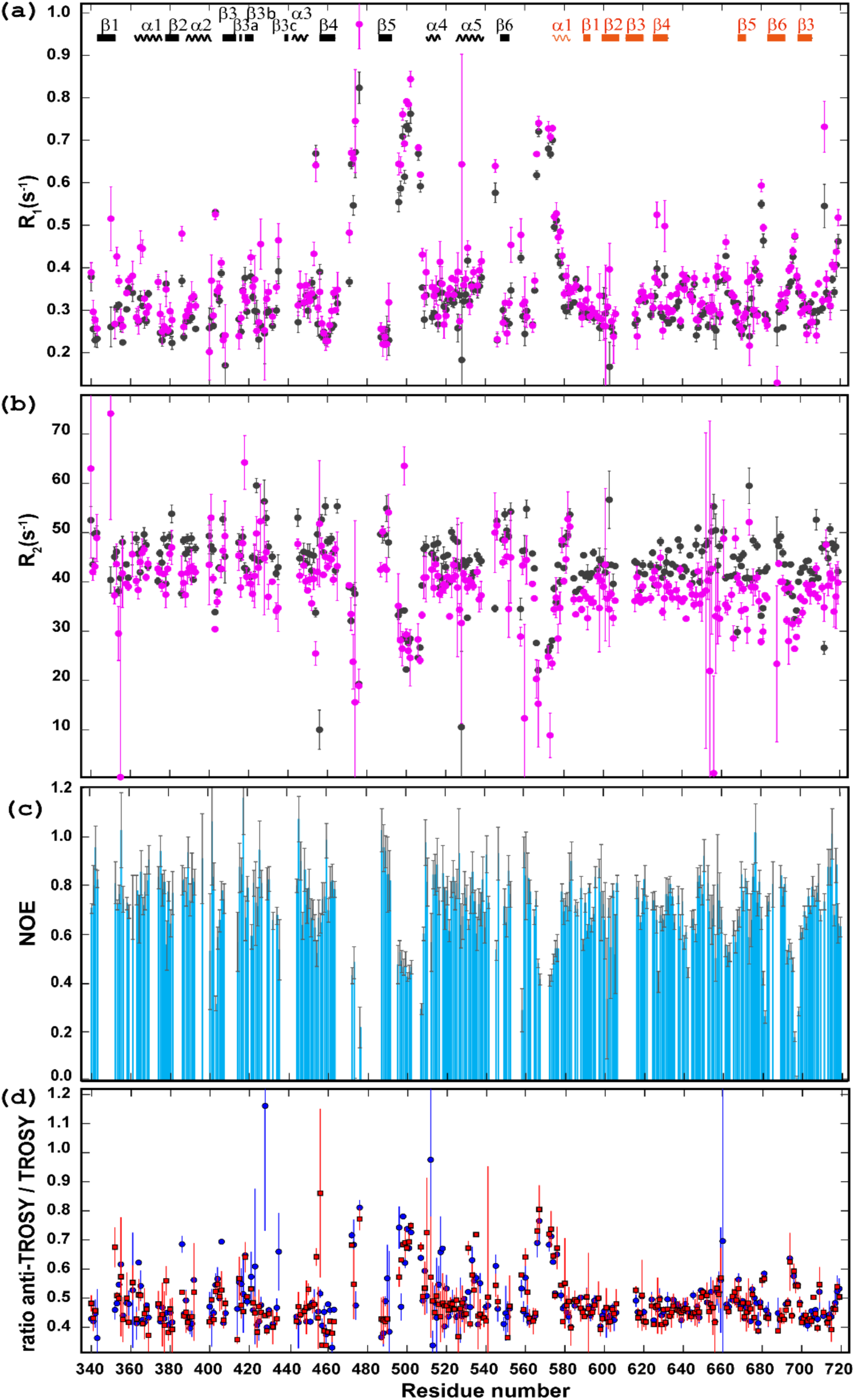
**(a)** ^15^N *R*_1_-relaxation rates and **(b)** ^15^N *R*_2_-relaxation rates for MALT1(PCASP-Ig3)_339–719_. Black dots represent 900 MHz data and magenta dots represent 800 MHz data. Secondary structure elements are indicated and numbered for each domain at the top. The start of the α1 helix at residue 574 marks the separation between the PCASP and Ig3 domains. Error bars show 1 SD. **(c)** ^15^N-(^1^H) NOE with error bars from 800 MHz. **(d)** ^15^N anti-TROSY/TROSY ratio for MALT1(PCASP-Ig3)_339–719_ (blue circle) and for MALT1(PCASP-Ig3)_339–719_(E549A) (red square). Data sets are from 900 MHz. Error bars show 1 SD calculated from the two intensities at t_CPMG_ = 0 for the anti-TROSY and the TROSY data sets. Secondary structure elements are indicated and numbered for each domain at the top with PCASP in black and Ig3 in red.

**Figure S3.**
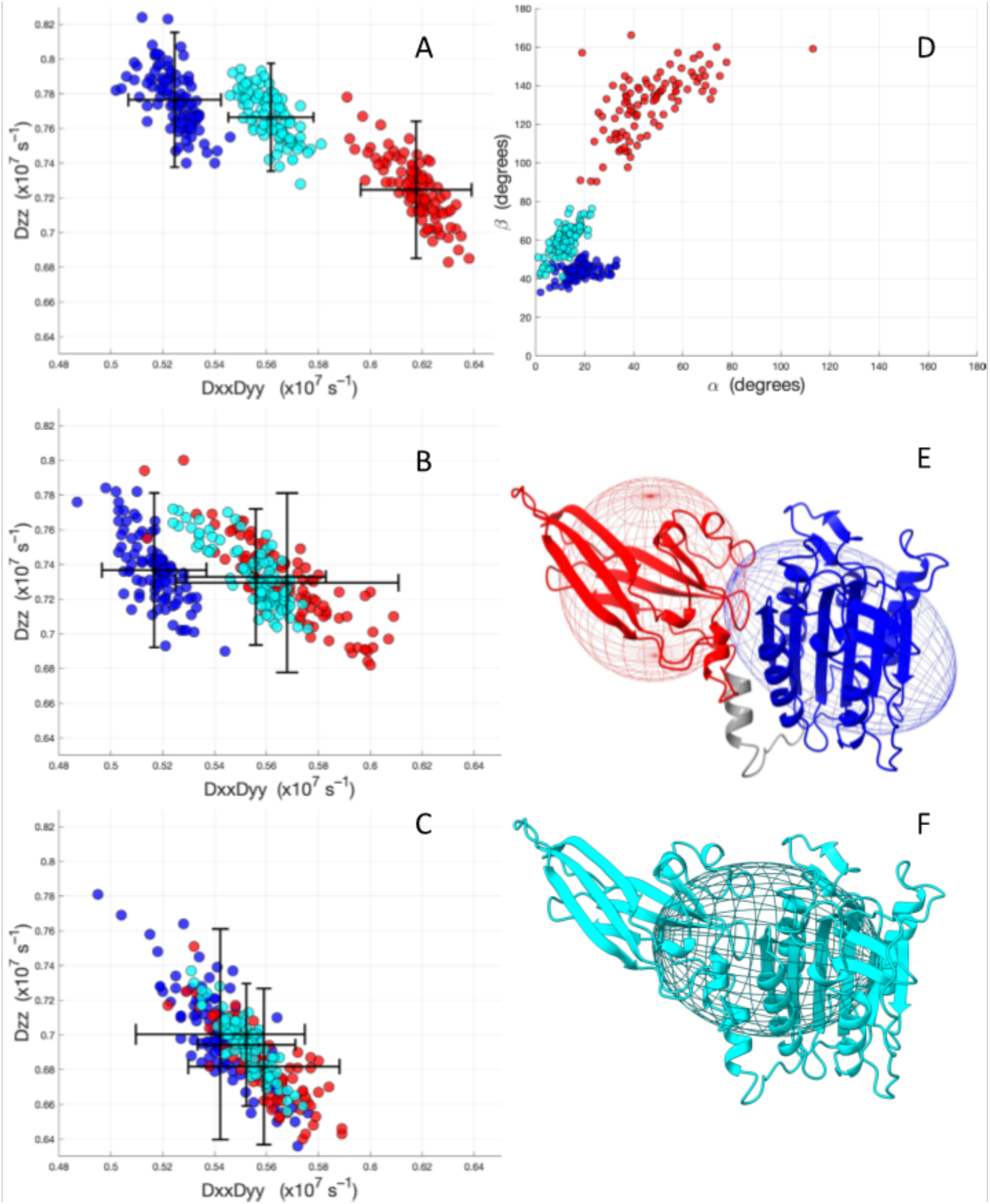
Diffusion tensor components D_ZZ_ and D_XX_D_YY_ from resampled ROTDIF optimisations. Each red dot corresponds to a tensor calculated for one resampling instance represent for Ig3, the blue dots represent the PCASP domain and cyan dots represent full MALT1(PCASP-Ig3)_339–719_ for **(a)** 900 MHz ^15^N *R*_2_/*R*_1_ data, **(b)** 800 MHz ^15^N *R*_2_/*R*_1_ data, **(c)** 900 MHz and cross-correlated transverse relaxation data. Error bars in (a)-(c) show one SD and are calculated taking the resampling fraction of *d=*20% into account. **(d)** α - and β -angles for the diffusion tensors for the 900 MHz ^15^N *R*_2_/*R*_1_ data set. **(e)** Ellipsoid representation of the diffusion tensors (900 MHz ^15^N *R*_2_/*R*_1_ data) mapped on the crystal structure of MALT1(PCASP-Ig3)_339–719_, with Ig3 depicted in red and PCASP in blue. The grey part, residue 563–582, comprising the loop and the α1 helix is excluded from the optimization in ROTDIF. See material and method section for the procedure of selection residues in the respective domains. The principal axis of the axially symmetric diffusion tensors crosses the poles of the ellipsoids. **(f)** The diffusion tensor calculated for the full protein.

**Figure S4.**
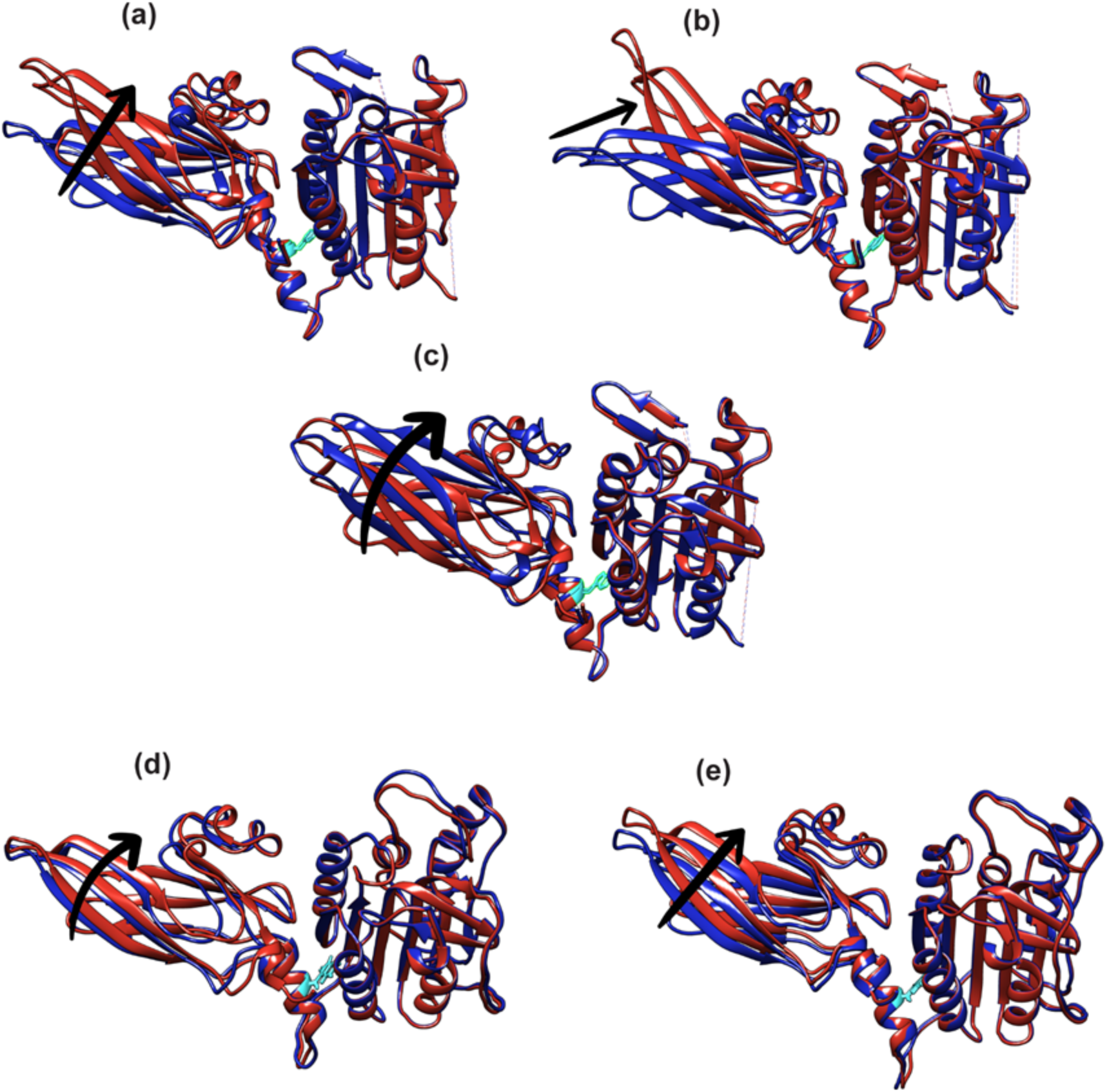
**(a)**, **(b)**, **(c)** Molecular models of MALT1(PCASP-Ig3)_339–719_ predicted using the NMA showing modes 7, 8 and 9, respectively. **(d)**, **(e)** PCA 1 and PCA 2 modes, respectively, obtained on confirmation ensemble of 2000 structures extracted from MD trajectory of 2μs after 1us equilibration.

**Figure S5.**
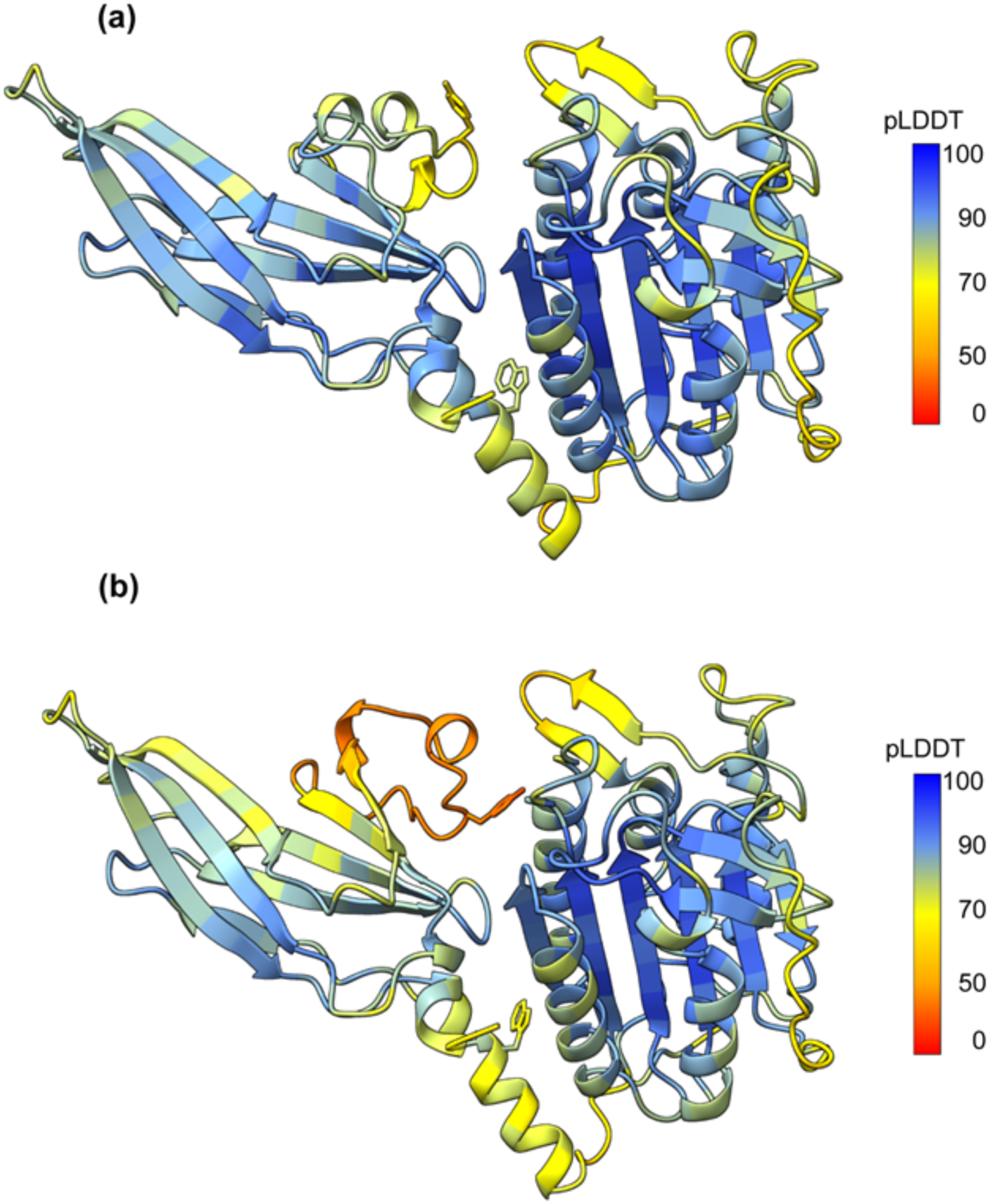
Molecular models of MALT1(PCASP-Ig3)_339–719_ predicted by **(a)** RoseTTAFold2 and **(b)** ESMFold2. Per-residue RoseTTAFold2 and ESMFold2 confidence scores (pLDDT) are mapped on the structures. The low pLDDT scores of 0-50 indicate confirmation heterogeneity.

**Figure S6.**
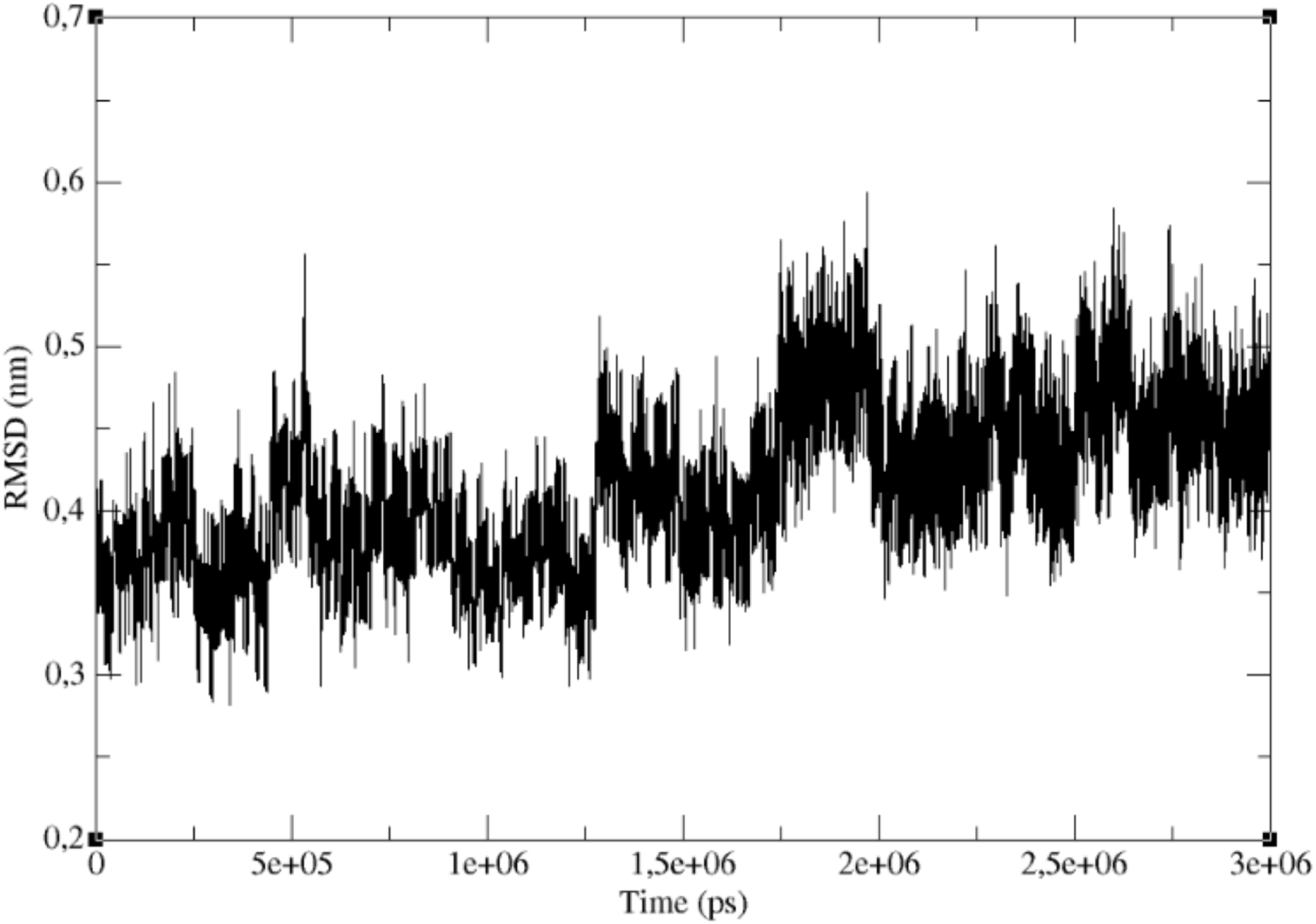
RMSD along the MD trajectory. RMSD trajectory obtained during 3us MD trajectory of MALT1(PCASP-Ig3)_339–719_. 2 us production run used for the analysis started after the first 1us equilibration. The starting structure was obtained from AF of MALT1(PCASP-Ig3)339–719 .

**Table S1.**
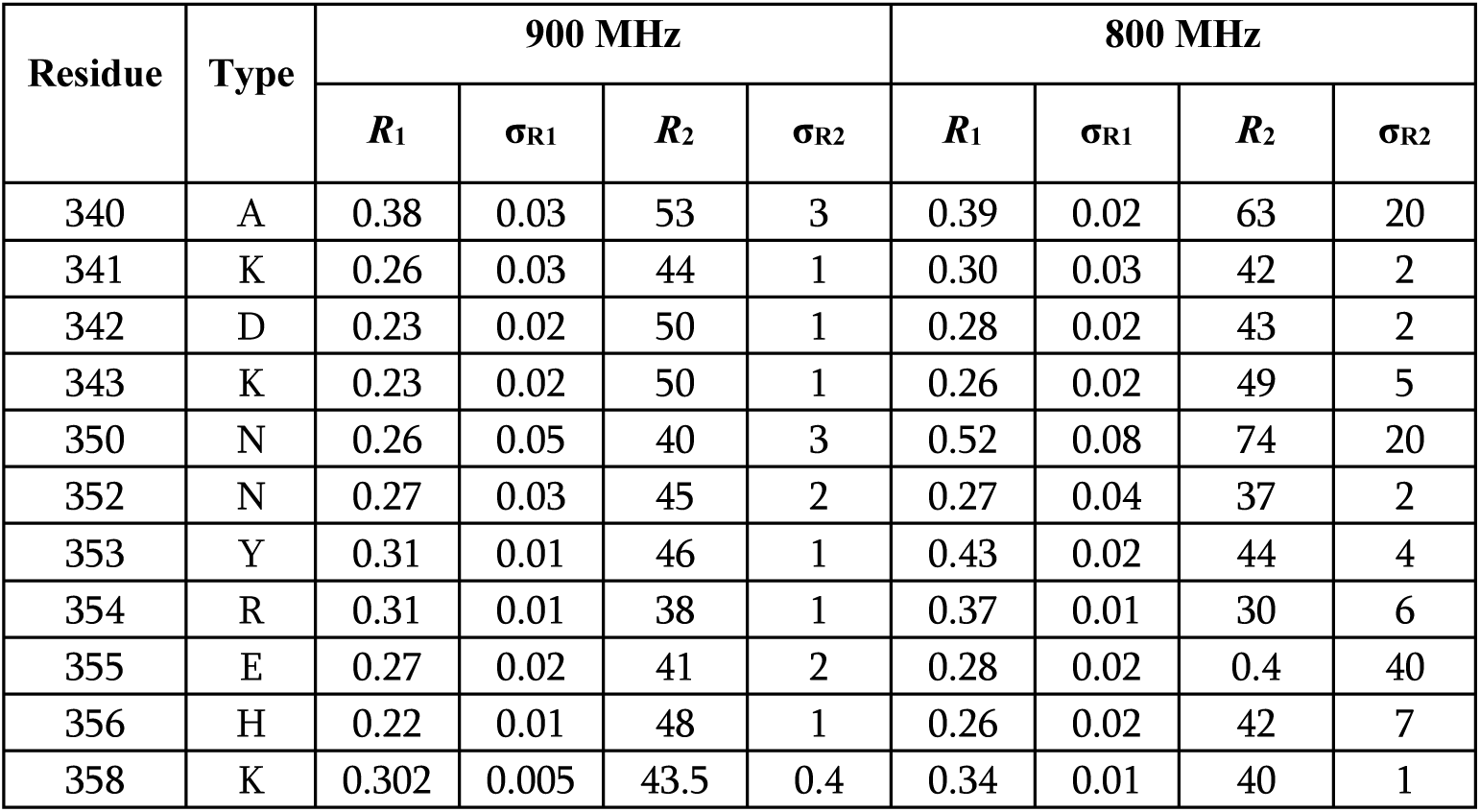
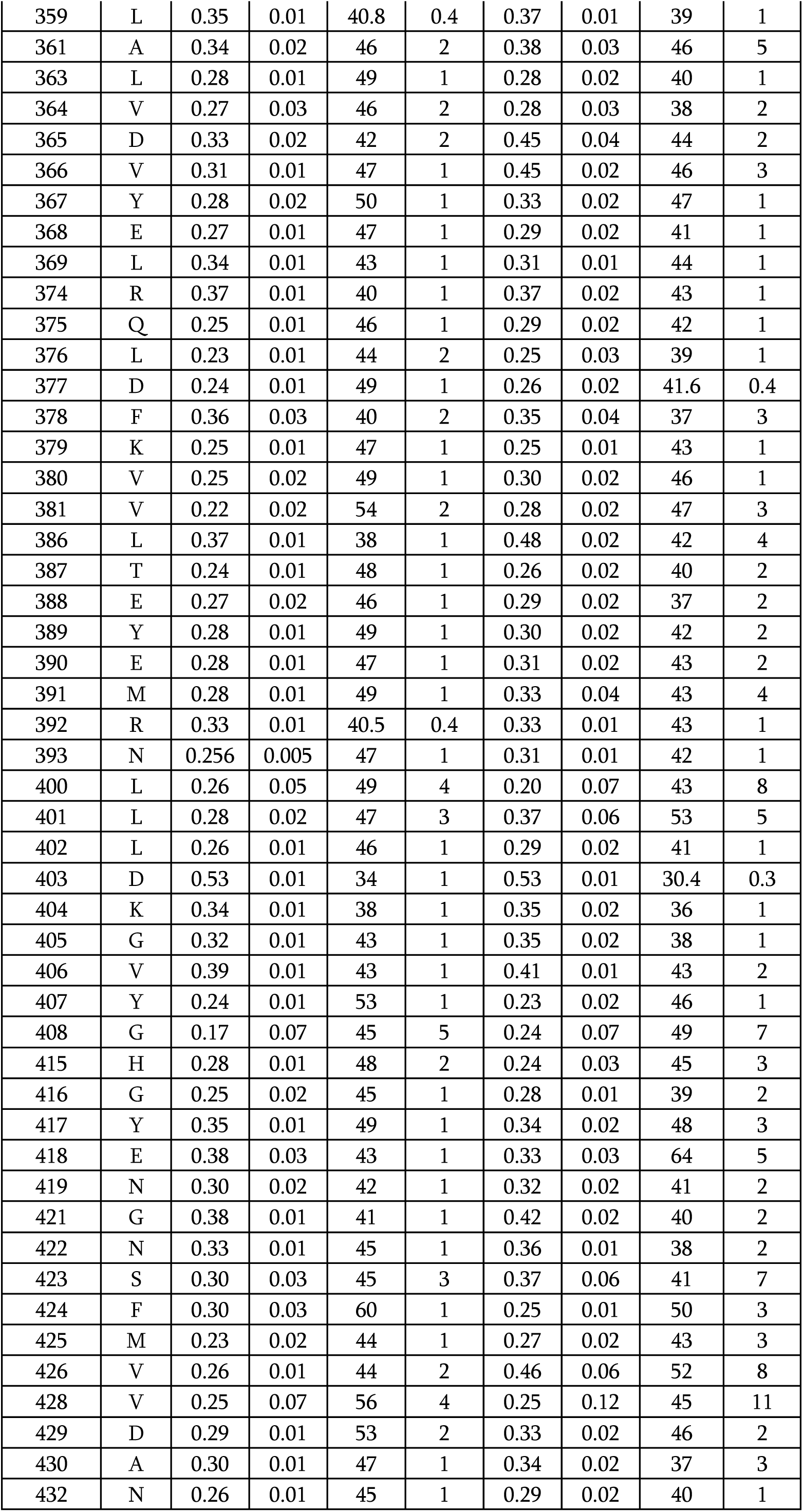
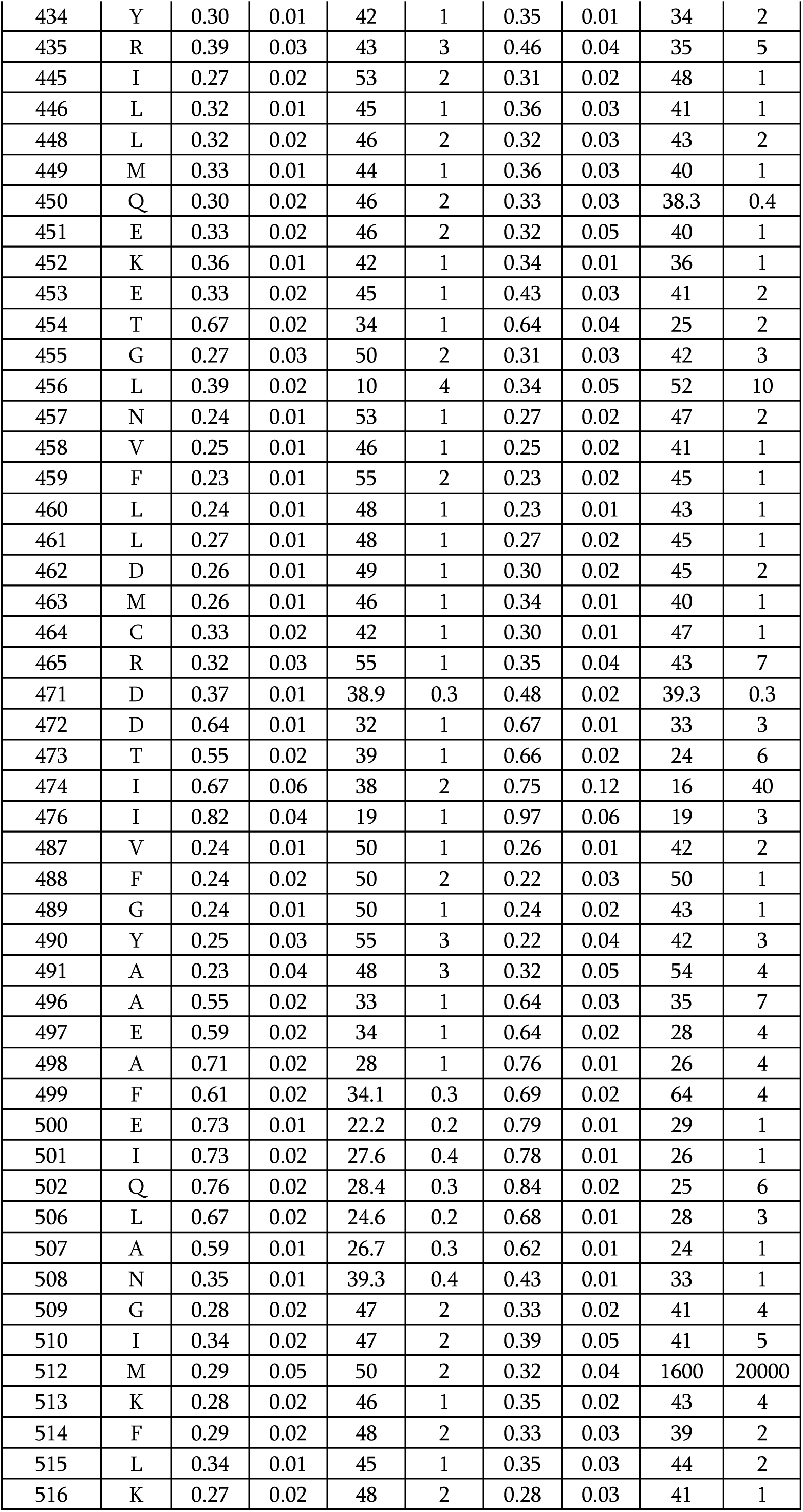
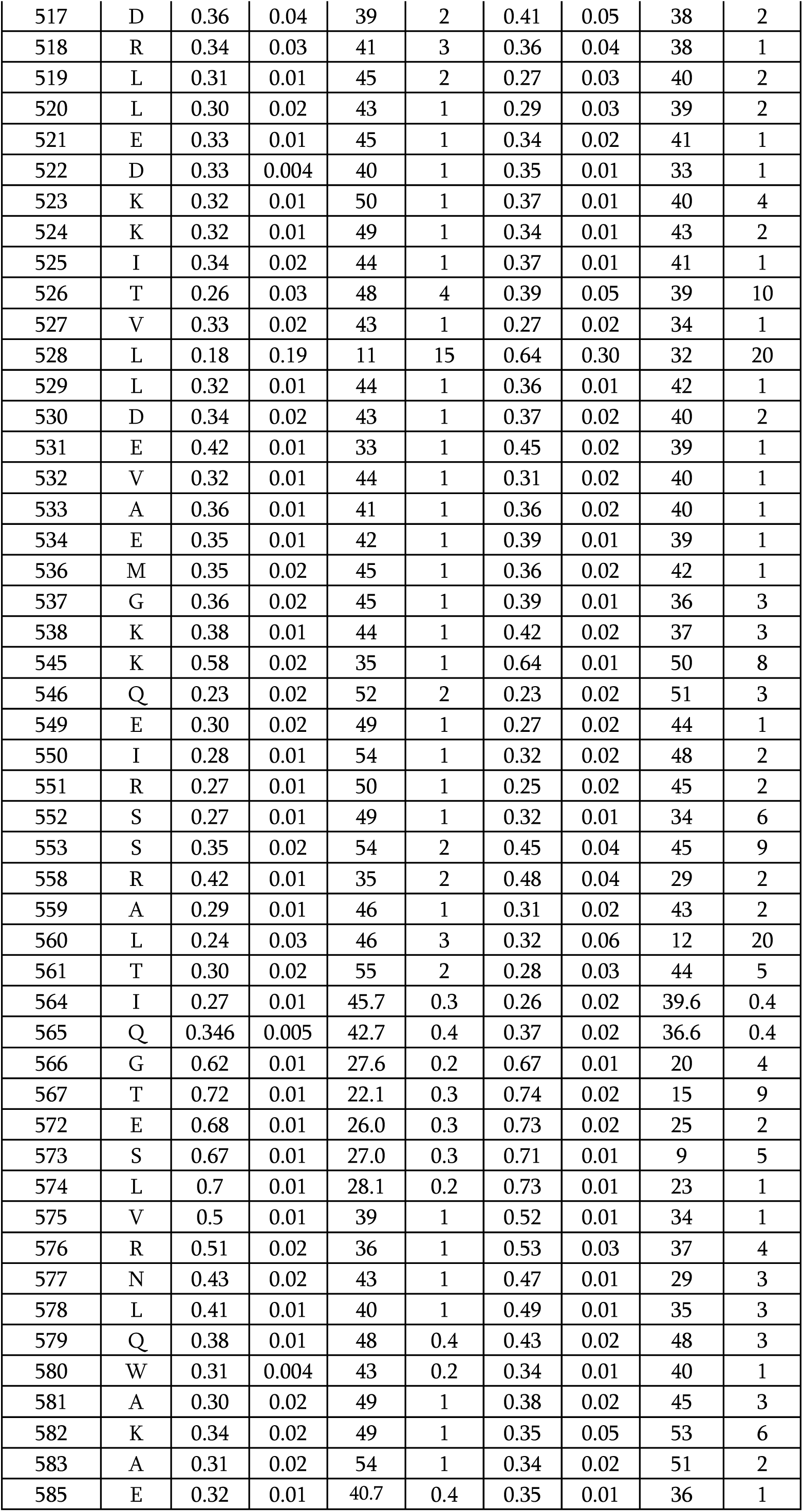
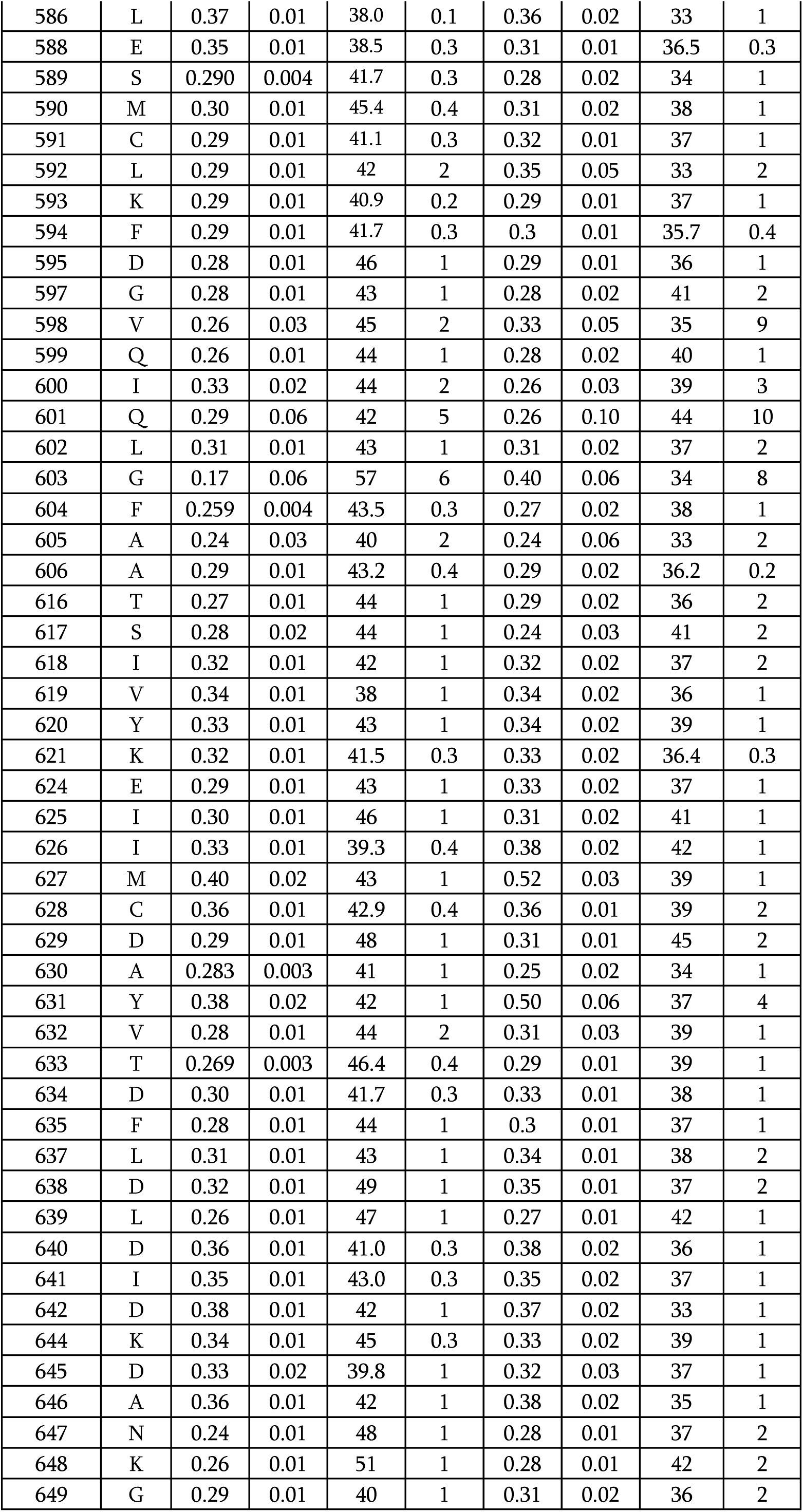
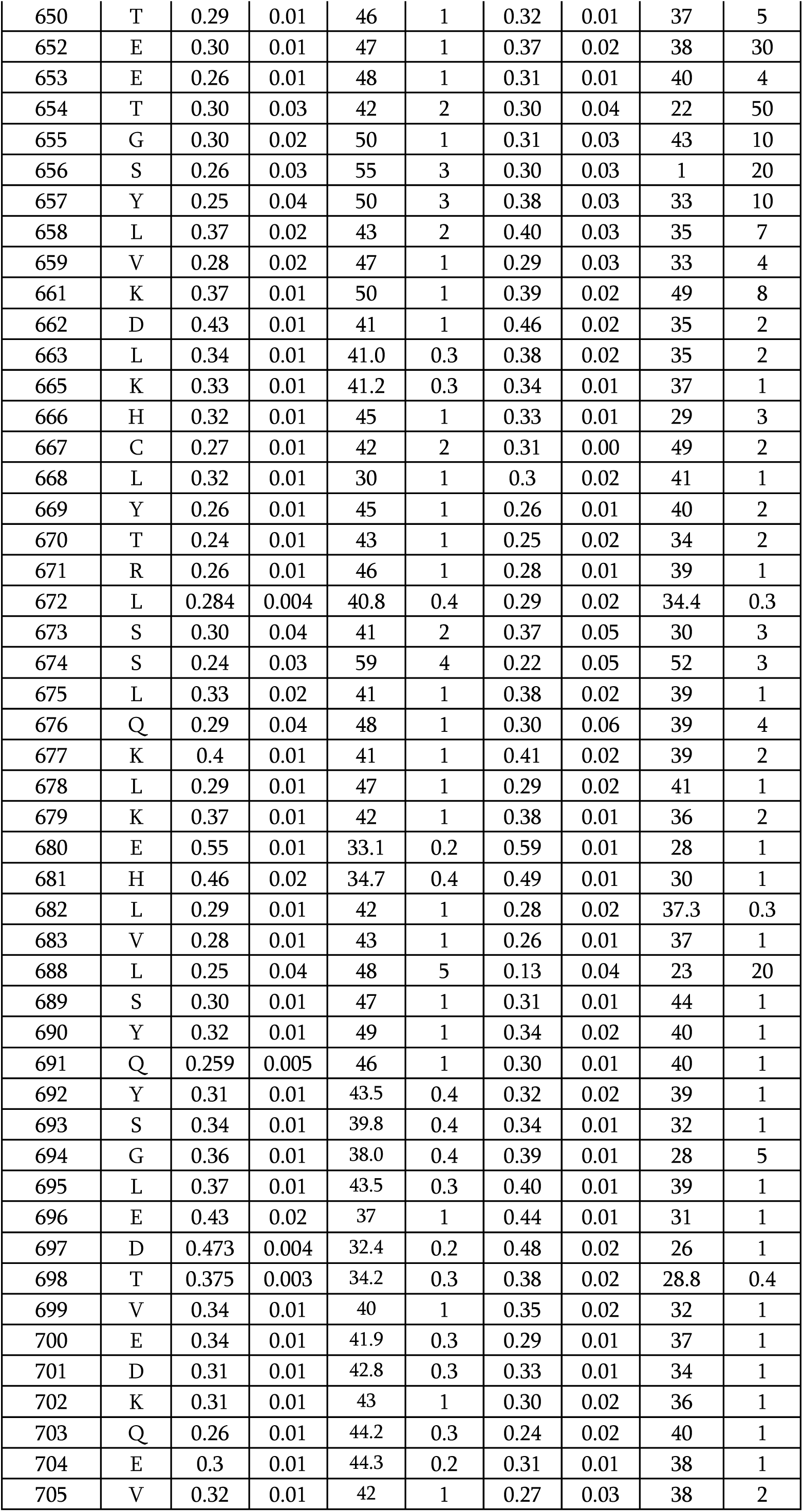
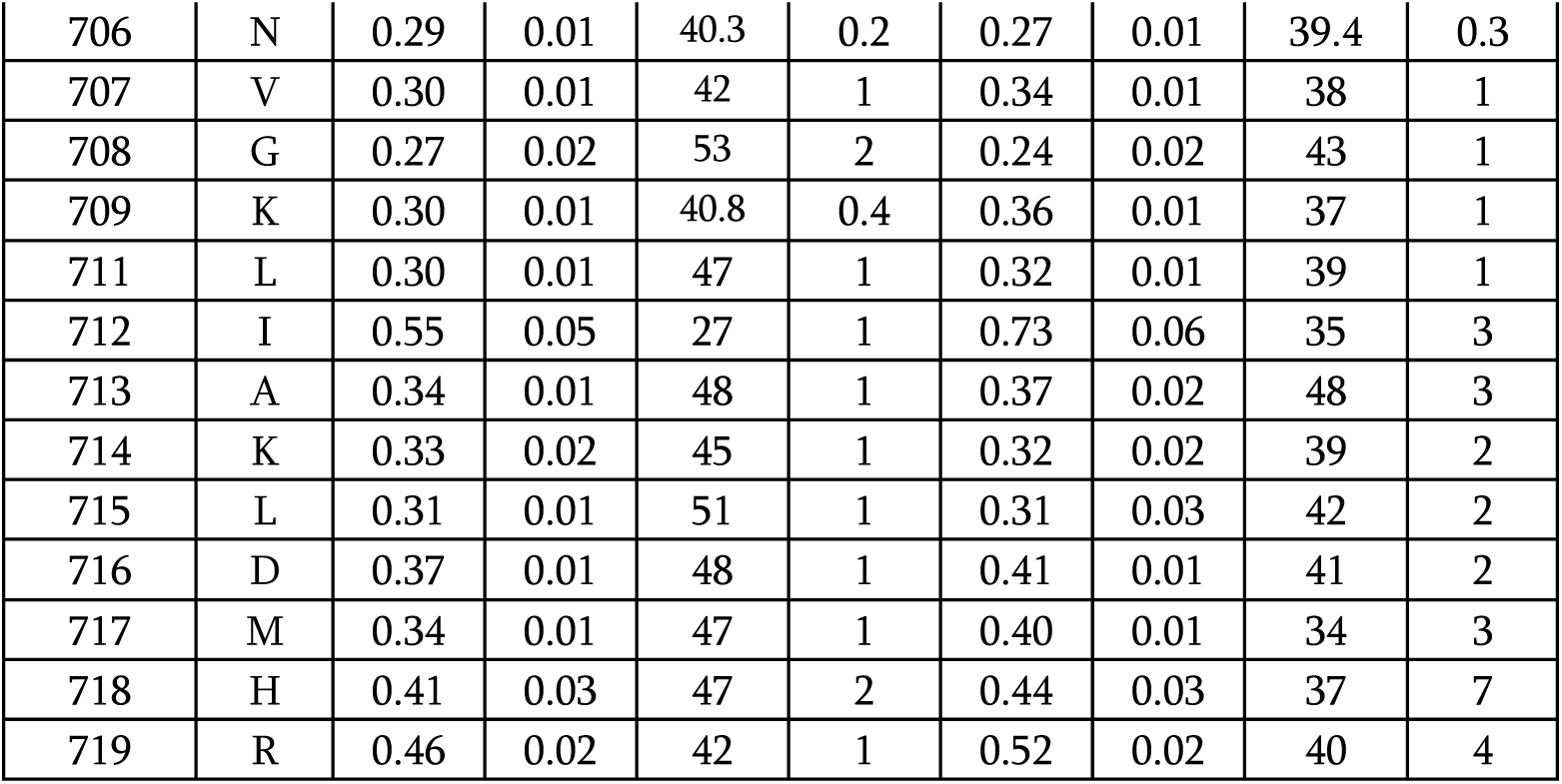
^15^N backbone relaxation rates *R*_1_, σ_R1_, *R*_2_, σ_R2_ for 900 MHz and 800 MHz.

**Table S2.** NOE ^1^H-^15^N relaxations for 800 MHz.

| Residue | Type | 800 MHz |  |
| --- | --- | --- | --- |
|  |  | NOE | error |
| 340 | A | 0.707 | 0.019 |
| 341 | K | 0.802 | 0.081 |
| 342 | D | 0.957 | 0.088 |
| 343 | K | 0.827 | 0.036 |
| 352 | N | 0.812 | 0.087 |
| 353 | Y | 0.722 | 0.022 |
| 354 | R | 0.764 | 0.040 |
| 355 | E | 1.027 | 0.156 |
| 356 | H | 0.742 | 0.053 |
| 358 | K | 0.727 | 0.024 |
| 359 | L | 0.694 | 0.029 |
| 361 | A | 0.713 | 0.131 |
| 363 | L | 0.782 | 0.062 |
| 364 | V | 0.663 | 0.098 |
| 365 | D | 0.853 | 0.092 |
| 366 | V | 0.717 | 0.035 |
| 367 | Y | 0.776 | 0.074 |
| 368 | E | 0.819 | 0.058 |
| 369 | L | 0.907 | 0.058 |
| 374 | R | 0.833 | 0.044 |
| 375 | Q | 0.941 | 0.065 |
| 376 | L | 0.783 | 0.066 |
| 377 | D | 0.815 | 0.047 |
| 378 | F | 0.555 | 0.102 |
| 379 | K | 0.776 | 0.039 |
| 380 | V | 0.646 | 0.054 |
| 381 | V | 0.768 | 0.089 |
| 386 | L | 0.821 | 0.061 |
| 387 | T | 0.824 | 0.048 |
| 388 | E | 0.773 | 0.057 |
| 389 | Y | 0.937 | 0.064 |
| 390 | E | 0.741 | 0.067 |
| 391 | M | 0.831 | 0.101 |
| 392 | R | 0.815 | 0.054 |
| 396 | D | 0.909 | 0.185 |
| 400 | L | 0.531 | 0.237 |
| 401 | L | 1.061 | 0.260 |
| 402 | L | 0.837 | 0.054 |
| 403 | D | 0.317 | 0.029 |
| 404 | K | 0.585 | 0.044 |
| 405 | G | 0.656 | 0.043 |
| 406 | V | 0.778 | 0.038 |
| 407 | Y | 0.745 | 0.061 |
| 414 | G | 0.570 | 0.121 |
| 415 | H | 0.874 | 0.101 |
| 416 | G | 0.845 | 0.068 |
| 417 | Y | 1.156 | 0.149 |
| 418 | E | 0.666 | 0.090 |
| 419 | N | 0.789 | 0.051 |
| 421 | G | 0.591 | 0.050 |
| 422 | N | 0.810 | 0.053 |
| 423 | S | 0.727 | 0.171 |
| 424 | F | 0.714 | 0.080 |
| 425 | M | 0.948 | 0.117 |
| 426 | V | 0.808 | 0.056 |
| 429 | D | 0.806 | 0.066 |
| 430 | A | 0.827 | 0.049 |
| 432 | N | 0.656 | 0.037 |
| 434 | Y | 0.677 | 0.035 |
| 435 | R | 0.539 | 0.124 |
| 444 | N | 0.799 | 0.029 |
| 445 | I | 1.072 | 0.092 |
| 446 | L | 0.899 | 0.079 |
| 448 | L | 0.637 | 0.055 |
| 449 | M | 0.865 | 0.087 |
| 450 | Q | 0.717 | 0.069 |
| 451 | E | 0.788 | 0.081 |
| 452 | K | 0.668 | 0.078 |
| 453 | E | 0.703 | 0.040 |
| 454 | T | 0.628 | 0.055 |
| 455 | G | 0.524 | 0.053 |
| 456 | L | 0.669 | 0.075 |
| 457 | N | 0.646 | 0.042 |
| 458 | V | 0.747 | 0.064 |
| 459 | F | 0.753 | 0.060 |
| 460 | L | 0.986 | 0.070 |
| 461 | L | 0.698 | 0.049 |
| 462 | D | 0.815 | 0.064 |
| 463 | M | 0.818 | 0.068 |
| 472 | D | 0.431 | 0.033 |
| 473 | T | 0.485 | 0.064 |
| 476 | I | 0.219 | 0.083 |
| 487 | V | 1.028 | 0.090 |
| 488 | F | 0.955 | 0.097 |
| 489 | G | 0.939 | 0.064 |
| 490 | Y | 0.846 | 0.154 |
| 491 | A | 0.821 | 0.123 |
| 495 | G | 0.478 | 0.079 |
| 496 | A | 0.525 | 0.052 |
| 497 | E | 0.490 | 0.045 |
| 498 | A | 0.466 | 0.034 |
| 499 | F | 0.502 | 0.031 |
| 500 | E | 0.426 | 0.016 |
| 501 | I | 0.445 | 0.028 |
| 502 | Q | 0.467 | 0.028 |
| 507 | A | 0.297 | 0.023 |
| 508 | N | 0.602 | 0.027 |
| 509 | G | 0.976 | 0.092 |
| 510 | I | 0.814 | 0.098 |
| 512 | M | 0.568 | 0.118 |
| 513 | K | 0.846 | 0.059 |
| 514 | F | 0.846 | 0.083 |
| 515 | L | 0.887 | 0.109 |
| 516 | K | 0.738 | 0.056 |
| 518 | R | 0.851 | 0.114 |
| 519 | L | 0.704 | 0.086 |
| 520 | L | 0.779 | 0.060 |
| 521 | E | 0.614 | 0.045 |
| 522 | D | 0.784 | 0.032 |
| 523 | K | 0.838 | 0.066 |
| 524 | K | 0.760 | 0.059 |
| 525 | I | 0.808 | 0.059 |
| 526 | T | 0.931 | 0.185 |
| 527 | V | 0.636 | 0.091 |
| 528 | L | 0.673 | 0.094 |
| 529 | L | 0.791 | 0.068 |
| 530 | D | 0.749 | 0.052 |
| 531 | E | 0.640 | 0.021 |
| 532 | V | 0.759 | 0.075 |
| 533 | A | 0.855 | 0.082 |
| 534 | E | 0.734 | 0.046 |
| 535 | D | 0.605 | 0.068 |
| 536 | M | 0.724 | 0.062 |
| 537 | G | 0.704 | 0.068 |
| 538 | K | 0.826 | 0.091 |
| 539 | C | 0.627 | 0.064 |
| 540 | H | 0.927 | 0.077 |
| 541 | L | 0.730 | 0.024 |
| 545 | K | 0.535 | 0.042 |
| 546 | Q | 0.931 | 0.108 |
| 549 | E | 0.712 | 0.073 |
| 550 | I | 0.650 | 0.062 |
| 551 | R | 0.862 | 0.062 |
| 552 | S | 0.765 | 0.054 |
| 558 | R | 0.292 | 0.087 |
| 559 | A | 0.937 | 0.063 |
| 560 | L | 0.842 | 0.179 |
| 561 | T | 0.830 | 0.101 |
| 562 | D | 0.774 | 0.049 |
| 564 | I | 0.671 | 0.031 |
| 565 | Q | 0.744 | 0.030 |
| 566 | G | 0.482 | 0.019 |
| 567 | T | 0.418 | 0.023 |
| 572 | E | 0.410 | 0.022 |
| 573 | S | 0.435 | 0.025 |
| 574 | L | 0.498 | 0.021 |
| 575 | V | 0.541 | 0.032 |
| 576 | R | 0.545 | 0.056 |
| 578 | L | 0.774 | 0.035 |
| 579 | Q | 0.694 | 0.054 |
| 580 | W | 0.740 | 0.025 |
| 581 | A | 0.715 | 0.059 |
| 582 | K | 0.813 | 0.119 |
| 583 | A | 0.900 | 0.079 |
| 585 | E | 0.717 | 0.024 |
| 586 | L | 0.704 | 0.027 |
| 588 | E | 0.791 | 0.045 |
| 589 | S | 0.556 | 0.023 |
| 590 | M | 0.728 | 0.037 |
| 591 | C | 0.613 | 0.029 |
| 592 | L | 0.791 | 0.018 |
| 593 | K | 0.639 | 0.026 |
| 594 | F | 0.685 | 0.028 |
| 595 | D | 0.835 | 0.045 |
| 597 | G | 0.870 | 0.039 |
| 598 | V | 0.772 | 0.197 |
| 599 | Q | 0.724 | 0.055 |
| 600 | I | 0.656 | 0.155 |
| 601 | Q | 0.338 | 0.246 |
| 602 | L | 0.708 | 0.039 |
| 603 | G | 0.547 | 0.207 |
| 604 | F | 0.780 | 0.028 |
| 605 | A | 0.519 | 0.160 |
| 606 | A | 0.811 | 0.032 |
| 616 | T | 0.793 | 0.049 |
| 617 | S | 0.747 | 0.069 |
| 618 | I | 0.648 | 0.027 |
| 619 | V | 0.735 | 0.050 |
| 620 | Y | 0.822 | 0.059 |
| 621 | K | 0.732 | 0.024 |
| 624 | E | 0.738 | 0.036 |
| 625 | I | 0.740 | 0.037 |
| 626 | I | 0.661 | 0.054 |
| 627 | M | 0.407 | 0.044 |
| 628 | C | 0.681 | 0.031 |
| 629 | D | 0.665 | 0.051 |
| 630 | A | 0.656 | 0.026 |
| 631 | Y | 0.764 | 0.066 |
| 632 | V | 0.755 | 0.073 |
| 633 | T | 0.805 | 0.031 |
| 634 | D | 0.757 | 0.033 |
| 635 | F | 0.650 | 0.031 |
| 637 | L | 0.760 | 0.036 |
| 638 | D | 0.631 | 0.052 |
| 639 | L | 0.735 | 0.053 |
| 640 | D | 0.568 | 0.027 |
| 641 | I | 0.610 | 0.025 |
| 642 | D | 0.445 | 0.021 |
| 644 | K | 0.664 | 0.031 |
| 645 | D | 0.661 | 0.041 |
| 646 | A | 0.605 | 0.034 |
| 647 | N | 0.812 | 0.062 |
| 648 | K | 0.840 | 0.045 |
| 649 | G | 0.762 | 0.042 |
| 650 | T | 0.923 | 0.065 |
| 652 | E | 0.812 | 0.070 |
| 653 | E | 0.693 | 0.068 |
| 654 | T | 0.661 | 0.102 |
| 655 | G | 0.720 | 0.082 |
| 656 | S | 0.503 | 0.096 |
| 657 | Y | 0.806 | 0.203 |
| 658 | L | 0.739 | 0.114 |
| 659 | V | 0.658 | 0.054 |
| 661 | K | 0.556 | 0.047 |
| 662 | D | 0.514 | 0.043 |
| 663 | L | 0.528 | 0.036 |
| 665 | K | 0.624 | 0.029 |
| 666 | H | 0.570 | 0.047 |
| 667 | C | 0.657 | 0.043 |
| 668 | L | 0.672 | 0.041 |
| 669 | Y | 0.845 | 0.054 |
| 670 | T | 0.738 | 0.041 |
| 671 | R | 0.830 | 0.038 |
| 672 | L | 0.790 | 0.025 |
| 673 | S | 0.589 | 0.111 |
| 674 | S | 0.658 | 0.109 |
| 675 | L | 0.833 | 0.054 |
| 676 | Q | 1.015 | 0.122 |
| 677 | K | 0.727 | 0.052 |
| 678 | L | 0.815 | 0.048 |
| 679 | K | 0.665 | 0.046 |
| 680 | E | 0.421 | 0.029 |
| 681 | H | 0.262 | 0.023 |
| 682 | L | 0.719 | 0.040 |
| 683 | V | 0.732 | 0.037 |
| 685 | T | 0.807 | 0.094 |
| 689 | S | 0.838 | 0.045 |
| 690 | Y | 0.755 | 0.036 |
| 691 | Q | 0.804 | 0.028 |
| 692 | Y | 0.509 | 0.028 |
| 693 | S | 0.556 | 0.026 |
| 694 | G | 0.531 | 0.033 |
| 695 | L | 0.478 | 0.028 |
| 696 | E | 0.180 | 0.020 |
| 697 | D | 0.004 | 0.017 |
| 698 | T | 0.285 | 0.017 |
| 699 | V | 0.609 | 0.027 |
| 700 | E | 0.607 | 0.020 |
| 701 | D | 0.694 | 0.020 |
| 702 | K | 0.644 | 0.025 |
| 703 | Q | 0.752 | 0.034 |
| 704 | E | 0.700 | 0.015 |
| 705 | V | 0.795 | 0.040 |
| 706 | N | 0.750 | 0.031 |
| 707 | V | 0.768 | 0.030 |
| 708 | G | 0.811 | 0.076 |
| 709 | K | 0.680 | 0.024 |
| 711 | L | 0.734 | 0.033 |
| 713 | A | 0.876 | 0.071 |
| 714 | K | 0.876 | 0.075 |
| 715 | L | 1.012 | 0.104 |
| 716 | D | 0.679 | 0.042 |
| 717 | M | 0.884 | 0.071 |
| 718 | H | 0.646 | 0.083 |
| 719 | R | 0.632 | 0.040 |

**Table S3.**
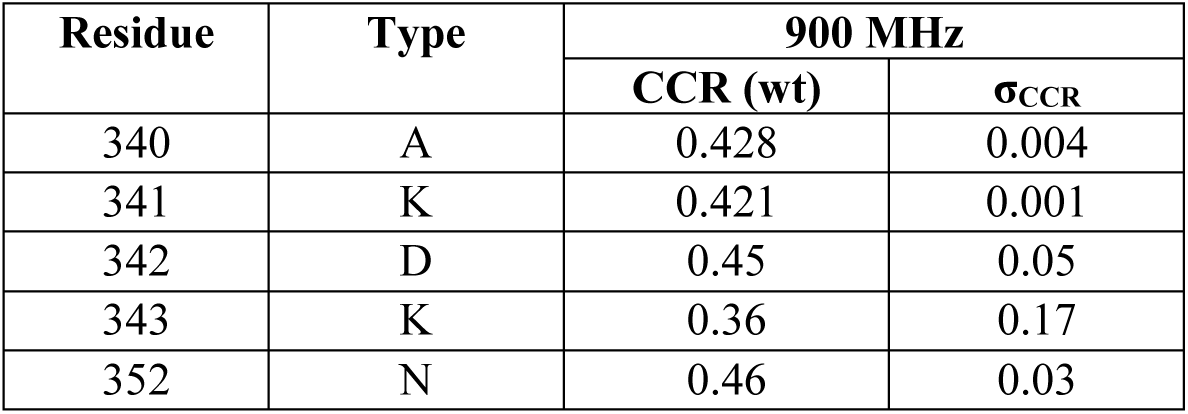
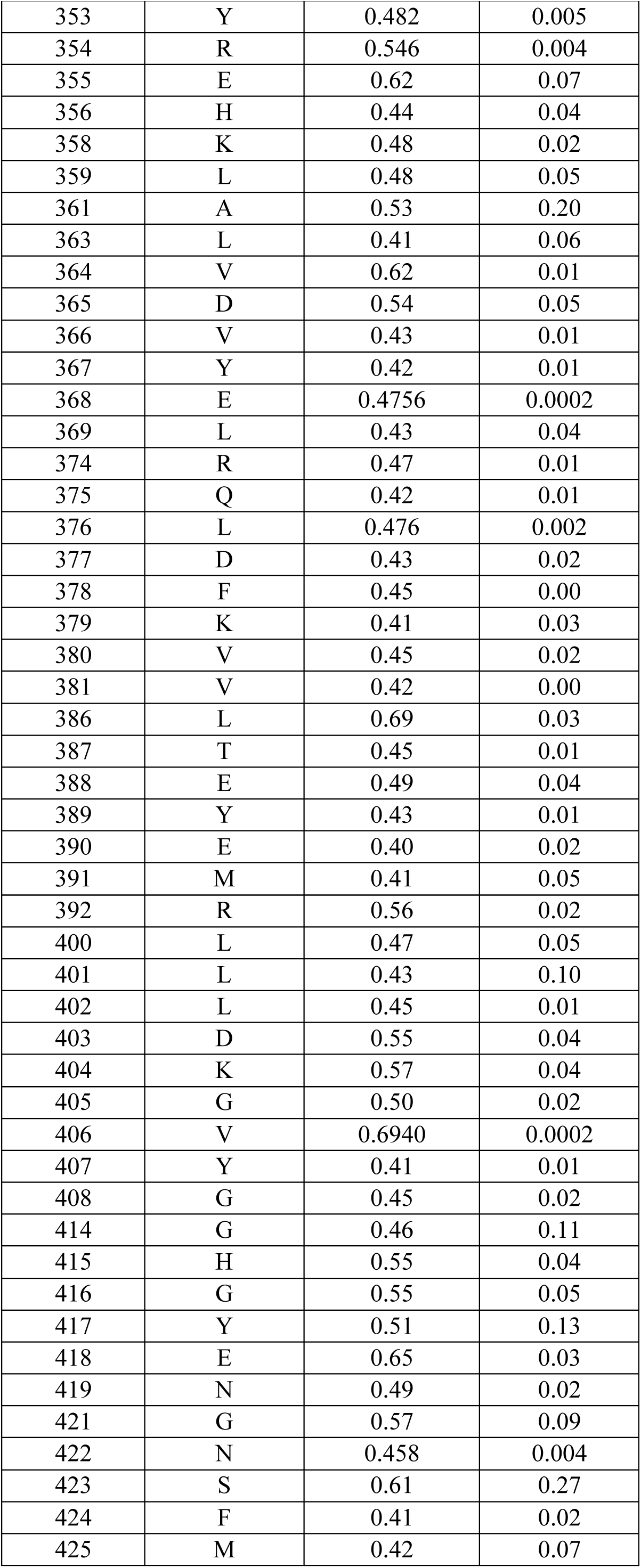
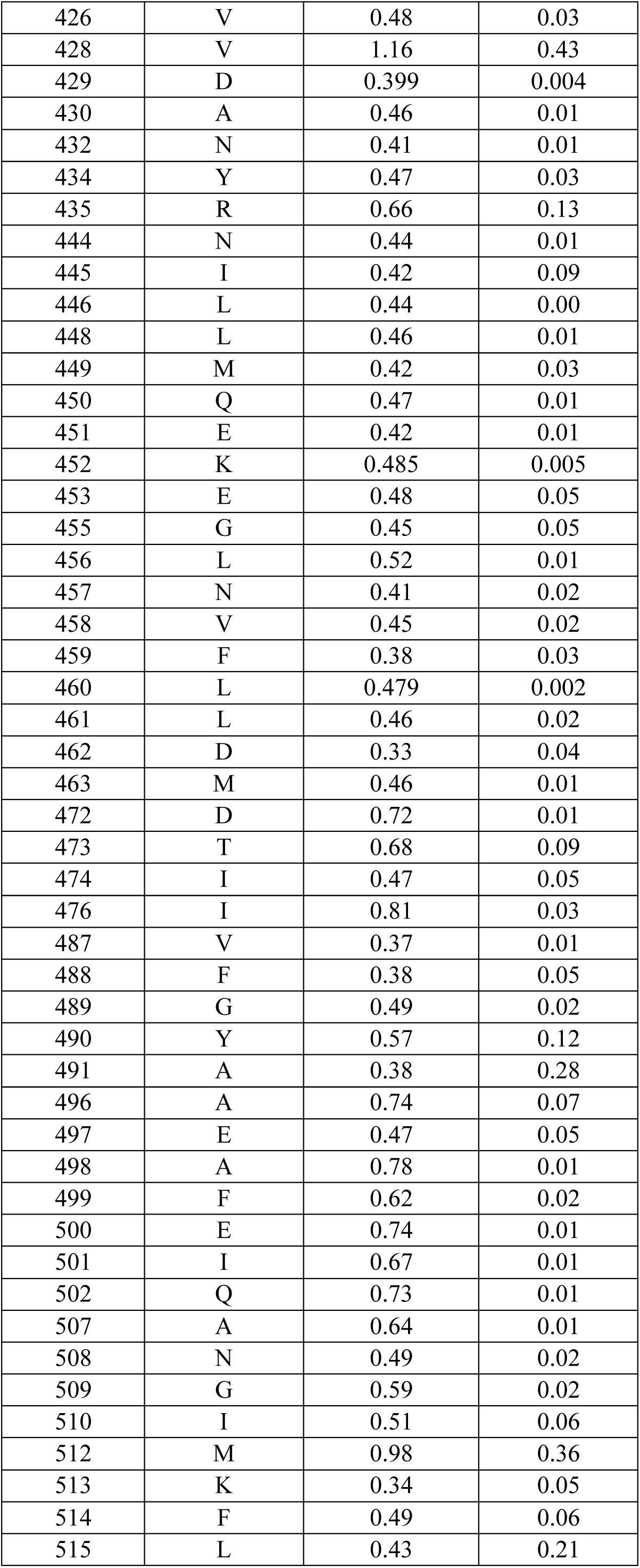
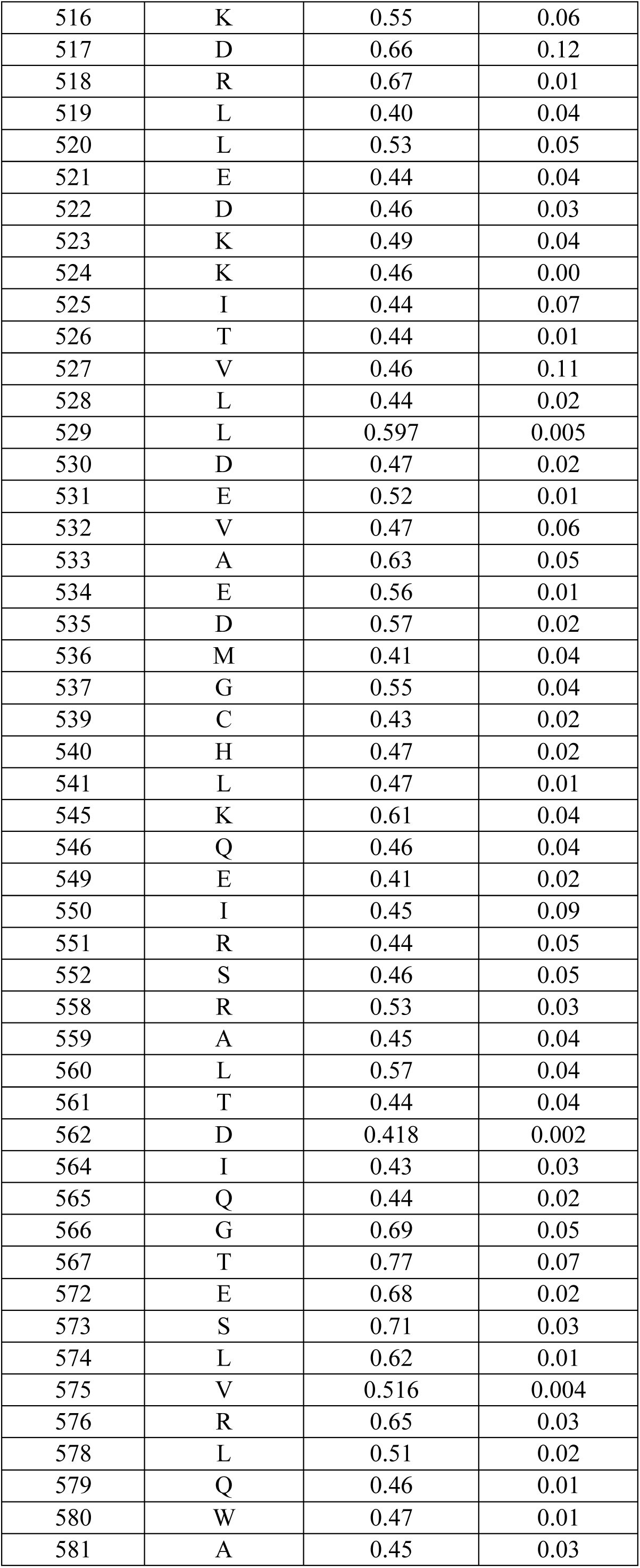
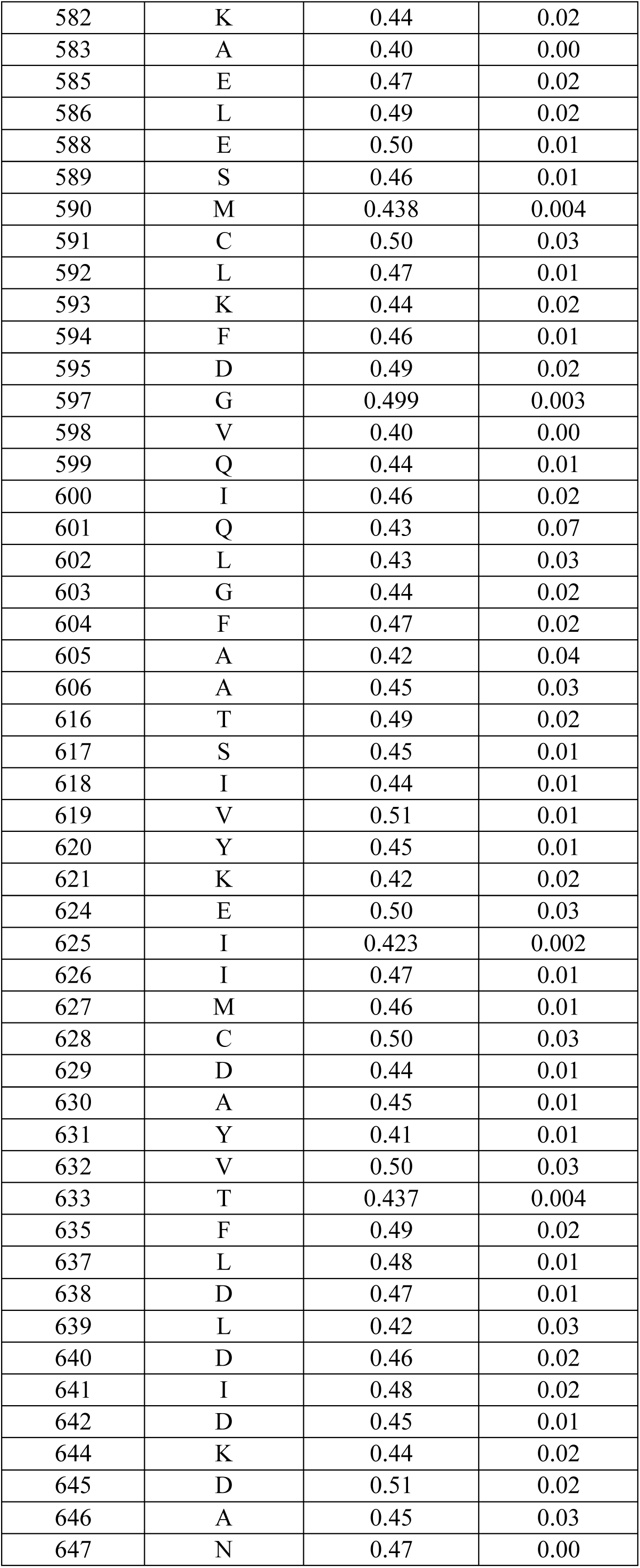
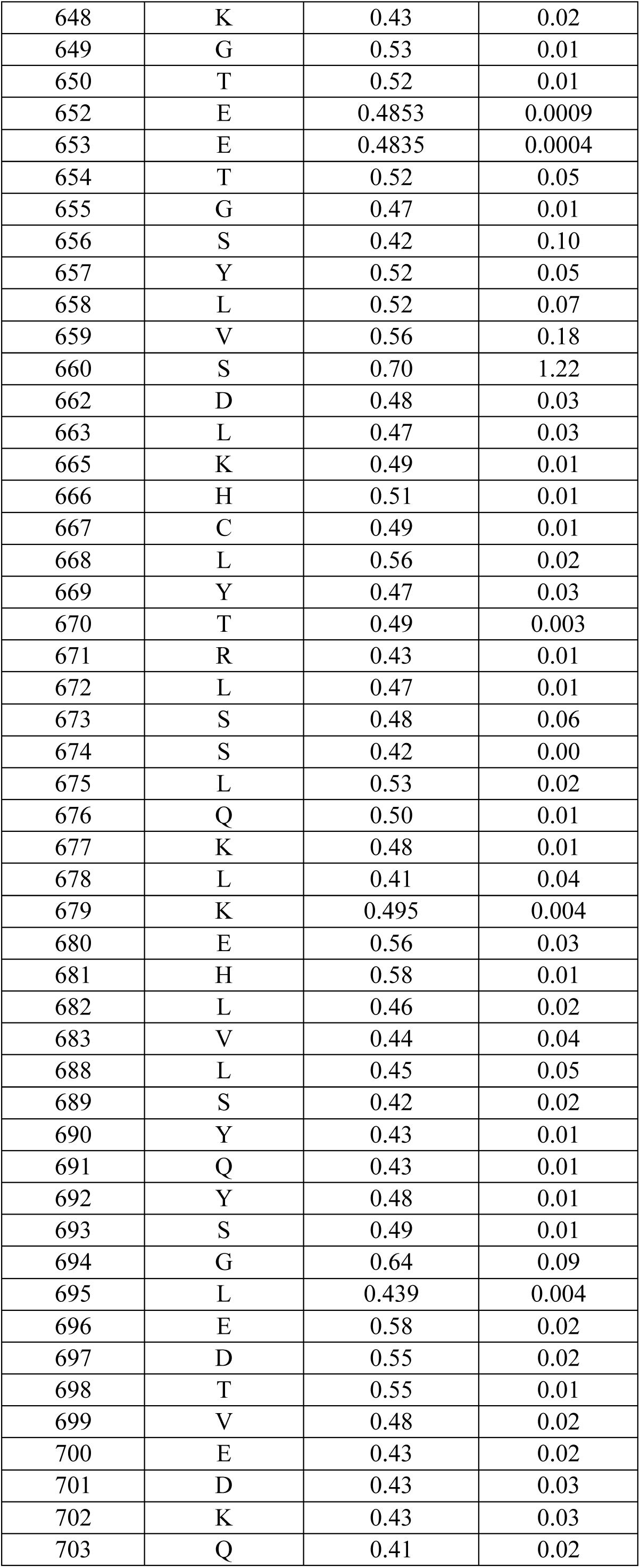
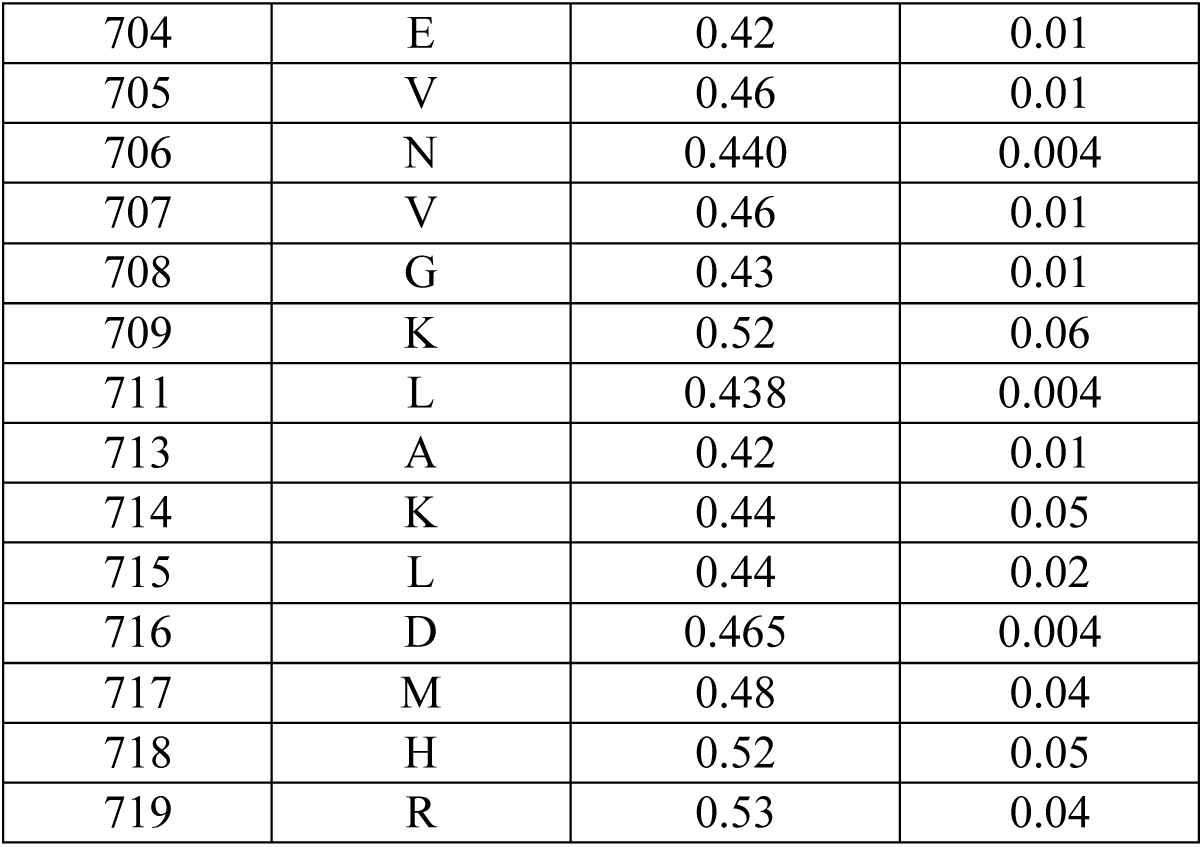
Cross correlation relaxation rates *CCR*, σ_CCR_, for 900.

| Residue | Type | 900 MHz |  |
| --- | --- | --- | --- |
| | | CCR (wt) | $\sigma_{CCR}$ |
| 340 | A | 0.428 | 0.004 |
| 341 | K | 0.421 | 0.001 |
| 342 | D | 0.45 | 0.05 |
| 343 | K | 0.36 | 0.17 |
| 352 | N | 0.46 | 0.03 |
| 353 | Y | 0.482 | 0.005 |
| 354 | R | 0.546 | 0.004 |
| 355 | E | 0.62 | 0.07 |
| 356 | H | 0.44 | 0.04 |
| 358 | K | 0.48 | 0.02 |
| 359 | L | 0.48 | 0.05 |
| 361 | A | 0.53 | 0.20 |
| 363 | L | 0.41 | 0.06 |
| 364 | V | 0.62 | 0.01 |
| 365 | D | 0.54 | 0.05 |
| 366 | V | 0.43 | 0.01 |
| 367 | Y | 0.42 | 0.01 |
| 368 | E | 0.4756 | 0.0002 |
| 369 | L | 0.43 | 0.04 |
| 374 | R | 0.47 | 0.01 |
| 375 | Q | 0.42 | 0.01 |
| 376 | L | 0.476 | 0.002 |
| 377 | D | 0.43 | 0.02 |
| 378 | F | 0.45 | 0.00 |
| 379 | K | 0.41 | 0.03 |
| 380 | V | 0.45 | 0.02 |
| 381 | V | 0.42 | 0.00 |
| 386 | L | 0.69 | 0.03 |
| 387 | T | 0.45 | 0.01 |
| 388 | E | 0.49 | 0.04 |
| 389 | Y | 0.43 | 0.01 |
| 390 | E | 0.40 | 0.02 |
| 391 | M | 0.41 | 0.05 |
| 392 | R | 0.56 | 0.02 |
| 400 | L | 0.47 | 0.05 |
| 401 | L | 0.43 | 0.10 |
| 402 | L | 0.45 | 0.01 |
| 403 | D | 0.55 | 0.04 |
| 404 | K | 0.57 | 0.04 |
| 405 | G | 0.50 | 0.02 |
| 406 | V | 0.6940 | 0.0002 |
| 407 | Y | 0.41 | 0.01 |
| 408 | G | 0.45 | 0.02 |
| 414 | G | 0.46 | 0.11 |
| 415 | H | 0.55 | 0.04 |
| 416 | G | 0.55 | 0.05 |
| 417 | Y | 0.51 | 0.13 |
| 418 | E | 0.65 | 0.03 |
| 419 | N | 0.49 | 0.02 |
| 421 | G | 0.57 | 0.09 |
| 422 | N | 0.458 | 0.004 |
| 423 | S | 0.61 | 0.27 |
| 424 | F | 0.41 | 0.02 |
| 425 | M | 0.42 | 0.07 |
| 426 | V | 0.48 | 0.03 |
| 428 | V | 1.16 | 0.43 |
| 429 | D | 0.399 | 0.004 |
| 430 | A | 0.46 | 0.01 |
| 432 | N | 0.41 | 0.01 |
| 434 | Y | 0.47 | 0.03 |
| 435 | R | 0.66 | 0.13 |
| 444 | N | 0.44 | 0.01 |
| 445 | I | 0.42 | 0.09 |
| 446 | L | 0.44 | 0.00 |
| 448 | L | 0.46 | 0.01 |
| 449 | M | 0.42 | 0.03 |
| 450 | Q | 0.47 | 0.01 |
| 451 | E | 0.42 | 0.01 |
| 452 | K | 0.485 | 0.005 |
| 453 | E | 0.48 | 0.05 |
| 455 | G | 0.45 | 0.05 |
| 456 | L | 0.52 | 0.01 |
| 457 | N | 0.41 | 0.02 |
| 458 | V | 0.45 | 0.02 |
| 459 | F | 0.38 | 0.03 |
| 460 | L | 0.479 | 0.002 |
| 461 | L | 0.46 | 0.02 |
| 462 | D | 0.33 | 0.04 |
| 463 | M | 0.46 | 0.01 |
| 472 | D | 0.72 | 0.01 |
| 473 | T | 0.68 | 0.09 |
| 474 | I | 0.47 | 0.05 |
| 476 | I | 0.81 | 0.03 |
| 487 | V | 0.37 | 0.01 |
| 488 | F | 0.38 | 0.05 |
| 489 | G | 0.49 | 0.02 |
| 490 | Y | 0.57 | 0.12 |
| 491 | A | 0.38 | 0.28 |
| 496 | A | 0.74 | 0.07 |
| 497 | E | 0.47 | 0.05 |
| 498 | A | 0.78 | 0.01 |
| 499 | F | 0.62 | 0.02 |
| 500 | E | 0.74 | 0.01 |
| 501 | I | 0.67 | 0.01 |
| 502 | Q | 0.73 | 0.01 |
| 507 | A | 0.64 | 0.01 |
| 508 | N | 0.49 | 0.02 |
| 509 | G | 0.59 | 0.02 |
| 510 | I | 0.51 | 0.06 |
| 512 | M | 0.98 | 0.36 |
| 513 | K | 0.34 | 0.05 |
| 514 | F | 0.49 | 0.06 |
| 515 | L | 0.43 | 0.21 |
| 516 | K | 0.55 | 0.06 |
| 517 | D | 0.66 | 0.12 |
| 518 | R | 0.67 | 0.01 |
| 519 | L | 0.40 | 0.04 |
| 520 | L | 0.53 | 0.05 |
| 521 | E | 0.44 | 0.04 |
| 522 | D | 0.46 | 0.03 |
| 523 | K | 0.49 | 0.04 |
| 524 | K | 0.46 | 0.00 |
| 525 | I | 0.44 | 0.07 |
| 526 | T | 0.44 | 0.01 |
| 527 | V | 0.46 | 0.11 |
| 528 | L | 0.44 | 0.02 |
| 529 | L | 0.597 | 0.005 |
| 530 | D | 0.47 | 0.02 |
| 531 | E | 0.52 | 0.01 |
| 532 | V | 0.47 | 0.06 |
| 533 | A | 0.63 | 0.05 |
| 534 | E | 0.56 | 0.01 |
| 535 | D | 0.57 | 0.02 |
| 536 | M | 0.41 | 0.04 |
| 537 | G | 0.55 | 0.04 |
| 539 | C | 0.43 | 0.02 |
| 540 | H | 0.47 | 0.02 |
| 541 | L | 0.47 | 0.01 |
| 545 | K | 0.61 | 0.04 |
| 546 | Q | 0.46 | 0.04 |
| 549 | E | 0.41 | 0.02 |
| 550 | I | 0.45 | 0.09 |
| 551 | R | 0.44 | 0.05 |
| 552 | S | 0.46 | 0.05 |
| 558 | R | 0.53 | 0.03 |
| 559 | A | 0.45 | 0.04 |
| 560 | L | 0.57 | 0.04 |
| 561 | T | 0.44 | 0.04 |
| 562 | D | 0.418 | 0.002 |
| 564 | I | 0.43 | 0.03 |
| 565 | Q | 0.44 | 0.02 |
| 566 | G | 0.69 | 0.05 |
| 567 | T | 0.77 | 0.07 |
| 572 | E | 0.68 | 0.02 |
| 573 | S | 0.71 | 0.03 |
| 574 | L | 0.62 | 0.01 |
| 575 | V | 0.516 | 0.004 |
| 576 | R | 0.65 | 0.03 |
| 578 | L | 0.51 | 0.02 |
| 579 | Q | 0.46 | 0.01 |
| 580 | W | 0.47 | 0.01 |
| 581 | A | 0.45 | 0.03 |
| 582 | K | 0.44 | 0.02 |
| 583 | A | 0.40 | 0.00 |
| 585 | E | 0.47 | 0.02 |
| 586 | L | 0.49 | 0.02 |
| 588 | E | 0.50 | 0.01 |
| 589 | S | 0.46 | 0.01 |
| 590 | M | 0.438 | 0.004 |
| 591 | C | 0.50 | 0.03 |
| 592 | L | 0.47 | 0.01 |
| 593 | K | 0.44 | 0.02 |
| 594 | F | 0.46 | 0.01 |
| 595 | D | 0.49 | 0.02 |
| 597 | G | 0.499 | 0.003 |
| 598 | V | 0.40 | 0.00 |
| 599 | Q | 0.44 | 0.01 |
| 600 | I | 0.46 | 0.02 |
| 601 | Q | 0.43 | 0.07 |
| 602 | L | 0.43 | 0.03 |
| 603 | G | 0.44 | 0.02 |
| 604 | F | 0.47 | 0.02 |
| 605 | A | 0.42 | 0.04 |
| 606 | A | 0.45 | 0.03 |
| 616 | T | 0.49 | 0.02 |
| 617 | S | 0.45 | 0.01 |
| 618 | I | 0.44 | 0.01 |
| 619 | V | 0.51 | 0.01 |
| 620 | Y | 0.45 | 0.01 |
| 621 | K | 0.42 | 0.02 |
| 624 | E | 0.50 | 0.03 |
| 625 | I | 0.423 | 0.002 |
| 626 | I | 0.47 | 0.01 |
| 627 | M | 0.46 | 0.01 |
| 628 | C | 0.50 | 0.03 |
| 629 | D | 0.44 | 0.01 |
| 630 | A | 0.45 | 0.01 |
| 631 | Y | 0.41 | 0.01 |
| 632 | V | 0.50 | 0.03 |
| 633 | T | 0.437 | 0.004 |
| 635 | F | 0.49 | 0.02 |
| 637 | L | 0.48 | 0.01 |
| 638 | D | 0.47 | 0.01 |
| 639 | L | 0.42 | 0.03 |
| 640 | D | 0.46 | 0.02 |
| 641 | I | 0.48 | 0.02 |
| 642 | D | 0.45 | 0.01 |
| 644 | K | 0.44 | 0.02 |
| 645 | D | 0.51 | 0.02 |
| 646 | A | 0.45 | 0.03 |
| 647 | N | 0.47 | 0.00 |
| 648 | K | 0.43 | 0.02 |
| 649 | G | 0.53 | 0.01 |
| 650 | T | 0.52 | 0.01 |
| 652 | E | 0.4853 | 0.0009 |
| 653 | E | 0.4835 | 0.0004 |
| 654 | T | 0.52 | 0.05 |
| 655 | G | 0.47 | 0.01 |
| 656 | S | 0.42 | 0.10 |
| 657 | Y | 0.52 | 0.05 |
| 658 | L | 0.52 | 0.07 |
| 659 | V | 0.56 | 0.18 |
| 660 | S | 0.70 | 1.22 |
| 662 | D | 0.48 | 0.03 |
| 663 | L | 0.47 | 0.03 |
| 665 | K | 0.49 | 0.01 |
| 666 | H | 0.51 | 0.01 |
| 667 | C | 0.49 | 0.01 |
| 668 | L | 0.56 | 0.02 |
| 669 | Y | 0.47 | 0.03 |
| 670 | T | 0.49 | 0.003 |
| 671 | R | 0.43 | 0.01 |
| 672 | L | 0.47 | 0.01 |
| 673 | S | 0.48 | 0.06 |
| 674 | S | 0.42 | 0.00 |
| 675 | L | 0.53 | 0.02 |
| 676 | Q | 0.50 | 0.01 |
| 677 | K | 0.48 | 0.01 |
| 678 | L | 0.41 | 0.04 |
| 679 | K | 0.495 | 0.004 |
| 680 | E | 0.56 | 0.03 |
| 681 | H | 0.58 | 0.01 |
| 682 | L | 0.46 | 0.02 |
| 683 | V | 0.44 | 0.04 |
| 688 | L | 0.45 | 0.05 |
| 689 | S | 0.42 | 0.02 |
| 690 | Y | 0.43 | 0.01 |
| 691 | Q | 0.43 | 0.01 |
| 692 | Y | 0.48 | 0.01 |
| 693 | S | 0.49 | 0.01 |
| 694 | G | 0.64 | 0.09 |
| 695 | L | 0.439 | 0.004 |
| 696 | E | 0.58 | 0.02 |
| 697 | D | 0.55 | 0.02 |
| 698 | T | 0.55 | 0.01 |
| 699 | V | 0.48 | 0.02 |
| 700 | E | 0.43 | 0.02 |
| 701 | D | 0.43 | 0.03 |
| 702 | K | 0.43 | 0.03 |
| 703 | Q | 0.41 | 0.02 |
| 704 | E | 0.42 | 0.01 |
| 705 | V | 0.46 | 0.01 |
| 706 | N | 0.440 | 0.004 |
| 707 | V | 0.46 | 0.01 |
| 708 | G | 0.43 | 0.01 |
| 709 | K | 0.52 | 0.06 |
| 711 | L | 0.438 | 0.004 |
| 713 | A | 0.42 | 0.01 |
| 714 | K | 0.44 | 0.05 |
| 715 | L | 0.44 | 0.02 |
| 716 | D | 0.465 | 0.004 |
| 717 | M | 0.48 | 0.04 |
| 718 | H | 0.52 | 0.05 |
| 719 | R | 0.53 | 0.04 |

